# Human ectodermal organoids reveal the cellular origin of DiGeorge Syndrome

**DOI:** 10.1101/2025.08.08.669417

**Authors:** Ed Zandro M. Taroc, Surangi Perera, Tunde Berecz, Jenny Hsin, Karla Barbosa-Sabanero, Sravya Pailla, Zarin Zainul, Ceren Pajanoja, Agota Apati, Daniel Martin, Laura Kerosuo

**Affiliations:** Neural Crest Development and Disease Unit, National Institute of Dental and Craniofacial Research, Intramural Research Program, National Institutes of Health, Bethesda, USA; Department of Biochemistry and Developmental Biology, Biomedicum, University of Helsinki, Finland; Institute of Molecular Life Sciences, HUN-REN RCNS, Budapest, Hungary; National Institute of Dental and Craniofacial Research, Intramural Research Program, Genomics and Computational Biology Core, National Institutes of Health, Bethesda, USA

## Abstract

Neurocristopathies account for half of all birth defects and several cancers highlighting the need to understand early neural crest (NC) development, for which suitable human models don’t exist. Here, we present a pluripotent-stem-cell-based 3D ectodermal organoid model that faithfully recapitulates early ectodermal patterning of future central nervous system, epidermis and cranial and trunk NC, as well as a diverse selection of NC derivatives —offering a comprehensive platform to study neurocristopathies from early induction of pluripotent-like stem cells at the neural plate border to differentiated cells. DiGeorge syndrome (DGS) is caused by a hemizygous microdeletion of ∼fifty genes, many of which play broad roles during embryogenesis. While DGS is traditionally considered to originate from all germ layers, its clinical manifestations—including craniofacial anomalies, cardiac outflow tract defects, thymic hypoplasia and thyroid dysfunction—are consistent with tissues requiring NC contributions. This raises the question why DGS mainly manifest in organs of NC origin? Using patient-derived iPSCs, we show that DGS organoids display reduced pluripotency gene expression and impaired maintenance of ectodermal stem cells, leading to defective NC specification, posteriorization of cranial NC, and failure to form cartilage. We identify a small subset of genes within the DGS deletion as potential drivers of these early NC defects. Consequently, impaired NC cells are further affected during cranial and vagal mesenchyme maturation, likely worsened by hemizygosity of additional DGS genes that individually, without the initial NC defect, are insufficient to cause the disease. We hypothesize that DGS is primarily, or possibly entirely, a neurocristopathy.

## Introduction

Millions of people are diagnosed with neurocristopathies yearly, primarily with defects in the craniofacial skeleton, cardiac outflow track and the gut; with conditions commonly requiring surgical intervention and life-long medical care ^1^. Furthermore, a significant proportion of cancers worldwide originate from neural crest cells including melanoma (∼ 300K diagnosed cases yearly) as well as Pheochomocytoma & paraganglioma as well as the pediatric lethal cancer neuroblastoma (∼ 5K diagnosed cases yearly of each). To date, there is no model to study early induced neurocristopathies or other defects that occur during the ectodermal patterning process.

The neural crest is an embryonic stem cell population unique to vertebrates. It rises from the ectodermal germ layer right after gastrulation with induction at the neural plate border. During neurulation, as the neural plate closes to form the neural tube, the neural crest is specified in the dorsal part of the neural folds, undergoes EMT and delaminates from the neural epithelium. Migratory neural crest cells follow stereotypical pathways to diverse destinations within the growing embryo, where they will give rise to distinct derivatives that range from typical ectodermal progeny, including the peripheral nervous system and melanocytes, to mesodermal-like cell types including the facial bone and cartilage, and various other mesenchymal cell types like pericytes, providing e.g. the forebrain with functional vasculature and meninges, adipocytes and connective tissue in the head and neck region such as in the thymus, thyroid, parathyroid, cornea. The neural crest also forms the cardiac septa and the smooth muscle lining of the aortic arch arteries in the outflow track, as well as “endoderm-like” neuroendocrine cells like the chromaffin cells of the adrenal medulla, and glomus cells of the carotid body that release hormones to the blood stream ^2–5^, as well as contribution to exocrine cells in the lacrimal gland and the acinar cells of the salivary glands ^6^. This astonishing pleistopotency, a category between pluripotent and multipotent stem cell potential that extends beyond the germ layer rule ^4^, is enabled by the ectodermal stem cells that maintain expression of pluripotency genes from blastula stages, which co-express neural crest and other lineage markers during ectoderm patterning ^7,8^. Importantly, the array of neural crest derivatives varies between different body axes, and in the commonly used vertebrate models, the mesodermal-like cell types are unique to the cranial and vagal regions ^4^.

In human embryos, gastrulation ends, and the ectoderm starts to pattern at the third week of gestation ^9,10^. Due to lack of availability of early human embryos caused by ethical and practical reasons, there are no in vivo studies on the premigratory stages of human cranial neural crest, and only a few descriptive reports on the later-forming trunk axial levels have been published ^11,12^, whereas data on the differentiation pathways of migratory neural crest derivatives from older embryos is starting to emerge ^13^. Consequently, knowledge of neural crest formation within the ectoderm largely stems from studies in chick, mouse, frog, and zebrafish embryos. While neural crest development is broadly conserved across model species, differences exist in gene sets, activation order, paralog usage, compensation mechanisms, and the timing of migration relative to neural tube closure ^14,15^. Importantly, neurocristopathies comprise at least half of all birth defects (Torres et al, manuscript will be available for reference in a few months), which include craniofacial malformations like lip and/or cleft palate, craniofrontonasal syndrome, coronal craniosynostosis, maxillary hypoplasia and the ocular defects causing Axenfeld -Rieger syndrome ^16,17^; cardiac septum and aortic arch artery defects, the intestinal agangliosis known as Hirsprung disease ^18^, and the melanocyte deficiency piebaldism. Several syndromes including DiGeorge, CHARGE, Treacher-Collins, and Waardenburg, to name a few ^9,19^, present symptoms in multiple neural crest derived organs, suggesting the disease onset took place before lineage diversication. Similarly, the malignancy of many cancers is thought to result from reversion to transcriptional states typical of their progenitor populations. Understanding human ectodermal patterning and neural crest induction, therefore, is critical to elucidating the etiology of these widespread and frequently life-threatening disorders. Moreover, as stem cell-based regenerative therapies progress, precise protocols for generating neural crest-derived cell types for replacement therapy are increasingly needed.

Multiple protocols for deriving neural crest cells from human pluripotent stem cells have been developed over the past 15 years. However, transcriptional characterization and purity assessments of these cultures remain superficial, and the early developmental stages of the developing ectodermal epithelium prior to migration have not been addressed ^20–32^.Yet, comprehensive understanding of the molecular events that take place on the dish is a prerequisite for successful disease modeling and interpretation of results.

DiGeorge syndrome (DGS, *aka* the 22q11.2 deletion syndrome; velocardiofacial syndrome) is a relatively common disease that manifests in approximately one in 3000 to one in 4000 live births. Main symptoms include prevalent conotruncal cardiac and septum anomalies, craniofacial defects, thymic and parathyroid hypoplasia, and gastrointestinal and renal abnormalities ^33–36^. Additionally, the symptoms, which vary between patients, include learning disabilities, delayed growth and a higher risk of schizophrenia, ADHD and autism spectrum disorders. Due to the hypocalcemia, immunological, cardiac and neurological defects, DiGeorge syndrome is traditionally not fully acknowledged as a neurocristopathy. However, growing evidence suggest that the affected organs can be impaired due to deficits in neural crest development hence it forms the craniofacial complex (bone, cartilage, sutures), mesenchymal counterparts of thymus and parathyroid as well as pericytes in the meninges of forebrain in addition to the septum and outflow track of the heart that are affected in the patients. ^2,4^. Interestingly, a recent study found a connection between abnormal skull formation and defects in neuropsychciatric behavior, further building the connection of neurological symptoms caused by cranial skeleton anomalies ^37^ Most cases of the typical ∼3MB microdeletion result from a *de novo* mutation ^35^ while inheritance is estimated to occur in 6–28% of patients ^36^. However, the complexity of the disease-causing molecular mechanisms that lead to the phenotypes is poorly understood and the effects of the microdeletion to early neural crest development are not known.

## Results

### Image analysis shows that ectodermal organoids consist of neural, neural crest and non-neural ectoderm domains and migratory cells are a pure population of neural crest

Our aim was to obtain an *in vitro* model that would faithfully recapitulate early embryonic ectoderm development during third and fourth week of human gestation. None of the existing NC differentiation protocols have addressed the developmental *in vitro* steps of the respective protocols, as the rudimentary characterization has only been performed on the mature, migratory NC cells. Additionally, the majority of the established protocols grow as 2D monolayers and do not provide a basis for evaluating ectodermal patterning which occurs via crosstalk between domains. However, in the Bajpai et al, 2010 – protocol cells cluster to form spheroids from which neural crest cells migrate out ^20^. We optimized the protocol by modifying the culture medium by doubling the previous concentration of the EGF, FGF and insulin growth factors, from now on referred to as ectodermal organoid culture medium. We analyzed the spheroids in these cultures asking whether they recapitulate the whole process of ectodermal patterning, similar to the embryo, or do they immediately obtain a neural crest oriented developmental trajectory? In short, after resuspending embryonic stem cell colonies into small aggregates (of 30-50 cells), they were placed on regular cell culture dishes in the ectodermal organoid medium. By day two, small floating spheroids were formed. On day five to six, first organoids started to adhere to the bottom of the tissue culture dish and first migratory neural crest cells are visibly delaminating out of the spheroids. By day ten, large migratory mats of neural crest cells were formed as halos around the attached organoids, although some remain floating (Fig. 1 A-E). As expected, immunofluorescence labeling of the neural crest specification marker SOX9 and the non-neural ectoderm/neural plate border/neural crest marker TFAP2A revealed high overlapping domains in the floating organoids at day 6 (Fig. 1F), but some TFAP2A positive cells in the spheroids were negative for SOX9 (arrows) (Fig. 1F^1–2^) indicating these cells were at an earlier stage of neural crest development or specified into non-neural ectoderm. Notably, based on immunoreactivity, the organoids didn’t only consist of the developing neural crest cells. As the neural plate and neural tube that will form the central nervous system are highly positive for SOX2 and PAX6 expressing neural stem cells ^38^, we tested whether the organoids were positive for SOX2 and PAX6, which indeed was the case (Fig. 1 F^3^,H). Finally, we used HCR-fluorescent *in situ* hybridization to show that the spheroids also contained domains of *DLX5*-positive future epidermal cells (Fig. 1G), suggesting that the spheroids are organoids that contain all ectodermal domains. Next, we assessed the purity of the migratory neural crest population by staining with TFAP2A and SOX9 at Day 9. Quantification of the immunopositive cells for SOX9 and TFAP2A, respectively, in the migratory mats revealed that all the cells expressed at least one of the two neural crest markers and their overlap was over 90% (Fig. 1J-J^3^). Notably, many of the cells in the migratory mats also expressed the migratory NC marker TWIST1 (Fig. 1I). Interestingly, while virtually all early and late migrating cells were positive for TFAP2A, another neural crest specifier protein, FOXD3, was expressed only in the newly migrating cells (arrowheads) (Fig. 1K), recapitulating its expression pattern in the embryo ^39,40^. Both TFAP2A and FOXD3 continued to be expressed in the spheroids indicating that production of premigratory neural crest cells remained while the first migratory cells had already emigrated the organoids (Fig. 1 K-K^3^).

**Figure 1.**
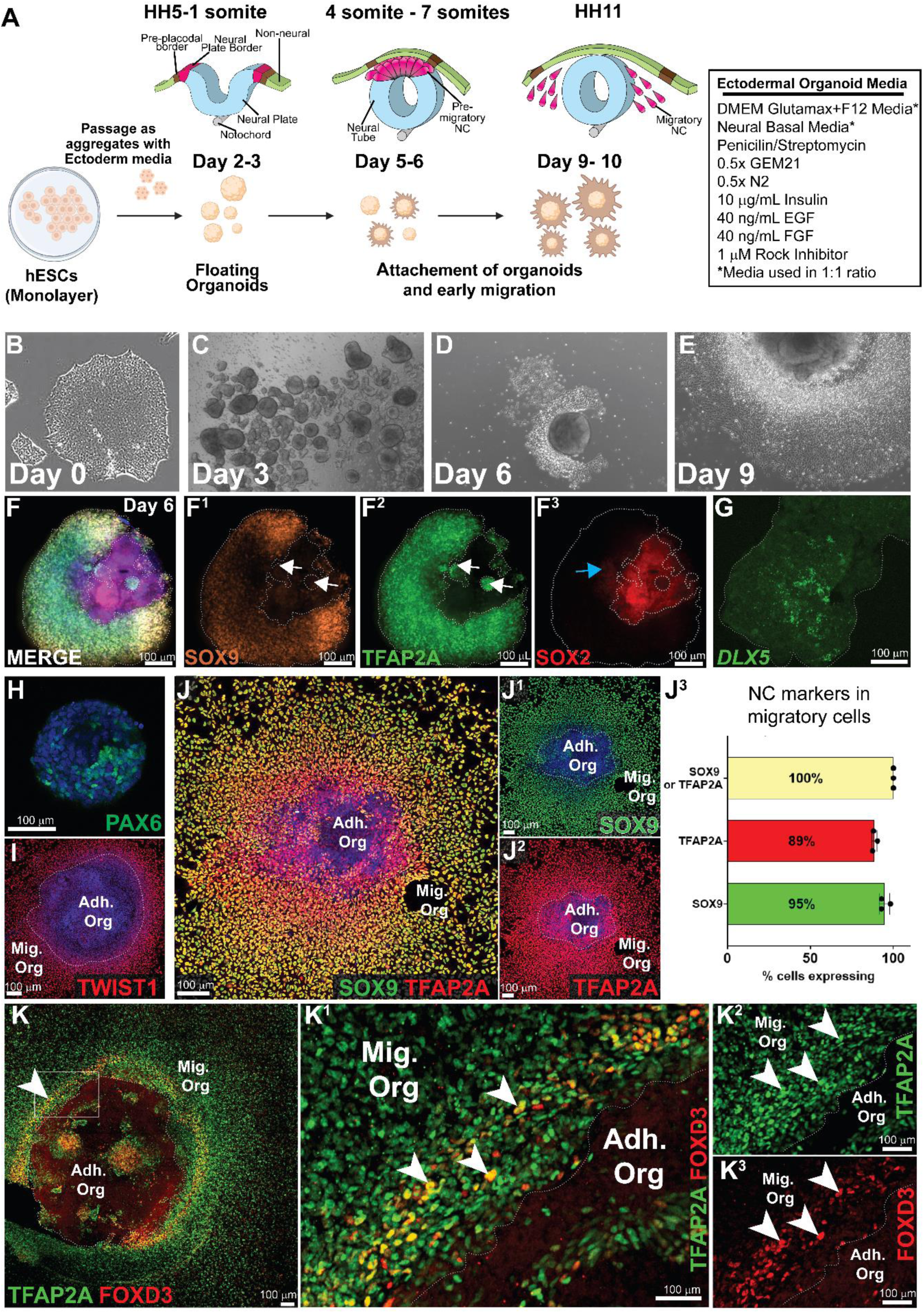
Image analysis shows that ectodermal organoids consist of neural, neural crest and non-neural ectoderm domains and generates a pure population of migratory neural cells. A) A cartoon of the ectodermal organoid culture protocol and equivalent stages in vivo. B) ESCs. **C**) Newly formed organoids. **D**) An adhered organoid with newly emigrated migratory NC cells. **E**) An adhered organoid with a large mat of migrating NC cells. **F**). NC markers SOX9 and TFAP2A mostly overlap in the floating epithelial organoids although TFAP2A also shows individual (NNE) immunopositive domains (white arrow, **F^2^**). The neural stem cell marker SOX2 depicts a neural domain, and SOX2/TFAP2A double positive dells (blue arrow **F^3^**) depict an immature (NPB) region. **G**) DLX5 HCR *in situ* demonstrates a NNE domain. **H**) PAX6 another early neural stem cell marker depicts the neural domain. **J)** Migratory cells are a pure NC population. **J^3^)** SOX9/TFAP2A 100%, SD= 0; TFAPA 88.58%, SD= ±1.58 SOX9 94.63%, SD= ±3.089. I) FOXD3 expression in premigratory and newly migrating NC cells. **K)** The migratory cells proximal to the adhered organoid expresses FOXD3 (white arrowheads).

### Serial Bulk RNAseq analysis of consequtive days of culture reveals temporal transcriptional changes in neural crest development similar to vertebrate embryos

To further validate whether the *in vitro* ectodermal organoids faithfully mimic the temporal changes of embryonic development, we performed bulk RNA sequencing on the organoids every second day of the ten-day culture protocol. At the last time point, we collected the remaining floating organoids separately (Day 10), as well as adhered organoids together with the migratory neural crest mats in another sample (Day10 Mig NC). The PCA plot demonstrates that the replicate samples from each day clustered together while they clearly separated according to collection day, which consisted of ∼70% of the variance (Fig. 2A). Overall, transcriptional changes were highest between the switch from embryonic stem cells to the neural crest medium (day 0 vs 2), substantial between days two and six of ectoderm patterning, modest between days six and eight, and again substantial between days eight and ten when the neural crest cells adhere and migrate out (Supp. Fig. 1A). Furthermore, stem cell related GO-terms were prevalent in the early samples (days 2-4) and cell to substrate adhesion and extracellular matrix organization related terms were found at later stages on days 6-10 (Supp. Fig. 1B).

**Figure 2.**
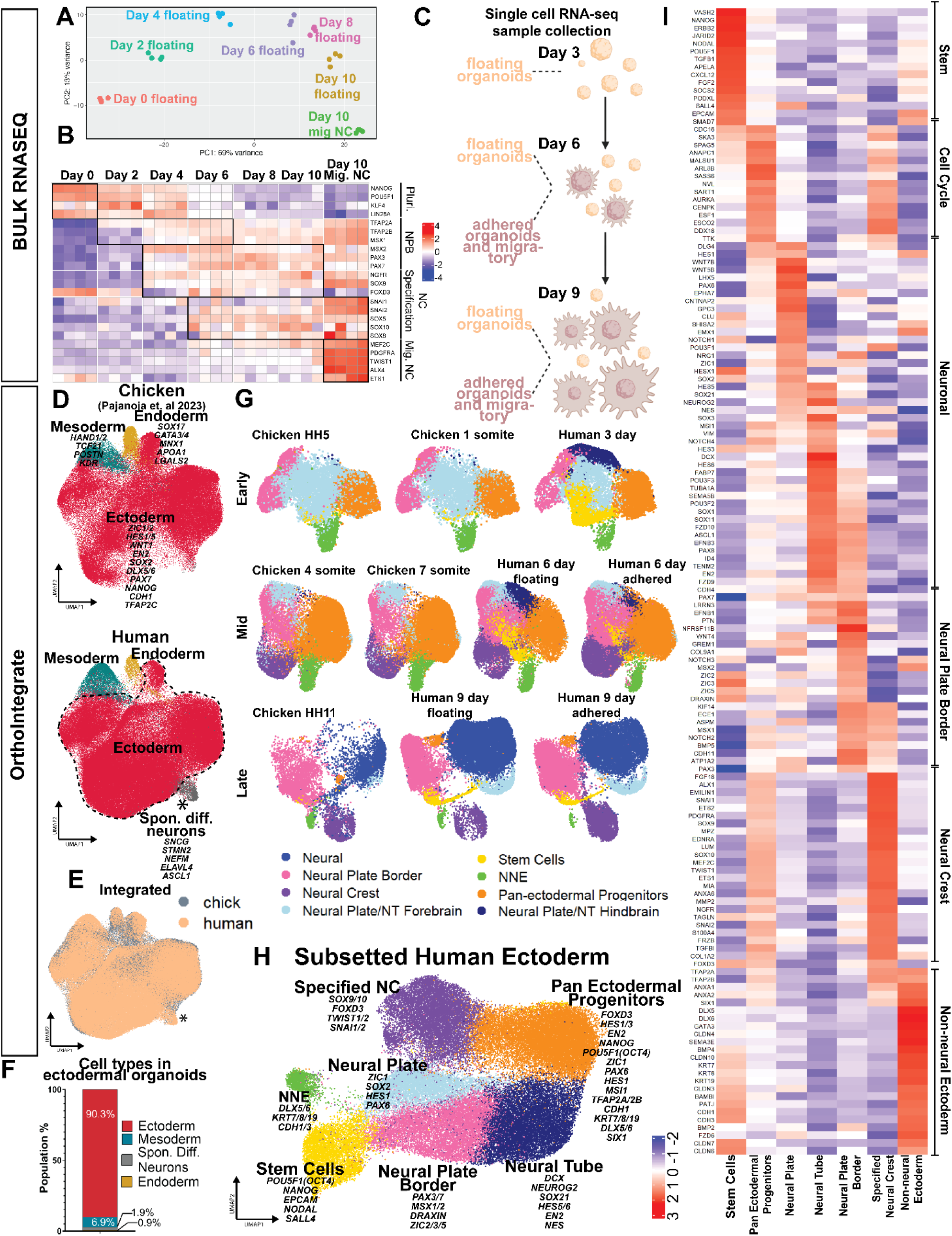
Bulk and scRNAseq analysis confirm transcriptional homology of ectodermal organoids to the ectodermal patterning process of avian embryos. **A)** PCA plot of the serial bulk RNAseq data set. **B**) Expression of conserved markers demonstrates in *vivo*-like maturation of NC cells. **C**) Cartoon of scRNAseq sample collection. **D**) Orthointegrate analysis reveals highly conserved gene expression between human ectodermal organoids and chicken embryo at the midbrain level. **E**) Chick-human integrated UMAP (**F**) ∼90% of the human organoids consist of ectodermal cells. **G**) Stage by stage Orthointegrate comparisons reveal high similarity between chick midbrain and human organoid cell populations during early, mid and late neurulation stages. Human organoids additionally consist of hindbrain CNS neural cells, and younger stage (equivalent to HH3/HH3 chick) ectodermal stem cells not included in the chick data set. **H)** Annotations of the subsetted human ectodermal cells shows presence of all ectodermal domains. **I**) Differentially expressed genes of the ectodermal subpopulations in the ectodermal organoids.

Next, we generated a heat map plotting known markers of the maturation process from early neural crest induction at the neural plate border to specification and EMT stages, followed by genes known to be highly expressed during neural crest migration (Fig. 2B). The heatmap shows a clear temporal correlation of the gradual maturation process of the neural crest according to the culture time. Early induction genes like TFAP2A were expressed on day two. At day three, more neural plate border genes like MSX2, PAX3/7, SOX9 and TFAP2B started to emerge. At day six, the neural crest specifier genes like SOX10, SNAI2 were expressed indicating a bona fide premigratory state and onset of EMT. Similar to the embryo, FOXD3 was first expressed in the embryonic stem cells but downregulated at the neural plate border stage and then re-expressed at the premigratory stage, followed by downregulation at the latest stages where neural crest cell migration has advanced, also confirming the immunostaining results. By day 10, expression of migratory and mesenchymal markers like PDGFRA and TWIST was high in the samples containing migratory halos of the neural crest cells, whereas their expression levels were much lower in the samples only containing the floating organoids (Fig. 2 B). Likewise, the temporal developmental dynamics of CNS neuronal and epidermal maturation followed the *in vitro* time course in the organoids (Supp. Fig. 1 C-D). Collectively, our immunostaining and bulk RNA sequencing results suggest that the 10-day 3D *in vitro* conditions promote spontaneous formation of ectodermal organoids that consist of a temporally maturing combination of ectodermal cell types mimicking *in vivo* development during neurulation.

### scRNAseq analysis confirms transcriptional homology of ectodermal organoids to the ectodermal patterning process of avian embryos

Next, to build a detailed picture of the transcriptional changes during developmental events in the organoids, we generated single cell RNA sequencing (scRNAseq) datasets that were, respectively, collected from the unattached floating organoids at respective days three, six, and nine, as well as from a mixture of adhered organoids and migrating neural crest cells at day six (when their halos are small consisting of newly migrated cells) and at day 9 (with large migratory neural crest halos, Fig. 2 C). To validate that our organoids were undergoing ectodermal patterning, we integrated our newly generated 3D human organoid dataset with our serial dataset from chicken embryo midbrain sections ^7^ by using OrthoIntegrate, which clustered cells in both species based on mutual ortholog expression ^41^. The results show high level of overlap between the chick and human cells (Fig. 2 D,E) and also confirmed that the majority (90%) of the cells in the cultures displayed ectodermal features justifying their description as ectodermal organoids (Fig. 2F). Additionally, we could identify small amounts of cells that had converted to mesoderm (6.9%) and endoderm (0.9%) based on expression of known marker genes for each of the germ-layers, such as *HAND1/2, TCF21, POSTN,* and *KDR* for the Mesoderm ^42–45^; *SOX17, GATA3/4, APOA1,* and *LGALS2* for the Endoderm ^46–48^; and *ZIC1/2, HES1/5, PAX7, WNT1, and TFAP2C* for the Ectoderm ^49–51^. We also detected a small population of spontaneously differentiated neurons (1.9%) that express high levels of neural developmental genes *SNCG, ASCL1, and ELAVL4* that were present already at Day 3 (Figs. 2D,E) but missing from the chick counterparts. Therefore, this cluster of cells were unlikely to have formed through the route of gradual neural crest differentiation, and we excluded it from further analysis. Next we subsetted the ectodermal cells and re-analyzed them stage by stage by using the OrthoIntegrate, which further confirmed the high transcriptional similarity to *in vivo* development (Fig. 2F). As expected, the noticeable differences were in line with sample collection: the chick data did not include the earliest ectodermal stem cells of the gastrula since the samples were collected right after gastrulation, and unlike the human organoids that contained cells of the entire cranial region, the chick samples were only collected from the midbrain region. (Fig. 2G).

### Ectodermal organoids are actively ongoing the whole process of ectodermal patterning - they consist of cells from ectodermal stem cells to immature and fully specified neural crest, future CNS and skin

The UMAP of the ectodermal cells separated them into seven unsupervised clusters, which we annotated by analyzing the lists of differentially expressed (DE) genes in each population (Supp. Table 1, Fig 2H). As showcased with a selection of DE and previously known marker genes for each group (Figs. 2H,I), the populations consisted of stem cells (yellow) with high expression of the pluripotency markers *POU5F1* (*OCT4*), and *NANOG*, and the early ectodermal adhesion marker *EPCAM*, known to be expressed in the entire ectoderm at early stages after gastrulation ^7,52^. Of note, this population also expressed *NODAL*, a landmark of the primitive streak that orients gastrulation, which is in line with the small amounts of mesoderm and endoderm detected in the organoids. We also identified another population of undecided Pan-Ectodermal progenitors, **PEP** (orange) that expressed lower levels of the pluripotency genes together with low levels of markers of all the ectodermal domains: non-neural ectoderm (**NNE**), neural tube (**NT**), and neural crest (**NC**), similar to our findings in the chick embryo ^7^. To further understand the transcriptional state of the PEP cells, Gene-Ontology analysis prediction highlighted that they were particularly active in RNA metabolism and biogenesis, cell cycle regulation and translation (Sup fig 1E). Consequently, the three transcriptionally distinct ectodermal domains that form during *in vivo* ectodermal patterning were all present in the ectodermal organoids. We identified two populations of developing neurons that expresses the neural cadherin *CDH2*: The neural plate / neural stem cell cluster (light blue) that expresses early neural induction markers like *SOX2*, *HES1, HESX1*, and *ZIC1* and *NES*, ^53,54^, and a neural tube population (dark blue) that already includes genes of the maturing cerebral cortex and spinal cord like *, NEUROGENIN2, SOX3/21, HES5/6, ASCL1* and *EN2* ^54–56^. The cells in the cluster that contain non-neural ectodermal and placodal cells (green) expresses typical early epidermal markers like *DLX5, KRT7/18, CHD1, SIX1,* and *EYA2* ^57^. Finally, neural crest development is also represented in two transcriptionally distinguished developmental populations: cells in the neural plate border cluster (pink) express the known markers of neural crest induction including *MSX1/2, PAX3/7, ZIC2/3/5* and the expression of specified neural crest genes like *SNAI2* and *SOX9* is emerging, while the cells in the purple population express high amounts of these markers, reflecting both premigratory/EMT stage (*TFAP2B, SOX9/10, FOXD3, SNAI2, ETS2)* as well as the migratory neural crest cells that express *TWIST1/2, MEF2C, TBX3,* and *PDGFRA* (Figs 2G,H).

To reveal temporal details of the subpopulations, we plotted the clusters in the UMAP (Supp. Figs. 2A,B), and the maintenance of their transcriptional signatures separately for each culture day and sample type (Supp. Fig. 2C). While the stem cell population represented a quarter of the cells on day three, their number was reduced in the later timepoint clusters. Similarly, the representation of non-neural ectoderm cells was ∼ 4% at Day 6 but reduced to less than 1% by day nine, suggesting that the ectodermal organoid culture conditions don’t promote further maturation of the epidermis. The pan-ectodermal progenitor cell cluster remained large (∼20-30%) throughout the cultures although they also gradually decreased their expression levels of the pluripotency genes (Supp. Fig. 2C). The neural tube cluster was proportionally highest represented in the floating organoids on day nine (36%) and the neural plate border cluster was highest represented in the floating organoids (∼20%). Lastly, the specified neural crest cell cluster only appeared at day six and was highest represented in the adhered organoids. In summary, the scRNAseq data confirms the findings from the bulk RNAseq analysis and reveals high transcriptional homology to ectodermal patterning of the chicken embryo where the maturation of the ectodermal domains occurs in a time-dependent manner. (Figs. 2C-H; Supp. Figs. 2A-C).

### Developmental trajectories predicted by RNA-Velocity and pseudotime resemble the timeline of embryogenesis

To better understand the dynamics of the developmental trajectories, we used RNA Velocity, which predicts cell state progression based on splicing dynamics (unprocessed vs. processed mRNA) and overall RNA expression ^58^. First, we re-clustered the cells from the ectoderm specific subset (Fig. 2F) based only on the expression of transcription factors allowing an analysis on the dynamics of the transcriptional regulatory networks of each developmental stage (Fig. 3 A)^59^. Overall, the results show a clear trajectory from stem cells via the pan-ectodermal progenitors to non-neural ectoderm, and the neural plate and neural plate border cell clusters, which further continue into the neural tube and specified neural crest clusters, thus mimicking the dynamics of ectodermal patterning in the embryo of various vertebrate model organisms (Fig. 3 A). To analyze the trajectories in more detail, we split the UMAPs to investigate each stage separately (without changing the analysis). In sum, these results show that the organoids mature in a faithful manner over biological time like in the embryo. Importantly, after the first days of culture, the organoids maintain an active ectodermal patterning process where the stem cells are the ongoing source of ectodermal derivatives, which transition via a pan-ectodermal progenitor state (Figs 3B-F).

**Figure 3.**
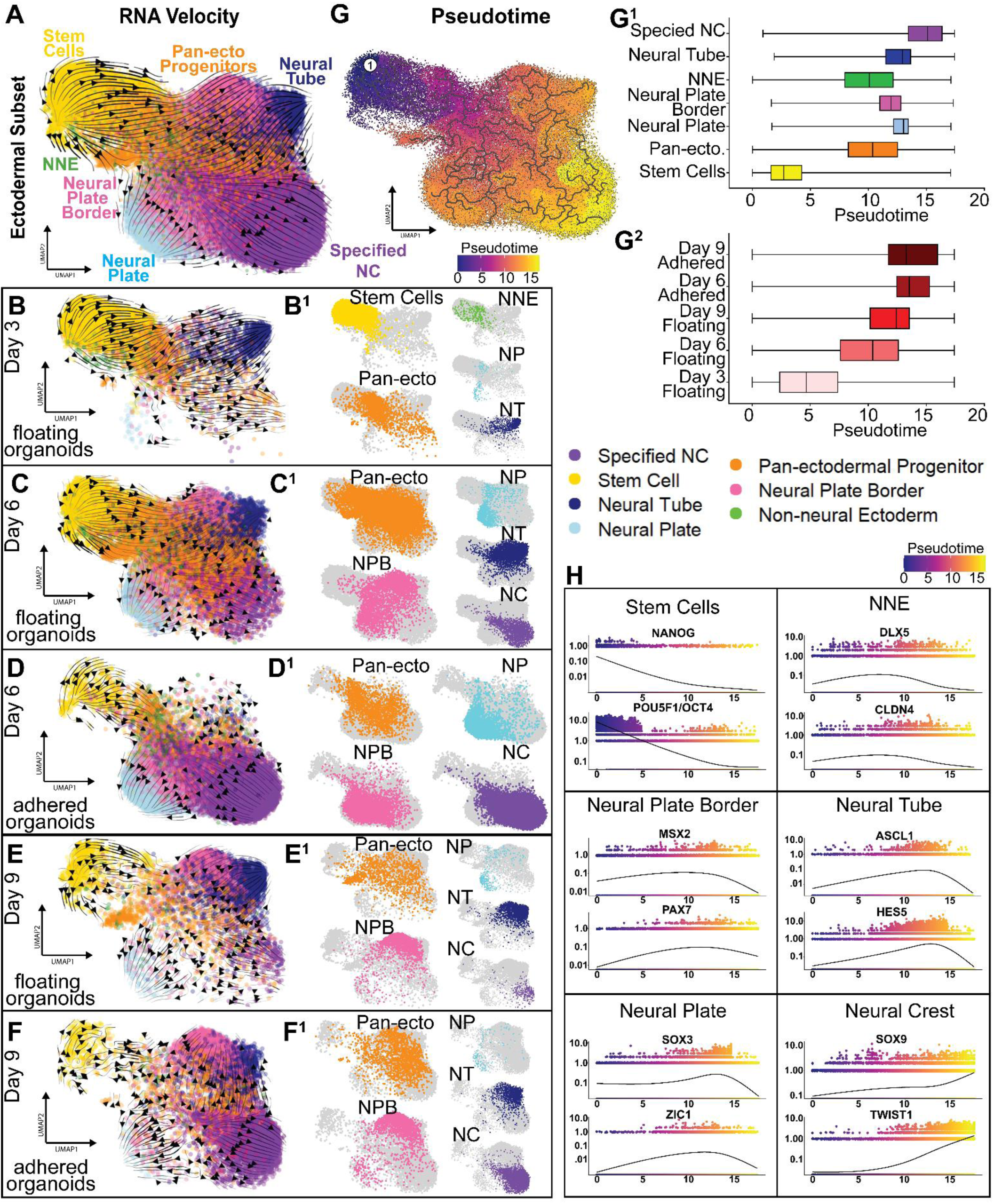
Developmental trajectories predicted by RNA-Velocity and pseudotime resemble the timeline of embryogenesis. **A)** RNA velocity projected on UMAP of ectodermal subset based on transcription factor expression show a trajectory of ectodermal stem cells giving rise to pan-ectodermal progenitors, which diverge to all domains. Neural and NC populations rise via neural plate and neural plate border. **B-F**) RNA velocity predictions on individual culture days of floating and adhered ectodermal organoids show consistent pattern resembling in vivo development. B’-F’) Individual featuring of respective clusters on the UMAP. **G**) Pseudotime reconstruction of the cell populations shows a similar, embryo-like developmental trajectory, (G^1^) predicting the longest pseudotime for the development of NC cells and shortest for the stem cells. (G^2^) The pseudotime predictions of different stage samples align with their biological age. **H**) Pseudotime of selected genes shows e.g. that the pluripotency gene NANOG has highest levels of expression in cells with the shortest pseudotime, and the migratory NC marker TWIST1 in the cells with the highest pseudotime.

Pseudo-temporal reconstruction using monocle3 also predicted a similar developmental trajectory where the stem cells progress to the pan-ectodermal progenitors, and it takes the longest pseudotime for the specified neural crest cluster to form (Fig. 3G)^60^. Individual graphing of pseudotime for each of the annotated cell clusters further supports the result showing that the stem cells have the shortest pseudotime, followed by the pan-ectodermal progenitors and the non-neural ectoderm, and the neural crest takes the longest pseudotime to form (Fig. 3G^1^). Also, as expected, the pseudotime was the longest for the biologically oldest samples, and interestingly, longer for the adhered than the floating organoids, most likely reflecting the larger proportion of NC cells in them (Fig. 3G^2^). Next, we plotted individual pseudotime dynamics for essential example genes of each cell type cluster. The results show similar dynamics as expected from embryonic data. The pluripotency genes *NANOG* and *OCT4 (POU5F1)* are expressed the highest in the cells with earliest pseudotime and their expression gradually declines as pseudotime progresses (Fig. 3H). Similarly, the neural plate border (*MSX2*, *PAX7*), non-neural ectoderm (*DLX5*, *CLDN4*), and neural plate (*SOX3*, *ZIC1*) genes start to be expressed in cells with early pseudotime and peak at mid-stage cells. Finally, genes of the most mature populations (neural tube *ASCL1*, *HES5* and neural crest *SOX9*, *TWIST*) gradually ascend and peak in the cells with the highest pseudotime (Fig. 3H).

### Cell-to-cell signaling of the ectodermal organoids reveal highly conserved mechanisms and novel interaction details

Proper ectodermal patterning requires a tight regulation of multiple signaling pathways. To better understand what kind of signaling is occurring in the ectodermal organoid model, we utilized another bioinformatics tool known as CellChat (v2) ^61^. This tool predicts potential cluster to cluster signaling interactions in single-cell RNA sequencing data sets (Fig. 4). We zoomed in on the signaling pathways that showed strongest interactions. As expected, these included FGF, WNT, and BMP signaling, which have established roles during ectodermal patterning in the embryo ^62–64^ (Figs 4A-C). Additionally, Midkine signaling (MDK), which mostly is known for its increase in cancer progression and poorly understood regarding its role in embryogenesis ^65^, is strongly communicating as a sender and receiver between all the ectodermal cell types as well as with the mesoderm in the organoids, except the neural crest and the stem cells that only receive the signal (Fig 4D). At the early stage, FGF signalling is produced by the stem cells and received by themselves, the early ectoderm (NP, NPB and NNE) as well as the endoderm. Specifically, the stem cell cluster send signals of FGF2 that is mediated via FGFR1 throughout the culture time in the floating organoids, but the signal is lost in the adhered organoids. Additionally, at Day3, the emerging neural tube secretes FGF2 to signal with other ectodermal populations and the stem cells. Finally, at Day 6, the newly specified neural crest cells in the adhered organoids are the source of FGF18, mostly known for its role in the developing posterior ectoderm and somites ^66,67^ which is received largely by all the ectodermal clusters and the mesoderm. These interactions are in line with reports on how FGF2 signaling maintains pluripotency and how FGFs also regulate neural stem cell maintenance and neural crest formation at the neural plate border ^68,69^ (Fig 4A and Supp. Figs. 4-8). Notably, at the neural crest specification and end of neurulation stage, the endoderm cluster is a strong source of FGF17, which is received by the entire ectoderm and the mesoderm and endoderm itself via FGFR1 (Supp. Figs. 4-8) – in line with reports that FGF17 plays a role in forebrain development, although details on early cranial development remain unstudied ^70^. Importantly, although on moderate levels, all the FGF interactions are also received by the pan-ectodermal progenitor population, which is predicted to have the lowest amount of overall signaling interactions and did not receive input from other signaling pathways (Fig. 4E and Supp. Figs. 4-8),

**Figure 4.**
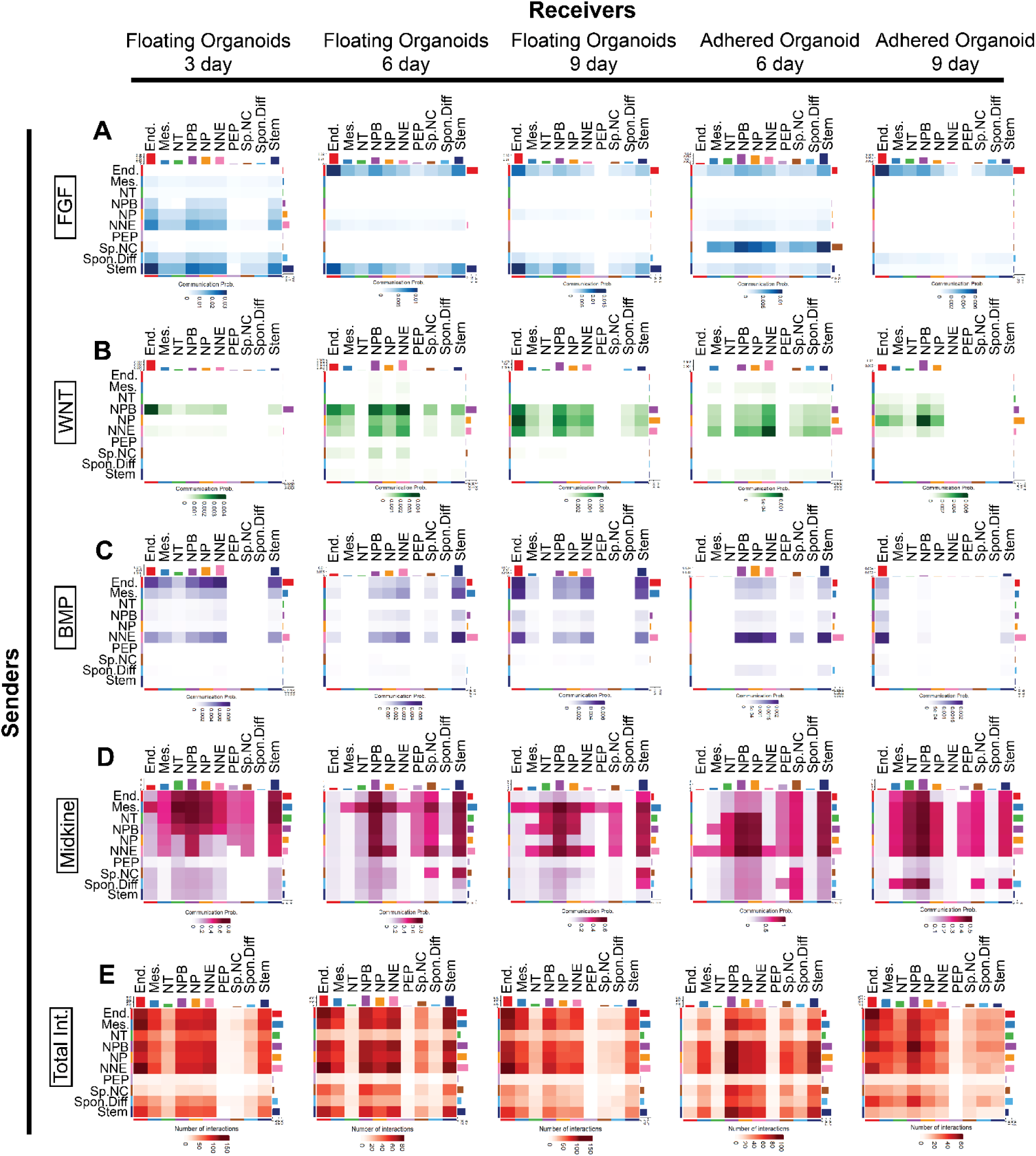
Cell-to-cell signaling prediction of the ectodermal organoids reveal highly conserved mechanisms and novel interaction details. **A)** Stem cells are the continues strong source of FGF signaling to itself and the NNE, NPB, and NP, and at Day 3 all four populations both send and receive the signal. In the adhered organoids at Day 6, the specified NC also sends FGF signals, which are reduced in the adhered Day9 cells. The small population of endoderm cells also actively produce FGF **B**) WNT signaling is highest in Day 6 and 9 floating organoids where NPB NP and NNE are the strongest signaling sources and receivers. Endoderm also receives signal. **C**) NNE send BMP signals throughout the culture days, which are received by the NNE, NPB, NP, the stem cells and the endoderm, which also produces BMP. BMP signaling is turned down in Day 9 adhered organoids. **D**) The developing ectoderm produces and receives strong midkine signaling across all populations and developmental days. **E**) Overall signaling strengths predict active cross talk between the ectodermal cell populations and with both mesoderm and endoderm. PEP cells are not participating in signaling events as actively as other populations. The small population of spontaneously differentiated neurons (that are an ectopic population not found in the chick data set), also receive and send significantly less signals than other populations. End=endoderm, Mes=mesoderm, NT=neural tube, NPB=neural plate border (early crest), NP=neural plate (neural stem cells), NNE=non-neural ectoderm, PEP= Pan-ectodermal Progenitors, Sp NC (specified neural crest), Spon Diff=(Spontaneously differentiated neurons), Stem=Ectodermal Stem Cells (The specific ligand -receptor pairs are shown in Supp figures 4-8). NNE: Non-Neural Ectoderm, Stem: Stem Cells, NPB: Neural Plate Border, Sp.NC: Specified Neural Crest, PEP: Pan-ectodermal Progenitors, NP: Neural Plate, NT: Neural Tube, Mes: Mesoderm, End: Endoderm, Spon.Diff: Spontaneously Differentiated Neurons.

WNT signaling, on the other hand, is first sent by the neural plate border cluster and received mostly by the endoderm, but later, it is exclusively produced and received by all the early ectoderm clusters NPB, NP and NNE (Fig. 4B). Specifically, throughout the stages, WNT4 ligand is secreted from the neural plate border and it interacts strongly with all cell clusters via a different, cell type specific combination of the receptor (FZD2/374/5/7 + LRP5/6) (Supp. Figs. 4-8). On the other hand, at days six to nine, WNT6 is secreted by the non-neural ectoderm and the signal is mediated to all clusters via FZD2/3/6/7 + LRP6. The third notable interaction is predicted between WNT7B secreted from the neural tube that communicates with all clusters (except endoderm) mainly in the adhered organoids at day six. These predictions are in line with established literature on how WNT signaling shapes the ectoderm and its particular importance to neural crest development^71^ and bring novel information on the specific ligands and their sources that remain less studied in general, and entirely unknown in human ectoderm.

Finally, the two consistent sources of BMP signaling are the endoderm and the non-neural ectoderm, and to some extent the mesoderm, which signal to themselves and also to the early ectodermal clusters NP, NPB, NNE and the stem cells (Fig. 4C). The role of different gradient levels of BMP signaling, and especially BMP4, in dictating ectodermal patterning as well as the dorsoventral axis of the neural tube has been established for decades *in vivo* and validated *in vitro*, but the ligand receptor sources and strengths in the process are poorly known ^72,73^. Interestingly, the results from the predicted interactions in the human ectodermal organoids show that signaling of the endodermal BMP2 to the neural tube, non-neural ectoderm and the stem cells via BMPR1A + ACVR2B provides a stronger connection throughout the stages in the floating organoids than BMP4, which is produced by the non-neural ectoderm and received by all the ectodermal clusters (Supp. Figs. 4-8). Similarly, mesodermal BMP4 signaling to the ectodermal cell clusters via the same receptors is strong throughout the stages in the floating organoids. Additionally, the ectodermal clusters, including the stem cells, receive significant BMP7 input from the non-neural ectoderm, the neural plate border, and the neural tube in the floating organoids from day three to nine. In sum, these results confirm that the predicted signaling interactions are in line with literature on *in vivo* ectodermal development via BMP signaling and bring novel details on the specific ligands and their sources in the process. Finally, CellChat analysis revealed that the small group of spontaneously differentiated neurons only modestly contributed to the overall signaling suggesting that they truly are an ectopic population that is not required for the ectodermal patterning process in the floating organoids. On the other hand, the small endoderm and mesoderm clusters actively communicate with the ectoderm as signaling sources and receivers in the floating organoids. The results suggest that the presence of all germ layers is an essential part of the ectodermal patterning process, which is often overlooked in development as many studies focus on the downstream effects of a signaling pathway in a single cell type without addressing the endogenous source.

### Ectodermal organoids posteriorize spontaneously over time

Although the central nervous system, neural crest, and epidermis develop along the entire body axis from head to tail, their characteristics vary significantly depending on the axial level—for example, the brain versus the spinal cord. Similarly, only the cranial epidermis contributes to sensory placode formation. Apart from the peripheral nervous system and melanocytes, axial-level specificity is also evident in neural crest derivatives: only at cranial levels, they give rise to the craniofacial skeleton, cranial sutures, pericytes of the meninges, and mesenchyme of the thyroid and thymus. In contrast, only neural crest cells originating from the vagal region uniquely contribute to formation of heart valves and septum as well as the enteric nervous system, while trunk-level neural crest cells generate chromaffin cells of the adrenal medulla. Therefore, we sought for signs of axial level positioning in the organoids to further elucidate their identity. We noticed that although the organoids had an overall cranial expression profile, the neural plate cluster expressed higher levels of anterior, forebrain specific markers like *OTX1/2*, *SIX3*, *LHX2*, *FEZF1/2*, *EMX2*, *FOXG1*, *SP8*, and *NKX2-1*, whereas markers of the mid- and hindbrain level such as *GBX2*, *CNPY1*, *EN2*, *IRX1/2/3*, *RARB*, *EGR2*, and *NKX2-2* were expressed more in the neural tube population (Fig. 5 A) ^74–76^. These data suggest that the organoids contain cells from different axial levels, which in the embryo form gradually over time as the forebrain level is the first to form raising the possibility that the ectodermal organoids posteriorize in a time dependent manner.

**Figure 5.**
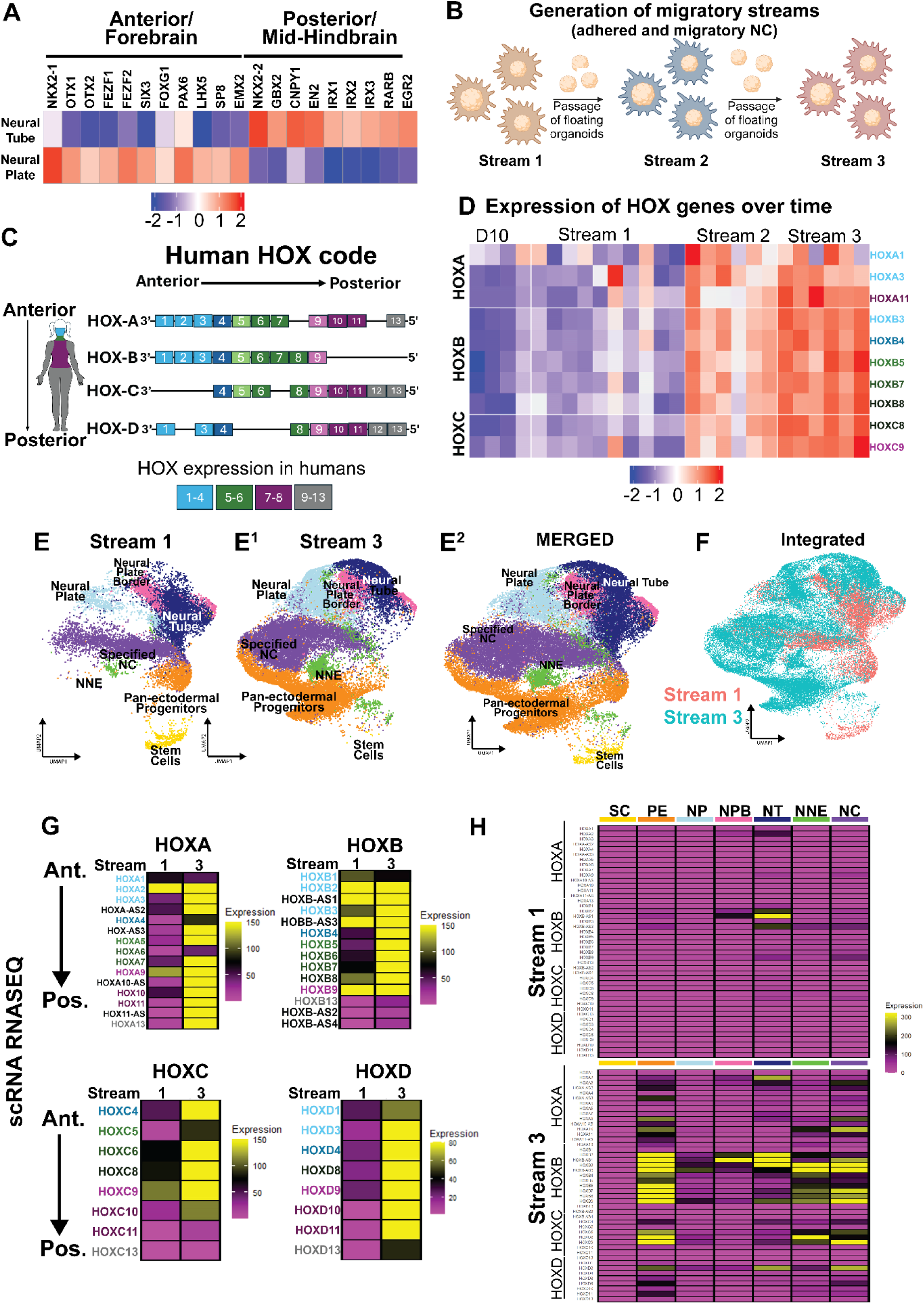
Ectodermal organoids posteriorize spontaneously over time. **A)** The neural plate cells on the scRNAseq data set express proportionally more anterior forebrain markers, and the neural tube cells express proportionally more mid-and hindbrain markers. **B**) A cartoon of the protocol of forming posterior ectodermal organoids. C) A cartoon of the human HOX genes located in a 3’ to 5’ sequence in the genome. **D**) Heatmap depicting expression of HOX-genes in the bulk RNAseq data showing increased expression over time. The first stream shows a cranial, and the third stream reflects a trunk transcriptional profile. **E**) Individual ectodermal subpopulations from scRNAseq featured on a UMAP of integrated samples of adhered organoids together with the migratory NC cells from Steam 1 and 3 shows representation of all cell types in both data sets. **F**) Integrated UMAP with separately colored origins of anterior and posterior streams. **G**) Comparison of expression levels of Hox genes (HOXA-D), shows posteriorization of the ectodermal organoid cultures in Stream 3. **H**) Comparative expression of HOX-genes in the individual cell populations of Stream 1 show anterior transcriptional profile. In stream 3, anterior and posterior Hox-genes are expressed particularly in the pan-ectodermal progenitors and the specified NC, NT and NNE, reflecting posterior axial levels. NNE: Non-Neural Ectoderm, SC: Stem Cells, NPB: Neural Plate Border, NC: Specified Neural Crest, PE: Pan-ectodermal Progenitors, NP: Neural Plate, NT: Neural Tube.

To test this hypothesis, we investigated if the organoids continue to posteriorize further if kept in culture for longer. For this, we separated the floating and lightly adhered organoids from the migratory mat and re-plated them on a new culture dish to test if new migratory neural crest would be generated. Indeed, the organoids re-adhered and produced a second migratory stream of neural crest cells, and after repeating the step once more, a third migratory stream was produced (Fig. 5 B), further evidencing the continued ectodermal patterning ongoing in the organoids. Bulk RNAseq samples that contained a combination of the adhered organoids, and the migratory cells were collected and, since HOX-genes are known to mirror the axial level identity across species (Fig. 5C), we tested whether their expression would reveal signs of posteriorization differences between the streams. Genes located at the 3′ end of the Hox clusters—associated with anterior identity—are activated earlier and expressed in more anterior regions of the embryo. In contrast, those positioned toward the 5′ end—linked to progressively more posterior fates—are expressed later and in increasingly posterior domains ^77^ (Fig. 5C). Strikingly, as compared to the first stream, which according to nominal levels of HOX gene expression were cranial cells, the results show a clear increase of HOX gene expression in stream two, which was further strengthened in stream three that expressed high levels of the trunk axial level genes *HOXA11*, *HOXB9*, *HOXC8* and *HOXC9*, indicating that the organoids indeed spontaneously posteriorized by time (Fig. 5D).

To further validate these results, we generated single-cell RNAseq data from the third stream and merged it with our data from the first stream, which were thus both collected on their respective Day 9 from the combination of adhered organoids and the migratory mat cells (Fig. 5E^1–2^). The results show that all the annotated cell clusters from the first stream were also present in the third stream although the shape of the clusters differed. Notably, the first stream contained more stem cells, suggesting that, like in the embryo, the earliest pluripotency-like stemness decreased in the cultures over time. Furthermore, since the overlap was not perfect between the streams, we created developmental gene modules from each cell type cluster from the first stream and confirmed that they were consistently expressed in the corresponding clusters of the third stream (Supp. Fig. 9A) (Supp. Table 3). Additionally, we used Seurat’s SCT-integration function to combine the two streams, which confirmed the similarity of the two streams, which both contributed to the overall shape of the UMAP, although with different emphasis (Fig. 5F). Moreover, the stream 3 cells still had a similar pattern of expression of the ectodermal domains seen from figure 2 (Supp. Fig. 9B compare to Fig. 2I). Similar to the bulk RNAseq results, expression of HOX genes was significantly increased in the third stream suggesting the anterior-posterior transcriptional profiles were contributing to the differences on the UMAP (Fig. 5G). Notably, cluster specific analysis showed that the pan-ectodermal progenitors, the neural tube, the non-neural ectoderm, and the specified neural crest were the ones that had gained HOX gene expression in the third stream (Fig. 5H). Similarly, other posteriorizing signaling pathways like Retinoic acid converting enzymes and posterior WNTs were upregulated in the third stream (Supp. Figs. 9C,D). In sum, these data suggest that our ectodermal organoid model also has the potential for studying anterior-to-posterior axis development of ectodermal populations, and to use the appropriate axial level to induce neural crest cells depending on the derivative type of interest.

### Differentiation of nine diverse neural crest derivative cell types

Neural crest cells give rise to more than 30 different cell types ranging from ectodermal to mesodermal-like as well as exo- and endocrine cells (most of which are typically derived from the endoderm), which populate the embryo throughout the body. We tested the pluripotency-like stem cell potential of the neural crest cells formed in the ectodermal organoids by differentiating them to nine different cell types; melanocytes, peripheral glia, sensory neurons, sympathetic neurons, adrenal chromaffin cells, chondrocytes, osteocytes, adipocytes, and smooth muscle. By combining and optimizing the culture conditions based on established protocols, as well as by adding novel components to the media, a rigorous outcome in terms of quality and purity of the respective differentiated cell types was achieved as analyzed by bulk RNAseq, which showed differential expression of typical genes of each differentiated cell type neural crest ^78^, melanocytes ^79,80^, glial cells ^81–90^ sensory neurons ^91–93^ sympathetic neurons ^94–104^, chromaffin cells ^105–113^, chondrocytes ^114–121^, adipocytes ^102,122–126^, human embryonic stem cells (hESCs) ^127–133^, osteocytes ^134–141^, smooth muscle ^142–147^ as well as qPCR and quantitative single cell analysis from immunostainings of mid- and endpoints of the cultures that varied from 10 to 35 days (Fig. 6 A-J; Supp Figs 10A-G, 11A-C). Furthermore, the heatmap in figure 6A indicated higher transcriptional similarity between certain cell types. We next calculated correlation scores between the different lineages. The results show highest correlation between the neuronal and chromaffin cells and, on the other hand, between the osteocytes, smooth muscle and chondrocytes, as well as early melanocytes, adipocytes and glial cells which all correlate with each other, and show less correlation with the neuronal lineage group. (Supp. Figs. 10 I,K).

**Figure 6.**
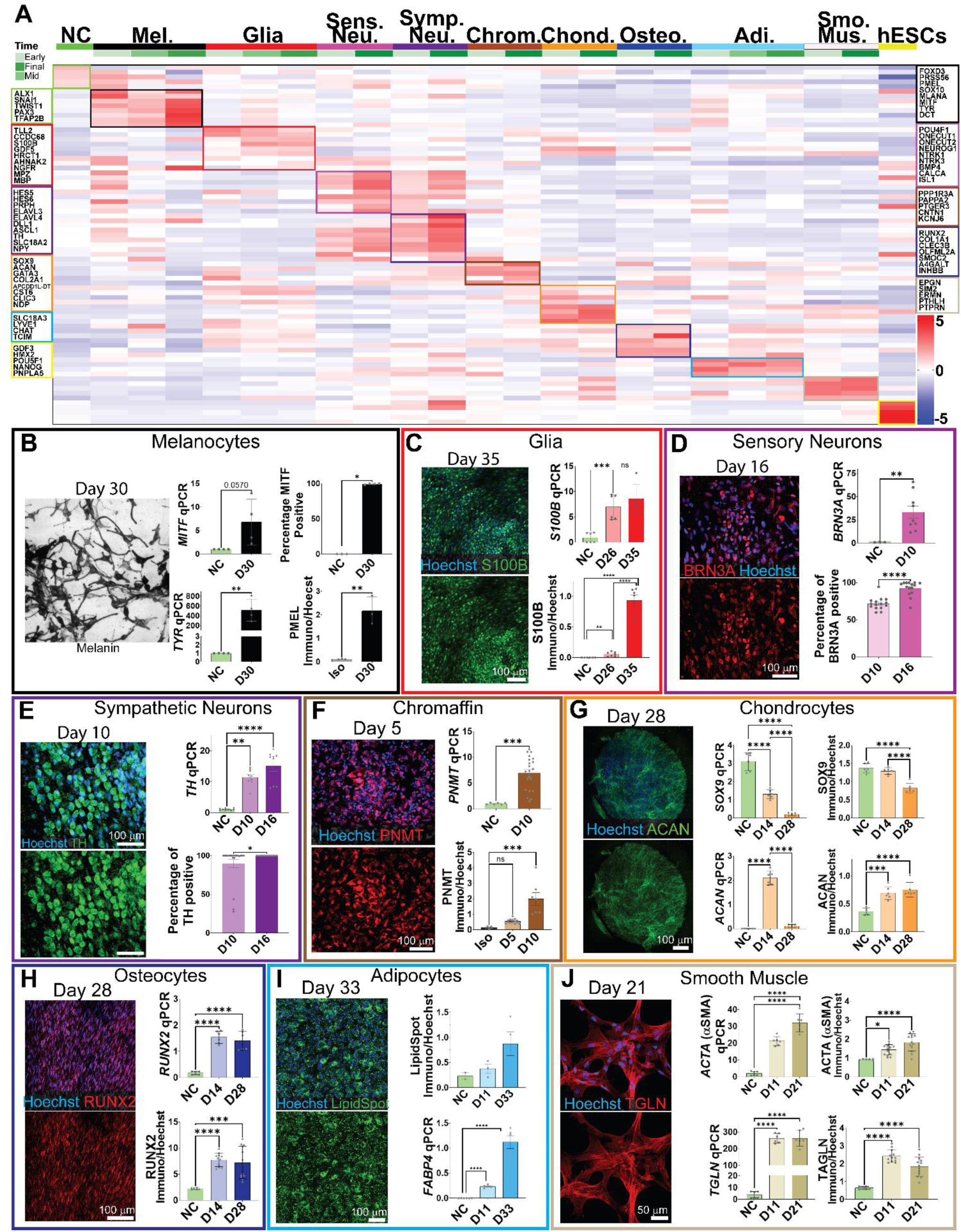
Differentiation of nine diverse neural crest derivative cell types from neural crest cells. **A)** A heatmap showing differentially expressed genes that also include known markers for each respective differentiated NC derivative cell type at different timepoints of early and fully differentiated stages. **B**) A 30-day protocol produces a pure population of pigmented MITF-immunopositive melanocytes that also express TYR and PMEL. *MITF* qPCR NC SD = ±0.002; Day 30 Melanocytes SD = ±4.919. MITF immuno percentage positive NC SD = 0; Day 30 Melanocytes SD = 0. *TYR* qPCR NC SD = ±0.008; Day 30 Melanocytes SD = ±223.7. PMEL immuno NC SD = ±0.021; Day 30 Melanocytes SD = ±0.6033. **C**) Virtually all cells in the glial cultures differentiated into S100B-immunopositive cells. *S100B* qPCR NC SD = ±0.9851; D26 SD = ±2.476; D35 SD = ±2.853. S100B immuno NC SD = ±0.0001; D26 SD = ±0.0355; D35 SD = ±0.2323 **D**) Virtually all sensory neurons express the cell type specific marker BRN3A. *BRN3A* qPCR NC SD = ±0.4091; D10 SD = ±20.05. BRN3A immuno D10 = ± 8.315, D16 = ±9.708. **E**) Symphatetic neurons differentiated into a pure population of TH-immunopositive cells that also significantly increased expression of TBH at the end of the 16-day protocol. *DBH* qPCR NC SD = ±0.6501, D10 SD = ±0.7824, D16 SD = ±5.811. DBH imunno D10 SD = ±21.46, D16 SD = ±0. *TH* qPCR NC SD = ±0.3162, D10 SD = ±2.576, D16 SD = ±5.205. **F**) Differentiation into chrommaffin cell lineage formed PNMT-positive cells in ten days. *PNMT* qPCR NC SD = ±0.1704, D10 SD = ±3.229. PNMT immuno Isotype SD = ±0.1115, D5 SD = ±0.1396, D10 SD = ±1.080. **G**) Chondrocyte cultures formed Aggrecan and SOX9 -positive mineralizing chondrospheroids in 28 days. The cartilage lineage initiation and NC marker SOX9 expression is highest in early and midpoint cultures. *SOX9* qPCR NC SD = ±0.4912, D14 SD = ±0.2429, D28 SD = ±0.07434. SOX9 immuno NC SD = ±0.1266, D14 SD = ±0.0939, D28 SD = ±0.1296. *ACAN* qPCR NC SD = ±0.0015, D14 SD = ±0.2845, D28 = ± 0.0596. ACAN immuno NC = ±0.0591, D14 SD = ±0.1144, D28 = ±0.1303. **H**) In 28 days, NC cells organized into RUNX2-immunopositive osteocytes that have nuclei synchronized in a symmetrical angle. *RUNX2* qPCR NC SD = ±0.0642, D14 SD = ±0.2367, D28 SD = ±0.3629. **I**) Adipocytes express increasing levels of *FABP4* during their 33 days on maturation resulting in a homogeneous culture of cells that form lipid-spot positive lipid droplets. *FABP4* qPCR NC SD = ±0.0010, D11 SD = ± 0.0288, D33 SD = ±0.2864. LipidSpot immuno NC SD = ± 0.0942, D11 = ±0.1615, D33 = ±0.4709 **J**) Transgelin and Smooth Muscle Actin expressing cells show a typical smooth muscle morphology after 21 days in a uniform culture. *ACTA* qPCR NC SD = ±1.269, D11 SD = ±2.676, D21 SD = ±5.413. ACTA immune NC SD = ±0.036, D11 SD = ±0.2735, D21 SD = ±0.4533. *TAGLN* qPCR NC SD = ±2.739, D11 SD = ±28.70, D21 = ±46.39. TAGLN immuno NC SD = ±0.0798, D11 SD = ±0.3547, D21 SD = ±0.5395. ISO= isotype IgG control (the images stained by additional antibodies are shown in Supp Fig 10A). Significance is annotated as such *: p-value > 0.05, **: p-value >0.002, ***: p-value >0.0002, ****: p-value >0.0001. NC: Neural Crest, Mel: Melanocytes, Sens Neu: Sensory Neurons, Symp Neu: Sympathetic Neurons, Chrom: Adrenal Chromaffin Cells, Chond: Chondrocytes, Osteo: Osteocytes, Adi: Adipocytes, Smo Mus: Smooth Muscle, hESC: human embryonic stem cells.

### Ectodermal organoids derived from DiGeorge patient cells show aberrant ectodermal pleistopotency and defected NC specification

Finally, to use our ectodermal organoids as a model for studying ectodermal birth defects, we applied it to iPSCs from a DiGeorge syndrome family of two patients and two healthy relatives as control. Since DiGeorge syndrome manifests with a broad range of symptoms in neural crest derived cell types, we analyzed the patient cells in comparison to controls at three time points: at the premigratory stage when neural crest cells are undecided regarding their future derivative lineage, at the migratory stage, and during differentiation into bone, cartilage and smooth muscle cells.

Gross morphological analysis of day 6 floating organoids did not reveal any obvious differences in the shape, or the size between patients and controls (Fig. 7A; Supp Fig. 12A). However, immunostaining for SOX9 and NANOG revealed a significant decrease in the expression intensity in the patient organoids (Figs. 7A-D). Pluripotency genes like NANOG are essential for establishing the pluripotency-like, pleistopotent stem cell capacity of the neural crest^7^ and these results indicate that DGS patient NC cells are not properly induced into a pleistopotent state, leading to incorrectly specified NC cells that express reduced amounts of NC markers like SOX9. Notably, the patient organoids were still able to transition to a migratory stage. However, the migrating patient neural crest cells expressed very low levels of SOX9 and TFAP2A as compared to the controls (Fig. 7E). Quantification of the number of cells in the migratory neural crest cells in the cultures over time additionally revealed severe growth retardation of the patient cells (Fig. 7F). However, the ability of the DGS patient neural crest cells to migrate was not statistically different from the healthy controls as measured by migration length in a scratch assay or by live imaging-based assessment of speed in individual cells (Supp. Figs. 12B,C). Finally, since DiGeorge patients present developmental defects in multiple mesenchymal neural crest derived cell types in the head and neck region, we differentiated them to bone, cartilage and smooth muscle cells. Formation of chondrocytes was severely impaired resulting in a dramatic reduction in the size of the chondrospheroids from the patient derived cells compared to the controls (Figs.7 G,H). Accordingly, the intensity of the chondrocytic lineage driver SOX9 was significantly decreased at the midpoint of the differentiation protocol (Supp. Fig. 12E). Based on gross morphology and cell type differentiation marker analysis, osteoblasts were less syncronized in the patient samples as compared to the stripe-shaped, organized controls, but expression levels of the osteoblast markers were not changed (Fig. 7J; Supp. Figs. 13 A-D). The phenotype suggests a more progenitor-like state and misregulated differentiation of the patient cells, where the mineralizing proteins ALPL and Osteocalcin are normally expressed despite the poor orientation (Supp. Figs. 13 C,D). Smooth muscle cells appeared morphologically normal (Fig. 7I; Supp. Fig. 13E) but the expression levels of SMA and MYH11 were slightly increased by immunostaining at the 21-day endpoint of the cultures (Supp. Fig.13F,G). In sum, neural crest cell differentiation to mesenchymal lineages was defective in the DGS patient cells, in line with the craniofacial anomalies of DGS.

**Figure 7.**
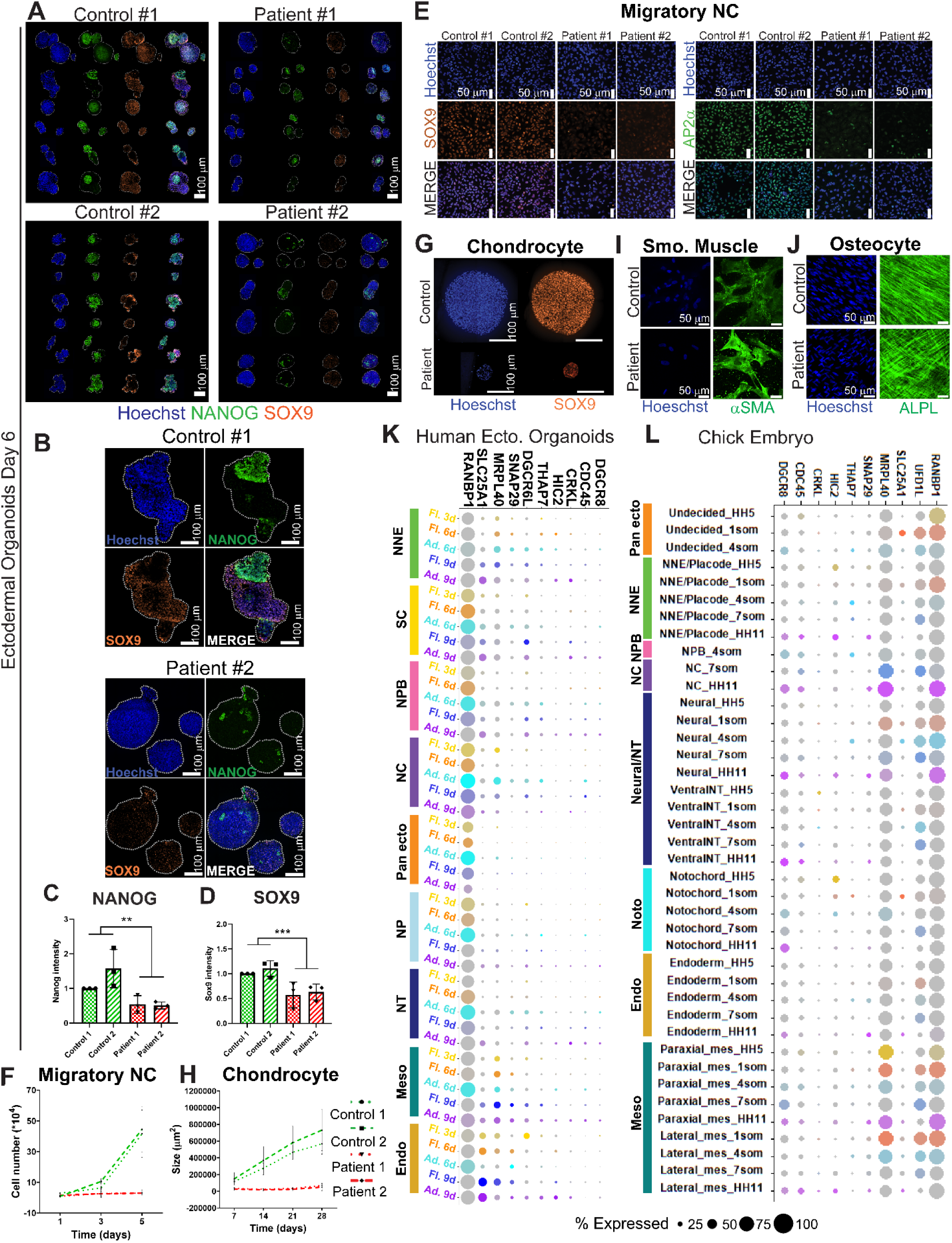
Neural crest cells from DGS patients show an early NC specification defect. **A**) Example images of floating ectodermal organoids from DGS patient cells and healthy relative controls immunostained with the ectodermal pluripotency marker NANOG and NC specification marker SOX9 at Day 6. **B**) Magnification of control and DGS patient example organoids. **C**) Quantification of immunostaining shows reduced expression of NANOG in floating DGS patient ectodermal organoids as compared to controls (Control 1 SD = ±7.422e-008, Control 2 SD = ±0.553, Patient 1 = ±0.2540, Patient 2 = ±0.1001) at day 6. n=3 biological replicates from 15-20 organoids per sample. **D**) Quantification of immunostaining shows reduced expression of SOX9 in floating DGS patient ectodermal organoids as compared to controls (Control 1 SD = ±2.887e-008, Control 2 SD = ±0.1547, Patient 1 SD = ±0.2601, Patient 2 SD = ±0.1603) at day 6. n=3 biological replicates from 15-20 organoids per sample. **E**) Example immunofluorescent images of migratory (P1) NC cells show reduced expression of SOX9 and TFAP2A in DGS patient cells. **F**) Quantification of cell counts of P1 migratory NC cells shows impaired growth of DGS patient cells over time (Control 1 SD = ±21.44, Control 2 SD = ±22.87, Patient 1 SD = ±1.127, Patient 2 = ±0.7373) **G**) Example images show reduced size of DGS patient derived chondrospheroids immunostained with SOX9 as compared to controls. **G**) Quantification of Chondrospheroids shows impaired differentiation ability of DGS NC cells into chondrocytes (Control 1 SD = ±200026, Control 2 = ±254270, Patient 1 = ±19753, Patient 2 = ±14051). H) Example image of DGS patient osteocytes with disorganized mineralizing Osteocalcin-immunopositive cells as compared to the organized controls **I**) Image based quantification of cell and nuclear (Supp fig 13 A) morphology shows that osteocytes from GDS patients have disorganized orientation as compared to contols. **J**) Dotplot highlighting the genes of the DGS microdeletion region that are expressed in the ectodermal organoids. **K**) Dotplot highlighting the genes of the DGS microdeletion region that are expressed in the different sucpopulations in the neurulating chicken embryo at midbrain axial level reveals high similarity with the expression profiles of ectodermal organoids in I. Significance is annotated as such *: p-value > 0.05, **: p-value >0.002, ***: p-value >0.0002, ****: p-value >0.0001. NNE: Non-Neural Ectoderm, SC: Stem Cells, NPB: Neural Plate Border, NC: Specified Neural Crest, Pan-ecto: Pan-ectodermal Progenitors, NP: Neural Plate, NT: Neural Tube, Meso: Mesoderm, Endo: Endoderm.

### Identification of ten DGS region genes that are expressed during ectoderm patterning

Due to this, we next analyzed the scRNAseq data set from WT organoids to ask which subpopulations express the DiGeorge genes in relevant amounts. Bubble plots were used to depict genes that were expressed in significant levels at least in 25% of the cells of at least one of the cell types in the organoids (Supp. Fig. 15). With this, the list of DGS was narrowed down to 10 genes that are potentially relevant for the early neural crest phenotype of the organoids (Fig 7I). *RANBP1*, that guides mitotic spindle assembly, was expressed in vast majority of the cells in the organoids but the intensity of its expression was highest in the specified neural crest and pan-ectodermal stem cells of the adhered organoids at day 6 ^148^. *MRPL40*, a protein shown to be necessary for mitochondrial ribosomal integrity and proteostasis in Drosophila synapses, had highest expression in the specified neural crest cells in the adhered organoids on Day 6, although it also was expressed both in the mesoderm and endoderm ^149^. Another mitochondrial protein, *SLC25A1*, which intriguingly was shown to interact with *MRPL40* in the same report, was not co-expressed with *MRPL40* in the ectodermal organoids but rather was expressed at Day 9 in the stem cells, non-neural ectoderm and neural plate border as well as in the endoderm. *DGCR6L*, a paralogue of the *DGCR6* gene, which also is in the same DGS locus, was expressed in roughly 30% of the stem cells and NPB cells at Days 6 and 9 as well as in the day 3 endoderm. Finally, *THAP7*, an epigenetic factor that augments proliferation was expressed in roughly a quarter of the neural crest cells and the neural palate border at Day 6, and *CRKL*, known for its oncogenic activity, was expressed roughly in a quarter of the stem cells and NNE and NT at Day 9 ^150,151^. Expression of *HIC2*, a transcriptional repressor associated with cardiac development but also an activator of *SIRT1* expression was found in a small proportion of NNE and NPB cells as well as in endoderm at Day 9 ^152^. Finally, a small proportion of NC, Stem cells and NNE cells expressed *SNAP29*, a SNARE-protein involved in membrane fusion, autophagy related to tissue homeostasis and remodeling, and protein trafficking, ^153^, which was also expressed throughout the stages in roughly a third of the cells in the mesoderm and endoderm clusters (Fig. 7K) To gain further validation of the expression of the DGS genes, we examined which of the DGS genes are expressed *in vivo* between gastrulation and end of neurulation in the chicken cranial region, which is the corresponding developmental time represented in the organoids. As shown by bubble plots, the expression patterns of the DGS genes in the chick embryo are very similar to the findings in the organoids (Fig. 7L, Supp. Fig. 16). To summarize these findings, we identified ten DGS region genes that are not functionally linked to neural crest before and are likely candidates for causing a loss-of-NANOG and consequent NC specification defect prior to neural crest migration. (Figures 7K and 7L).

To build a more comprehensive picture of the changes caused by DiGeorge syndrome on ectoderm and neural crest development, we performed bulk RNAseq analysis on the newly specified, premigratory neural crest cells (Day 5 organoids, which consist of all the cell types in the organoids in addition to neural crest) as well as on the cells of the primary, late migratory neural crest cells (Day12, passage 0) as well as the first migratory passage (P1), which both consist of only neural crest cells. To exclude the possibility that the phenotypic differences originate from changes at the pluripotent iPSC stage, we confirmed that expression of the core pluripotency complex genes were unchanged at day 0 in controls vs patients (Supp. Table 4). The results also validate that the DiGeorge genes (that were expressed in detectable levels in the bulk data set) are expressed at higher levels in the healthy control organoids, while a few show higher RNA expression levels in the patients despite the deletion (Supp. Fig 14). Importantly, while the heatmaps show a difference in the relative gene expression levels between controls and patients (Supp. Fig. 14), our scRNAseq data from WT samples, as indicated before, showed that most of the genes are expressed in nominal, biologically irrelevant levels (counts of less than 50 transcripts per sample). First, to validate our immunofluorescence-based characterization, we analyzed transcriptional changes of known neural crest markers in the DGS patient cells. The results of the premigratory stage show that rather than the neural crest gene regulatory network being simply downregulated, we detected a deranged specification process. The pluripotency genes *OCT4* (*POU5F1*) and *NANOG*, and other stem cell markers like *LIN28A/B*, neural plate border (*PAX3/7*) and later key neural crest specifier genes (*FOXD3*, *SOX10*) as well as the WNT signaling mediator *DRAXIN* ^154^ were downregulated, but several other markers of early induction, late specification, and EMT (*TFAP2A/B/C*, *MSX1/2*, *SNAI1/2*), were instead prematurely expressed as compared to the healthy relative control cells ^155–157^ (Fig. 8A). These genes are also largely involved in lineage commitment of cranial neural crest derived mesenchymal cell types ^158,159^. These results indicate that the compromised maintenance of ectodermal stem cells impairs a proper neural crest induction process, which results in incorrectly specified neural crest cells. Intriguingly, the downregulated genes are crucial for migratory NC commitment of melanocytic (*SOX10*, *PAX3*, *FOXD3*), neuronal (*FOXD3*), and glial (*SOX10*) lineages ^160–164^, as well as for the Schwann Cell Precursors (SCP) that are neural crest stem cells that reside on the axons of peripheral nerves until their late developing target organs, such as the adrenal medulla, gut, hair follicles, dental papilla, and parts of the craniofacial skeleton are formed and able to receive the neural crest, which was not possible earlier at the time the primary neural crest cells ended their migration ^4,165^. Next, analysis of the primary migratory neural crest cells from Day 12 (P0), showed that unlike at the premigratory stage, majority of the neural crest genes were now downregulated, except *MSX2* and *SNAI1*, which were still upregulated in the DiGeorge migratory neural crest cells as compared to the healthy controls (Fig. 8B). Similarly, in the passaged migrating neural crest cells (P1), both the stem cell (*LIN28A/B*) and majority of the neural crest genes (*TFAP2A/B/C*, *SOX9*, *LIN28B*, *TWIST*, *PAX3*, *NGFR*, *ERBB3*) were downregulated as compared to the controls. However, *MSX2* and *SNAI1* as well as *ETS1* and *DRAXIN*, which drives EMT and is normally downregulated in migratory crest cells, were upregulated in the patient cells further supporting the finding that NC cells from DGS patients express key NC genes in wrong proportions and at wrong times (Fig. 8C). Combined, our transcriptional analysis suggests that the neural crest cells from DGS patients were not properly specified at the premigratory stage despite the fact that they were able to migrate out from the neural ectoderm (Fig 8A-C).

**Figure 8.**
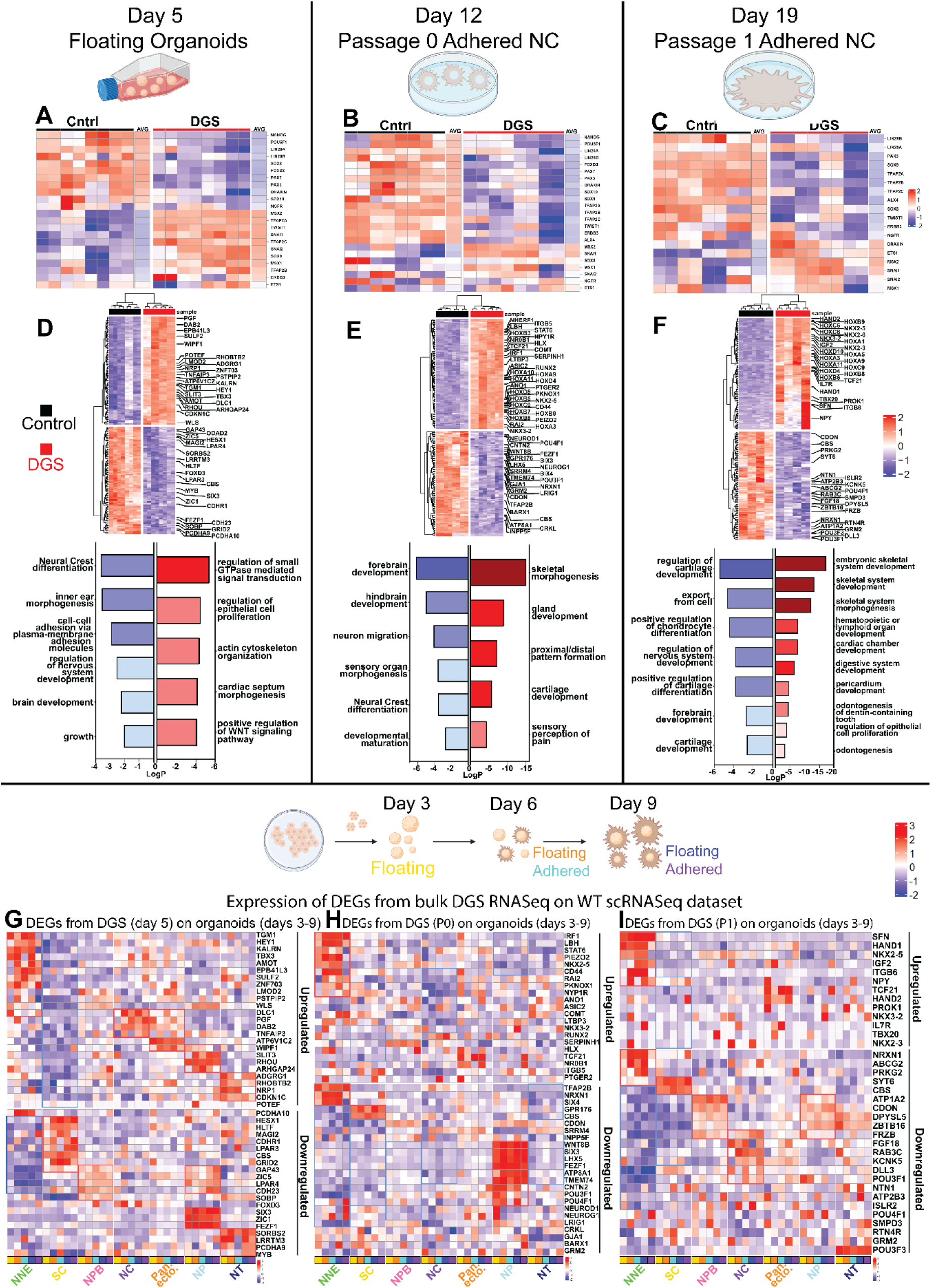
Transcriptional profiles of ectodermal organoids derived from DiGeorge patient cells show aberrant ectodermal pleistopotency, unbalanced NC specification, and a loss of cranial axial level phenotype. **A)** Heatmaps of bulk RNAseq data comparison of floating DGS patient ectodermal organoid gene expression on day 5 as compared to controls reflects aberrant NC induction and specification with downregulation of pluripotency genes and some NC specifier genes, while others, such as the epithelial to mesenchymal transition related genes SNAI1/2 and Twist are expressed prematurely. **B**) Transcriptional comparison of NC genes at P0 shows continued phenotype of aberrant specification in DGS patient NC cells at late primary migratory stage. **C**) Transcriptional comparison of NC genes at P0 shows continued phenotype of aberrant specification in passaged migratory (P1) DGS patient NC cells. **D**) Genes that constitute the highlighted Gene Ontology terms (selected from top 20) are shown on a heatmap of differentially expressed genes in day 5 floating ectodermal organoids. **E**) Genes that constitute the highlighted Gene Ontology terms (selected from top 20) are shown on a heatmap of differentially expressed genes in primary P0 migratory NC cells. **F**) Genes that constitute the highlighted Gene Ontology terms (selected from top 20) are shown on a heatmap of differentially expressed genes in P1 migratory NC cells. **G**) A heatmap showing the expression of the up and downregulated genes from highlighted GO-terms (in D) in floating day 5 DGS floated ectodermal organoids plotted on scRNaseq WT data from ectodermal organoids during normal development on Days 3-9. Upregulated genes are expressed in committed ectodermal domains and absent from ectodermal stem cells. Downregulated genes are expressed in ectodermal stem cells, the NPB and neural domains and absent from NNE. **H**) A heatmap showing the expression of the up and downregulated genes from highlighted GO-terms (in E) in passaged (P0) DGS migratory NC cells on day 12 plotted on scRNaseq WT data from ectodermal organoids during normal development on Days 3-9. Upregulated genes are expressed abundantly in NNE and downregulated genes in neural cells. **I)** A heatmap showing the expression of the up and downregulated genes from highlighted GO-terms (in F) in passaged (P1) DGS migratory NC cells plotted on scRNaseq WT data from ectodermal organoids during normal development on Days 3-9. Upregulated genes are expressed abundantly in NNE and downregulated genes in all ectodermal cell populations. (Heatmaps with full list of too 200 DEGs is shown in in Supp Fig 17). NNE: Non-Neural Ectoderm, SC: Stem Cells, NPB: Neural Plate Border, NC: Specified Neural Crest, Pan-ecto: Pan-ectodermal Progenitors, NP: Neural Plate, NT: Neural Tube.

To gain a broader understanding of the transcriptional changes in the DGS patient cells in the ectodermal organoids, we next plotted all the DE-genes from the DGS patient cells (in comparison to healthy control cells) on a heatmap and investigated their potential functional roles by creating Gene Ontology (GO) terms (Figs. 8D-F, Supp. Tables 5-10). In the floating organoids at Day 6, the top 20 downregulated terms associated with the differentially expressed genes included “Neural Crest Differentiation”, “regulation of nervous system development”, and cell-cell adhesion via plasma membrane adhesion molecules”, and the upregulated top 20 GO-terms consisted of regulation of “small GTPase mediated signal transduction” and “epithelial cell proliferation” as well as “actin cytoskeleton organization”, “cardiac septum morphogenesis” and “positive regulation of WNT signaling pathway”. All the genes from the twelve highlighted GO-terms are shown on the heatmap with a total of top 200 up and downregulated DEGs (Fig. 8D). At P0, in the samples that only consisted of the primary migratory neural crest cells, the GO-terms from the list of top 20 downregulated DE genes included similar terms like “neural crest differentiation”, “forebrain and hindbrain development”, and the upregulated top 20 terms contained “skeletal morphogenesis”, “gland formation”, “cartilage development”, and “proximal/distal pattern formation” (Fig. 8E). Finally, the P1 migratory neural crest cells of the DGS patients upregulated terms (within top 20) like “skeletal system development”, “digestive system development”, “cardiac chamber development”, “hematopoietic or lymphoid organ development”. Intriguingly, downregulated terms were related to “cartilage development”, “nervous system development” and “export from cell” (Fig. 8F). The genes contributing to the GO-terms are highlighted on the respective heatmaps of top 200 DEGs (Figs. 8D-F). Interestingly, the migratory neural crest cells (P1) from DGS patient cells showed upregulation of multiple posterior HOX-genes, indicating a shift from the cranial and vagal axial level identity into the transcriptional state of trunk neural crest cells. Plotting all the Hox-genes further verified the posteriorized phenotype (Supp. Fig 17A). Additionally, the patient cells displayed a downregulation of *FGF18*, which regulates both osteo- and chondrocytosis and FZB1, an antagonist of WNT signaling that positively influences the formation of mineralized matrix in the late stages of osteogenic differentiation of the calvarian bone and its expression is reported in the developing neural crest, neural tube and cephalic neurons, spinal ganglia, and cephalic, vertebral and limb cartilage in the chick embryo ^166–168^ (Fig. 8F).

Our results so far suggest the establishment of the pluripotent-like pleistopotent stem cell capacity of the NC cells (required to form the broad variety of derivative cell types later in embryogenesis) is defected, as shown by unbalanced transcriptional NC marker signature from early induction to late migratory cells. As compromised lineage specification is often accompanied with increased ectopic expression of genes from the neighboring domains, we next investigated which cell types during normal ectoderm development express the genes that were included in the top 200 of differentially expressed genes in the bulk analysis of DGS cells as compared to controls. For this, we first plotted the expression of all the up- and downregulated genes from the highlighted GO terms in the Day 6 floating organoids on the scRNAseq data set of all WT ectodermal subpopulations. In line with our hypothesis of the loss of pluripotency-like stem cell capacity in DGS, none of the upregulated genes were expressed in the ectodermal stem cell population and were only present in low levels in the neural plate border cells. Rather, the upregulated genes were expressed in the committed cell clusters of non-neural ectoderm, neural crest, and the neural populations, suggesting a premature loss of ectodermal and NC stemness. The upregulated neural crest population genes included *DLC1*, which was recently identified as a marker of neural crest cells that are in an intermediate state of EMT, in line with the premature expression of other EMT markers *SNAI1* and *TWIST1* ^169^ (Fig 8G).

On the other hand, the genes that were downregulated in the DiGeorge organoids, were expressed by the stem cells, the neural plate border, neural plate and tube populations (including known neural genes like (*ZIC1*, *FEZF1*, and *SIX3*) (Fig 8G). We also plotted all the top 200 up and down regulated DE-genes from the DGS bulkRNAseq data set onto the scRNAseq data set, which further strengthened the findings of the cell type specific patterns (Supp. Fig. 17B,C). It is important to note, that since the Day 6 bulk RNAseq data was collected from the organoids that contain all ectodermal cell types, unlike the P0 and P1 neural crest cell samples, we cannot distiunguish whether these results reflect changed expression levels of the genes in the stem-, neural plate border and neural crest cells, or do they reflect a proportional loss of cells in the stem and neural plate border clusters, and concomitantly, a gain of neural and non-neural cell types.

Next, we examined the P0 samples that consist of late, primary migratory neural crest cells. The results demonstrated two patters: several genes that were upregulated in in the DGS patient P0 neural crest were normally expressed by the non-neural ectoderm, and the downregulated genes were found to be expressed by neural progenitor cells of the neural plate cluster population (Fig. 8H and Supp. Fig. 17D,E). Finally, the pattern of P0 was similar to P1: upregulated genes in the DGS P1 migratory neural crest cells were genes that normally were expressed in the non-neural ectoderm, and also in the pan-ectodermal progenitors, whereas the upregulated genes represented all the ectodermal populations (Fig. 8I and Supp. Fig. 17F,G). Importantly, the upregulated genes of the DGS patient cells that were expressed highest in the non-neural ectoderm cluster of the WT organoids included genes like the homeobox gene NKX2.5, which plays a role in keratinocyte progenitor cell differentiation ^170^ but also in neural crest cell derivatives such as in pharyngeal arch patterning ^171^ and cardiac development of the neural crest derived outflow track and septa^172^. Nkx2.3 was also upregulated in the P0 migratory neural crest cells, which in turn regulates development of teeth and the identity of mucous acinar cells in the salivary glands ^173,174^ and function of various mesenchymal cell types such as homing of T-lymphocytes ^175^, hematopoietic stem cell self-renewal ^176^ and prevention of lymphatic endothelial differentiation^177^. Similarly, the transcription factor HAND2 is not *per se* associated with skin development but instead known for its role in the formation of the heart outflow trac ^178^, enteric and sympathetic nervous system ^179^ and ^180^, and adipocytes ^181^. Finally, the mechanosensitive ion channel PIEZO2 mediates pain and touch in the skin ^182^ but also regulates neural crest migration via interaction with semaphorins ^183^, and the development of the jaw bones ^184^. In sum, it should be acknowledged that although the expression of the upregulated genes found in the DGS migratory neural crest cells in our ectodermal organoids were expressed relatively the highest in the non-neural ectoderm population, we interpret their expression in the DGS patient neural crest cells to rather reflect an aberrant non-neural/mesenchymal transcriptional profile rather than an indication of epidermal lineage commitment.

## Discussion

Neurocristopathies comprise half of birth defects and several cancers and thus cause a significant clinical burden. *In vitro* disease modeling using CRISPR and patient-derived iPSCs has advanced rapidly, currently there are no models to study human ectodermal patterning, of which neural crest induction is a part of. Due to restricted access to embryos and ethical concerns, our knowledge on early cranial development between gastrulation and the end of neurulation is extremely limited. While technologies like microchips and microfluidics have improved neural tube modeling, they fail to fully replicate ectodermal patterning and are costly and labor-intensive ^185,186^, and these current models thus lack the depth needed for accurate study of neural crest development.

Our human ectodermal organoid model addresses these gaps as it recapitulates early embryonic development, showing interactions between future skin, CNS, and neural crest domains, along with minor mesodermal and endodermal contributions. Transcriptomic comparisons with chick neurulation stages ^7,78,187^ and ortholog-based analyses confirm high conservation across species, validating the model. Importantly, the migrating cells represent a pure population of *in vivo* -like pleistopotent neural crest cells capable of generating derivatives beyond the germ layer rule ^4^, This positions the organoids as a robust tool for modeling neurocristopathies and engineering neural crest-derived organs for regenerative medicine.

Ectodermal development depends on signals from surrounding tissues—a complexity mirrored in our organoid model. All major signaling pathways known from vertebrate embryos were active^188^, but our analysis revealed novel insights into both signal sources and recipients, often overlooked in studies. CellChat predictions highlighted key interactions: FGF2 from ectodermal stem cells induced cell types across the developing ectoderm; WNT4 originated from neural plate border while WNT6 was secreted by multiple ectodermal and mesodermal sources; BMP2/7 from endoderm influenced non-neural ectoderm more strongly than the traditionally recognized non-neural ectodermal BMP4. Unexpectedly, strong MidKine signaling also emerged, suggesting roles needing further study. Notably, stringent analysis cutoffs may have excluded some low-intensity but relevant signals. Importantly, the conserved signaling interactions detected in the human organoids also suggest that the cultures are self-sufficient in creating the endogenous crosstalk despite the added growth factors in the ectodermal organoid medium.

The remarkable, spontaneous recapitulation of colinear Hox gene activation over time in the organoids further validates the fidelity of the organoid model in mimicking *in vivo* axial organization and enables the generation of ectodermal derivatives from all axial levels. This includes generation of trunk neural crest cells for studies relevant to cancers like pheochromocytoma and the pediatric neuroblastoma, for which the requirement of creating trunk neural crest cells for achieving proper modeling is emerging in the field ^22,189^.

Pan-ectodermal progenitors in the human organoids are a newly identified, uncommitted cell population in early ectoderm, named after a similar “undecided” population few found in chick embryos, which co-express low levels of multiple lineage markers alongside pluripotency genes ^7^. Though transcriptionally quiet in signaling, RNA velocity places them as intermediates between stem cells and differentiated ectodermal domains. GO-term analysis also suggests active RNA processing, proliferation, and translation, indicating a potentially important yet unexplored role in ectodermal patterning.

DiGeorge syndrome is a multifaceted syndrome typically caused by a hemizygous 3Mb microdeletion that harbors 55 coding, 7 long non-coding RNA, and 7 micro-RNA genes ^36,190^. Although traditionally not classified as a neurocristopathy but rather thought to be caused by defects in organs from multiple origins, given that multiple neural crest derivatives are affected, DiGeorge strongly suggests a neural crest origin. Previous studies have focused on later developmental outcomes in DiGeorge syndrome, without addressing how early neural crest development is affected—the stage from which these structures originate. Our organoid model allowed us to pinpoint which of the early embryonic cell types and stages express genes of the DiGeorge region, narrowing potential candidates to a few genes active during ectodermal patterning. These likely contribute to the ectodermal stem cell and neural crest specification defect we observed at stages equivalent to weeks 3–4 of gestation. Notably, the top candidate genes identified (*RANBP1*, *DGCR6L*, *SLC25A1*, *MRPL40*, *SNAP29*, *THAP7*, *CDC45*, *HIC2*, and *DGCR8*) are unstudied in early neural crest or ectoderm development.

RANBP1, the most highly expressed DGS region gene in neural crest cells in our study, was recently shown to replicate DiGeorge midline facial anomalies when knocked out ^191^, and is essential for neural crest cell polarity and migration ^14^. It also regulates spindle assembly via RAN-GTP production ^148^. The mitochondrial protein MRPL40, another key candidate, showed peak— but not exclusive—expression in neural crest cells, and has been shown a functional role in neural progenitor proliferation in zebrafish ^192^. DGCR6L, expressed highest in early stem cells and neural crest, remains poorly understood although later in the embryo, downregulation leads to cardiovascular defects via TBX1/UFD1L regulation in branchial arches ^193^. CRKL, a mediator of RAS and JUN kinase and Yap signaling pathways, that is well known for oncogenic functions including proliferation and growth, migration, invasion, metastasis and EMT ^151,194^, was expressed across early ectoderm populations. CRKL is essential for differentiation of neural crest derived cell types in the branchial arches ^195^ and required for FGF8 responsiveness (Moon et al., 2006). We also detected CDC45 and THAP7 in neural plate border and neural crest cells, both involved in proliferation control in other cell types ^196,197^.

We observed a reduction in immunopositive cells expressing pluripotency and early neural crest markers, respectively, accompanied by downregulated RNA levels of key pluripotency genes, including *NANOG*, *OCT4*(POU5F1), and *LIN28A/B*. Furthermore, balanced expression of different early neural crest specifier genes was disrupted in the DGS patient organoids. These genes are expressed in undecided ectodermal stem cells, pan-ectodermal progenitors and neural crest stem cells. ^7,198^. Combined, the transcriptional changes and functional cellular data suggests improper NC specification, premature EMT, and insufficient establishment of growth-competent NC cells. Notably, many of the downregulated NC genes are also expressed in Schwann Cell Precursors (SCPs), the residing mid-gestational neural crest stem cell reservoir that gives rise to a wide range of NC derivatives that form later in the embryo ^165^. Impaired potential of the neural crest to form a proper pool of SCPs would further limit the contribution of NC cells during embryogenesis.

Finally, the importance of an early neural crest specification defect in the pathogenesis of DGS is supported by evidence from animal models. First, mutations in *TBX1*, *CRKL*, *LZTR1*, *HIRA*, *SCARF2*, *PI4KA* and others—genes of the DGS microdeletion region functionally associated with the disease—recapitulate DGS-like phenotypes, but only in homozygous deletion models ^190,199–201^. This suggest that a previous insult, or a pathological combination of multiple hemizygous DGS genes of the microdeleted region are required for disease formation (a combination of both alternatives is likely). Second, many of the genes located in the DGS microdeletion area are essential for the development of multiple cell types around the body, but the severe defects of DGS patients predominantly occur in the neural crest derived cells. For example, as reviewed in ^174^, TBX1 expression in the surface ectoderm, which is not neural crest derived, is crucial for ameloblast differentiation, as well as for the maintenance of hair follicles, but defects of enamel are rarely reported and hair phenotypes are not included in DGS patients ^202^. Similarly, in addition to the neural crest derived cardiac outflow tract and septation defects of DGS, TBX1-/- mice also show defects in the mesodermally derived subpulmonary myocardium, which is not part of DGS^203^. Also, homozygous loss of SNAP29 results in various symptoms of the CEDNIK syndrome, (Cerebral Dysgenesis, Neuropathy, Ichtyosis, and Keratoderma), but only facial anomalies are shared symptoms with DGS ^190^; Patients with biallelic mutations in the mitochondrial membrane transporter *SLC25A1* gene show facial dysmorphism but the main severe symptoms of combined D-2- and L-2-hydroxyglutaric aciduria, a rare neurometabolic disorder characterized by hypotonia, respiratory insufficiency, and severe neurodevelopmental dysfunction are not included in DGS. Finally, although loss of RANBP1 is strongly linked to both craniofacial defects and a thinner cortex, the specific CNS phenotype of altered cortex is not a part of DGS ^190^.

In sum, while additional work is required to determine detailed functional mechanisms, we hypothesize that the neural crest cells that show late phenotypic changes in multiple cranial tissues in DiGeorge syndrome have already gone through compromised neural crest specification much earlier in the embryo caused by the hemizygous deletion of the DGS critical region genes we identify here. We propose that this early vulnerability causing mechanism predisposes for the observed late phenotypes, which are further exacerbated by hemizygosity of additional DGS genes such as TBX1 and various other genes in the DGS microdeletion.

## Materials and Methods

All materials are listed in the STAR table below.

### Pluripotent Stem Cell Culture and cell lines

H1 and H9 human embryonic stem cell (hESC) lines (WA09 (H9) and WA01 (H1) from WiCell, were used for the characterization of the ectodermal organoids. For the DiGeorge syndrome study, induced pluripotent stem cell (hiPSC) lines from two patient and two controls from healthy relatives were used (https://hpscreg.eu/ control 1: GM - RCNSi003-A; control 2: F - RCNSi006-A; patient 1: M - RCNSi007-A; Patient 2: CH - RCNSi008-A and RCNSi008-B).

All Pluripotent cell lines were cultured on Matrigel coated plates in mTESR1 Basal Media (STEMCELL Technologies) and passaged with Accutase (Versene can also be used)) when ∼85% confluency had been reached. The cells were incubated at 37C in a humidified incubator at 5% CO2 and 20% O2.

### Ectodermal organoid cultures

The ectodermal organoid culture protocol was modified from ^20^ by doubling the original concentrations of Insulin, FGF, EGF. Human embryonic stem cell (hESC or iPSC) cultures at ∼80-90% confluence were dissociated using Collagenase IV at 37℃ for ∼ 45 minutes. Once the cells showed signs of colony retraction with the presence of small holes in the colonies, collagenase was discarded, and the cells were scraped using 5mL of DMEM-F12 with GlutaMAX, titurated a few times to form aggregates of 30-50 cells (smaller aggregates will hinder successful organoid formation) and transferred to ectodermal organoid media and cultured as floating aggregates in a T25 flask at 37°C in a humidified incubator at 5% CO2 and 20% O2. In the first 24 hours, the aggregates formed spheroids (and debri), which, starting at day 5-6 spontaneously adhered to the bottom of the flask. Neural crest cells started to migrate out and form large halos. For the generation of posterior neural crest cells, floating and lightly adhered spheroids were changed to new flasks in the same ectodermal organoid medium to generate further streams (streams #2 and #3). For imaging purposes of migratory neural crest cells together with the fused organoids in the center, floating organoids were changed to MatTek dishes prior to their adherence. For passaging of cranial migratory neural crest cells, the adhered cells were separated from the organoids that were lightly fused in the middle by brief incubation with accutase that allowed the organoids to detach and be removed, and only the migratory cells were further passaged into new flasks in the same ectodermal organoid media (passage 1). Alternatively, for imaging purposes, neural crest cells from the migratory mats were passages onto MatTek glass bottom dishes. Furthermore, we tested if the passages can be further expanded without them changing transcriptionally, and the results show high similarity of passages 1,2,and 3. The passaged cells can also be frozen and stored without them changing, as demonstrated by PCA plots from bulk RNAseq data (Supp. Fig. 3A).

### Neural crest derived cell type differentiation protocols

Passage 1 from cranial NCC was used as the starting point for the differentiation of the derivatives, except for the sympathetic neurons and the chromaffin cells, which were generated from Passage 1 of Stream #3 NCCs. The melanocytic protocol was started at the premigratory stage of cranial ectodermal organoids (see below). Since these culture protocols are long, we used bulk RNAseq to confirm that the Passaged neural crest cells can be frozen and thawed at a later time point, which was confirmed by a PCA plotshowing that the initial and thawed samples clustered together (Supp. Fig. 3A). We also compared transcriptional differences of sequenced cells from different passages. While P1 and a further passage P2 clustered together, the P0 that consisted of primary Day 12 migratory neural crest cells were slightly different but still more similar to the other passages than to the floating organoids and pluripotent cells (Supp. Fig. 3A).

### Adipocyte Differentiation

Neural crest cells were dissociated with Accutase and plated at 35,000-45,000 cells per cm^2^ density to 0.1 mg/mL poly-L-lysine coated wells in adipocyte media (α-MEM medium containing 10% FBS, supplemented with 1 uM dexamethasone, 10 μg/ml Insulin, and 0.5 mM IBMX) with media changes every 2–4 days for 21 days. The media was switched to a commercial adipocyte differentiation media (iXCells #MD-0005) with media changes every 2–4 days for another 12 days for further differentiation. Samples were stained with Hoechst and LipidSpot (Fisher Scientific #NC1669425) according to manufacturer’s instructions to stain for lipid droplets. Samples were also immunostained for UCP1 (Abcam #AB10983; 0.7:150).

#### Glial Differentiation

The differentiation of Schwann cells was adapted from ^24^. Neural crest cells were dissociated with Accutase and plated at 50,000 cells per cm^2^ density into poly-L-ornithine/laminin/fibronectin (PO/Lam/FN)-coated wells in DMEM/F12 medium supplemented with N2, 10 ng/ml of FGF2 and 10 ng/ml of EGF for 14 days for initial differentiation. After expansion and initial differentiation of cells in FGF2/EGF culture, cells are differentiated towards Schwann cell lineage in DMEM/F12 medium supplemented with N2, 10ng/ml ciliary neurotrophic factor, 20ng/ml neuregulin, 10ng/ml FGF2, and 0.5mM cyclic AMP for an additional 21 days. During this time the top cell layer is lost and cells differentiating into Schwann cells are formed. Media was changed every 2-4 days. Cells were collected at D14, D26/28 and D35. Cells were stained with S100β (Abcam #AB52642, 1:200, Sox10 (gift of Vivian Lee, previously at Medical College of Wisconsin, WI, 1:3000), and GFAP (MP Biomedical #0869110, 1:300).

### Osteocyte Differentiation

We adapted protocols for osteogenic and chondrogenic differentiation from ^25,204,205^. For osteogenic differentiation, hPSC–derived neural crest cells were plated onto poly-L-lysine (Merck, P5899) - coated glass coverslips (Stellar Scientific, 801007) in 24 well plates at 50,000 cells/well density in ectodermal organoid medium. The next day media was changed to osteogenic medium consisting of DMEM medium (Merck, M0450) containing 10% FBS (Merck, F4135), 10 mM D-glycerophosphate (Merck, G9422), 0.1 uM dexamethasone (Merck, D4902), 200 mM L-ascorbic acid 2-phosphaye (Merck, A8960) and 1% Pen-Strep. Media were changed every 2 to 3 days for 4 weeks. Cells were fixed in 4 % PFA and RNA was collected using Trizol Reagent (minimum of 6 wells/sample) at day 14 and day 28 of differentiation for immunostaining and Alizarin Red (Merck, TMS-008) staining purposes and for RNA isolation respectively. Of note, during the osteogenic cell culture maturation, a multiple-cell-layered structure emerges with mature cells located on the top with a stripe-like symmetrical morphology, and progenitor cell-like layers below (Suppl. Fig. 11). Occasionally, we noticed the multi-cell-layered structure had detached from the surface and formed calcified spheroids. However, after further culturing, the cells from the spheroids migrated out and repopulated the wells and reformed the layers (Suppl. Fig. 11).

#### Chondrocyte Differentiation

We adapted protocols for chondrogenic differentiation from ^25,204,205^. 50,000 hPSC–derived P1 neural crest cells/well were seeded into 96-well round bottom ULA plates (Greiner, 650185) in chondrocyte differentiation medium consisting of aMEM (Merck, M0450) supplemented with 10 % FBS (Merck, F4135), 10 ng/ml TGF-b3 (ThermoFisher, 100-36E-100UG), 0.1 mM dexamethasone (Merck, D4902), 200 mM L-ascorbic acid 2-phosphaye (Merck, A8960) and 1 % Pen-Strep. Media was changed every 2 to 3 days for 4 weeks. Chordospheres were fixed in 4 % PFA and RNA was collected using Trizol Reagent (minimum of 6 spheroids/sample) at day 14 and day 28 of differentiation for immunostaining and Alcian Blue (Merck, TMS-010) staining purposes. For RNA isolation, to improve RNA extraction from the chondrospheres, cold Trizol treatment was applied for 15 min while pipetting the sample.

#### Smooth muscle differentiation

Smooth muscle cell differentiation was based on optimization and a combination of two previously established protocols ^204,205^ 50,000 hPSC–derived P1 neural crest cells/well were plated onto poly-L-lysine (Merck, P5899) - coated glass coverslips (Stellar Scientific, 801007) in 24 well plates in ectodermal organoid medium. The next day media was changed to smooth muscle medium consisting of high glucose DMEM (Merck, D5671) supplemented with 1x GlutaMAX Supplement and 0.5 mM sodium pyruvate (ThermoFisher, 11360070), 10 % FBS (Merck, F4135), 10 ng/ml TGF-b1 (ThermoFisher, 100-21-100UG), optionally in some experiments 10 ng/ml PDGF-bb (ThermoFisher, 100-14B-100UG), 1% Pen-Strep. Media were changed every 2 to 3 days for 3 weeks. Of note, some established protocols used PDGF in their smooth muscle differentiation protocol. We tested the differentiation with and without and found no benefit but rather a worse outcome of its usage based on the biomarker analysis of the differentiating cells. Cells were fixed in 4 % PFA and RNA was collected using Trizol Reagent (minimum of 6 wells/sample) at day 10 and day 21 of differentiation for immunostaining purposes and for RNA isolation respectively.

### Sympathetic Neuron Differentiation

The sympathetic neuronal differentiation from the P1 neural crest stem cells was carry out using the previously published protocol ^206^ with a key modification: migratory neural crest cells were used instead of neural crest spheroids to initiate differentiation. Differentiation was performed in 24-well plates coated with poly-L-ornithine (PO), laminin (LAM), and fibronectin (FN). First the well plates were coated overnight with 15 µg/mL PO in 1× PBS. Then plates were then washed twice with 1× PBS and subsequently coated with 2 µg/mL LAM and FN in 1× PBS, followed by overnight incubation. The coating solution was aspirated just prior to cell seeding. The differentiation medium (per 100 mL) consisted of Neurobasal medium supplemented with 2 mL B27 (50×), 1 mL N2 supplement (100×), 2 mM L-glutamate, 200 µM ascorbic acid, 0.2 mM cyclic AMP (cAMP), and 25 ng/mL of glial derived neurotrophic factor (GDNF), 25 ng/mL brain-derived neurotrophic factor (BDNF), 25 ng/mL nerve growth factor (NGF). Freshly prepared 0.125 µM retinoic acid (RA) was added immediately before use. The medium was used within two weeks of preparation.

Neural crest cell suspensions were prepared in differentiation medium, and 120,000 cells were seeded per well. Cultures were maintained for 16 days with 80% medium changes every other day. Differentiation was assessed at day 10 (mid-point) and day 16 (endpoint) using markers specific to sympathetic neurons.

### Sensory Neuron Differentiation

The protocol for differentiating sensory neurons from the NC cells was adopted from ^207^ with a few modifications. Neural crest (NC) cell suspensions were prepared in sensory neuron differentiation medium comprising DMEM/F12 supplemented with 10% Knockout Serum Replacement (KSR), 1% penicillin-streptomycin, 0.3mM LDN-193189, 2mM A83-01, 6μM CHIR99021, 2mM RO4929097, and 3mM SU5402. Cells were seeded at a density of 120,000 cells per well in 24-well plates pre-coated with polyornithine, laminin, and fibronectin (PO/LAM/FN). All-trans retinoic acid (RA; Sigma-Aldrich, St. Louis, MO) was freshly prepared at a final concentration of 0.3μM and added to the differentiation medium. The medium was replaced by 80% every 2–3 days until day 16. Sensory neuron differentiation was assessed by evaluating the expression of sensory neuron-specific markers on days 10 and 16. No additional tests were performed for extended culture maintenance, in accordance with the original protocol^207^.

#### Chromaffin Cell Differentiation

The chromaffin cell differentiation was performed following a previously published protocol ^208^ with modifications using hESC-derived migratory P1 neural crest cells from the ectodermal organoids. Poly-L-Lysine (sigma P4707) coated (1.0mL/cm2), 24 well plates were used for chromaffin cell differentiation. The chromaffin cell differentiation medium was prepared by supplementing DMEM/F12 medium with 1xGlutaMAX, 2% B27, 0.5% BSA, 1x pen/step with 2.5 ng/mL recombinant human BMP4 (R&D Systems) with 50 uM of dexamethasone (Sigma-Aldrich) and 100 nM PMA (Millipore). The neural crest cells were seeded between 120,000-150,000 per well on a 24-well plate. The differentiation protocol was continued for 10 days and culture was tested for marker expression at days 5 (mid-point) and 10.

#### Melanocyte Differentiation

The melanocyte differentiation protocol was modified from ^28,209^. From day 3, the ectodermal organoids were cultured in their regular media supplemented with 600nM CHIR99021 (Wnt agonist). Of note, if the melanocyte differentiatiation protocol was started on migratory neural crest cells at P1, the cells uniformly differentiated into MITF-positive melanocytes, but the cells produced limited amounts of pigment. Once the organoids adhered, they were allowed to form a confluent mat of migratory neural crest. The melanocyte differentiation protocol was started once the migratory neural crest reached confluency, which was referred to as “Day 0”. On day 0 and day 1, cells were cultured with Essential 6 base medium (Gibco), 600nM CHIR99021, 10µM SB431542 (Selleckchem), and 1ng/mL BMP4 (R&D Systems). Soon after the addition of CHIR at day (6-8) all of the spheroids will adhere to the plate and the organoids flatten forming a monolayer of migratory neural crest cells. On days 2, 3, and 4, cells were fed with media consisting of Essential 6 base medium, 1.5µM CHIR99021, and 10µM SB431542. On days 5, 6, and 7, cells were fed with media consisting of Essential 6 base medium and 1.5µM CHIR99021. On days 8, 9, and 10, cells were fed with media consisting of Essential 6 base medium (Gibco), 1.5µM CHIR99021 (Biogems), 5ng/ml BMP4 (R&D Systems), 100ng/ml EDN3 (Sigma), and 50nM Dexamethasone. On day 11, cells were dissociated with TrypLE Express (Gibco), and FACS sorted using the following antibodies FITC-p75 (Cat 345104 from (BioLegend) and APC-cKIT Cat#17117942 (Invitrogen). The double positive cells were plated on poly-L-ornithine (PO)/Fibronectin/Laminin coated plates and fed with Melanocyte Maturation Media 1 (50% Neurobasal, 50% DMEM/F12 (Gibco), 1% B27 supplement, 1%N2, L-Glutamine, MEM NEAA, Pyruvate, Pen/Strep, 50nM Dexamethasone (Sigma), 100ng/ml EDN3, 0.8% ITS+ (Gibco), 5ng/mlBMP4, 50ng/ml SCF (Peprotech), 10ng/ml FGF2, 3µM CHIR99021, 100µM Ascorbic Acid (Sigma), 500µM cAMP, Dexamethasone (Sigma)). On day 19, the cells were dissociated with TrypLE Express (Gibco), and replated on poly-L-ornithine (PO)/Fibronectin/Laminin coated plates and feed with Melanocyte Maturation media (50% Neurobasal, 30% Low glucose DMEM/F12 (Gibco), 20% MCDB201 base medium (Sigma), 2% B27 supplement, 1% N2, L-glutamine, MEM NEAA, Pyruvate, Pen/Strep, 0.8% ITS+ (Gibco), 5ng/ml BMP4, 100ng/ml EDN3, 50ng/ml SCF (Peprotech), 4ng/ml FGF2, 3µM CHIR99021, 200µM Ascorbic Acid (Sigma), 500uM dbcAMP, 50nM Dexamethasone (Sigma)). Cells were incubated at 37°C with 5% CO2.

### Immunocytochemistry

*Floating organoids* were fixed in 4% paraformaldehyde for 10 minutes at room temperature and washed twice with 1X Phosphate Buffer Saline (PBS) and either immediately processed or stored in 1X PBS at 4C for future use. First, organoids were permeabilized with 3x PBST (Triton 0.5%) 30 minutes each at room temperature and blocked in PBST - 5% Bovine Serum Albumin for overnight at 4 °C on an agitator. Primary antibodies were diluted in the blocking buffer and incubated for two days at 4°C (with change to new antibody solution on Day 2) to ensure the antibodies can penetrate deeply into the organoids followed by five washes in PBST for 30 minutes each. The organoids were incubated with secondary antibodies that were diluted in blocking buffer (1:250) incubated at 4°C for at least 24 hours on an agitator and finally washed five times for 30 minutes each in PBST. For nuclear staining, the organoids were incubate in Hoechst solution (5ug/ml) for 1 hour at room temperature followed by two washes in PBST for 10 mins each. For mounting onto slides, SecureSeal Imaging Spacers were on slides. As much as possible of the PBST was removed following addition of 2 drops of the ProLong Glass Antifade mounting media onto the organoids which was applied in the middle of the spacers before the slides were coverslipped and sealed with nail polish.

Adhered organoids, and P0 and P1 migratory neural crest mats were cultured on MatTek glass bottom dishes to a confluency of ∼80%, fixed in 4% paraformaldehyde for 10 minutes at room temperature, and washed twice with 1x PBS. Samples were then either immediately processed or stored in 1X PBS at 4C for future use. Fixed cells were permeabilized with PBST for 30 minutes at room temperature. Cells were then treated with blocking solution (5% BSA in PBST) for one hour. Primary antibodies were diluted in blocking solution and added to the cells and incubate in 4°C for overnight followed by three times of five-minute washes in PBST. Secondary antibodies were diluted in blocking solution and incubated on cells for 1 hour at room temperature followed by three washes in PBST for 5 minutes each. For nuclear staining, Hoechst was diluted to PBST (5ug/ml) for 10 minutes at room temperature followed by two 5 minute washes in PBST and mounted with 2-3 drops of ProLong Glass Antifade mounting media. All antibodies are listed in the STAR table below.

### Melanocytes Immunocytochemistry

At the end of the melanocyte differentiation protocol, cells were seeded onto MatTek plates pre-coated with poly-L-ornithine (PO), fibronectin, and laminin. Cells were fixed with 4% paraformaldehyde (PFA) for 10 minutes at room temperature (RT), followed by three washes with phosphate-buffered saline (PBS). Permeabilization was performed using 0.5% PBST (PBS containing 0.5% Triton X-100) for 20 minutes at RT. After three additional PBS washes, cells were incubated overnight at 4 °C with primary antibodies diluted in blocking buffer (5% BSA in 0.5% PBST). The following day, samples were incubated for 2 hours at RT with secondary antibodies (1:1000) and Hoechst nuclear stain (1:2000). Finally, cells were washed three times with PBS and mounted using 2–3 drops of ProLong Gold Antifade and a coverslip.

### Antibodies

We used the following commercially available antibodies: anti-Mitf (1:100) GTX00738 GeneTex; anti-PMEL (1:100,) ab787 Abcam; Hoechst (1:2000 5ug/ml) Invitrogen.

### Sensory neurons, Sympathetic neurons and Chromaffin cells Immunocytochemistry

We use a general protocol for immunocytochemistry briefly Cells were fixed with 4% paraformaldehyde (PFA), permeabilized with 0.5% Triton X-100 in PBS for 30 min, blocked with 5%BSA in PBS for 1 h and incubated with the primary antibodies in 0.5% Triton X-100/5%BSA in PBS at 4°C overnight. 0.5% PBST incubated with the secondary antibodies (1:1000) and Hoechst nuclear stain (1:2000) (see Key resources table) at room temperature for 2h. Finally, cells were mounted using 2–3 drops of ProLong Gold Antifade and a coverslip.

### RNA Extraction and qPCR

Total RNA was isolated using NucleoSpin RNA Plus kit (Macherey-Nagel, 740984), or Direct-zol RNA miniprep plus kit (Zimo R2072) following manufacturer’s instructions. The cDNA synthesis was performed using iScript cDNA synthesis kit (Bio-Rad, 1708891). qRT-PCR was performed on QuantStudio 3 Real-Time PCR System (Thermo Fisher Scientific) using TaqMan gene expression master mix (Applied Biosystems 4369016). Relative mRNA expression levels were normalized to housekeeping gene expression by the ΔΔCT method. Primer sequences are listed.

### Bulk RNA Sequencing and sample preparation

Bulk RNA Samples were prepared in triplicates and sent to Azenta Life Sciences for sequencing. For wild-type human ectodermal organoid development, the following samples were collected for sequencing in two-day increments: H1 ESC reference, Day 2, Day 4, Day 6, Day 8, Day 10 floating organoids and Day 10 (adhered organoids and migratory neural crest). For neural crest derivatives, the following samples were sequenced: H1 ESC, Passage 1 migratory neural crest cells from Stream 1 cranial and Stream 3 trunk, respectively, sensory neurons at Day 10 and 16, sympathetic neurons at Day 10 and 16, chromaffin cells at Day 5 and 10, chondrocytes at Day 14 and 28, osteocytes at Day 14 and 28, smooth muscle cells at Day 11 and 21, adipocytes at Day 11, 21, and 33, glial cells at Day 14, 26, 28, 30 and 35. Sympathetic neurons with retinoic acid treatment at Day 2, 4, and 6 post-treatment were also sequenced. For the melanocyte differentiation protocol, migratory neural crest cells treated with CHIR99021 were sequenced at early, mid, and late differentiation timepoints (Day 0, 15, 35). DiGeorge patient and healthy relative controls were sequenced from Days 0 (iPSC), 5 (premigratory floating organoids), 12 (Passage 0, primary migratory NCCs), and 19 (Passage 1, migratory NCCs).A reference sample from a large quantity of a single isolation of H1 ESC RNA was sequenced as part of every sequencing set, which was used for correction of batch variances.

### Single-Cell RNA sequencing

Cell samples for scRNA-seq were dissociated using the Multi Tissue Dissociation Kit 3 (Miltenyi Biotec, 130-110-204). Briefly, spheroids or the migratory mat were rinsed with PBS, followed by the addition of the enzyme mix provided in the kit. Samples were incubated at 37°C for 10 minutes, then gently triturated by pipetting. This was followed by a second 10-minute incubation at 37°C and additional pipetting to obtain a single-cell suspension. The resulting suspensions were washed twice with cold PBS and filtered through a 70 μm MACS SmartStrainer (Miltenyi Biotec) to remove cell clumps. Single-cell capture and library preparation were carried out at the NIDCR Genomics and Computational Biology Core using the Chromium Single Cell 3′ Library & Gel Bead Kit v3 (10x Genomics), according to the manufacturer’s instructions. Pooled libraries were sequenced on an Illumina NextSeq 500 platform. Raw sequencing data were processed using the Cell Ranger Single-Cell Software Suite (10x Genomics) for demultiplexing, barcode assignment, and unique molecular identifier (UMI) quantification. Reads were aligned to the human reference genome (GRCh38-2020-A) that is publicly available on the 10X website.

### Imaging and Quantification

The Zeiss LSM 880 with AiryScan was used for confocal images. Images were quantified using ImageJ, for which z-stacks of organoids and differentiated cells were segmented using the Huang threshold and Region of Interest (ROI) manager to generate masks that were overlayed on all channels. For single cell level quantification of percentage of positive nuclei from the total number of nuclei. For bulk immunofluorescence intensity measurements, Raw Integrated Density (RawIntDen)/Intensity values were measured for each channel and were normalized to Hoecst staining intensity per area. The area values of chondrospheroids were measured in *μM*^2^.

For Melanocytes confocal images were captured with the Zeiss LSM 880 Images were quantified using ImageJ. Nuclei were segmented using the Huang threshold to generate a binary mask which was then subjected to watershed that was overlayed on the channels of interest. Then we use the Analyze particles tool to count the number of positive nuclei for each channel. To have the percentage of positive nuclei in each channel we divided the number of positive nuclei in each channel by the total number nuclei in using the DAPI channel.

### Bulk RNA Sequencing Analysis

Counts were filtered to remove all genes which did not have more than 5 reads in at least 3 of the samples. To ensure that there was no batch effect we performed a PCA and observed that the reference samples were clustering together. Downstream analysis and DGE analysis was performed using DESeq2 ^210^. Gene Ontology (GO) analysis was run using Metascape ^211^; (https://metascape.org/) on genes that were found to be significantly regulated (padj < 0.05, and LogFC > 1 or < -1). The list of all GO terms generated from DiGeorge patient and control cells at the various stages are provided in supplemental tables 5-10.

### Single-Cell Sequencing Analysis

FASTQ files were processed using 10X CELLRANGER v 5.0 and aligned to the human reference genome (GRCh38-2020-A) that is publicly available on the 10X website. CELLRANGER output files were then further analyzed using R/RStudio with the Seurat version 3.0.1 package ^212^. To limit the inclusion of poor-quality cells (e.g., dead cells, doublets) cell with mito-genes expressed higher than 90% were excluded, and a filter for nFeature (genes) of more than 200 and less than 5000 was used. Normalization and scaling of the dataset were performed using ‘LogNormalization’ and ‘ScaleData’ functions as following the standard Seurat workflow and vignette. The datasets were integrated using the Harmony package ^213^. Clustering was generated by using the parameters of resolution = 0.2, and dimensions = 1:20. Cell communication analysis was performed using the ligand-receptor based CellChat v2 ^61^ package by predicting communication probability between cell types using the ‘truncatedMean’ method with a trim = 0.1. Clustered cells were visualized via UMAP plot and manually annotated based on differentially expressed genes in each cluster (FindAllMarkers function, logfc.threshold = 0.2, min.pct = 0.25). The clusters that were identified as “Ectoderm” were extracted and re-clustered using the ‘subset’ function.

The Chick and Human integrated dataset was generated using the R Package OrthoIntegrate ^41^ following the workflow and vignette. The human and chicken ortholog gene list was generated with help from Mariano Ruz Jurado, provided as supplemental table 2. The integrated dataset was further processed and analyzed using Seurat in a similar fashion as mentioned above.

Neural crest organoid stream analysis (Fig. 5 E-G) was performed using Seurat 5.1v package (Hao et al. 2024). Streams 1 and 3 were integrated using the ‘sctransform’ function as per the Seurat vignette. Following integration, the overlap between stream 1 and stream 3 was not aligning precisely. To ensure that the clustering from stream 3 was still following the same clustering as stream 1 we generated Developmental Gene Modules (DGM) from stream 1 (Suppl. Table 3) using the DGM classification tool ^214^ and verified the expression of these DGMs on the respective stream 3 clusters (Suppl. Fig. 9 A).

The Cellranger output files were also processed using Velocyto ^58^ to generate loom files. Velocyto loom files were then further analyzed using Seurat and clustered in the same fashion as mentioned above. To follow gene regulatory networks controlling development clustering was performed using transcription factors expressed in the dataset. Full list of transcription factors expressed in human were generated from KEGGBRITE. Seurat clusters were converted to hda5 files in Seurat for use on Python. Dynamical RNA velocity construction was done on Python using the package scVelo ^215^. Pseudo-temporal analysis was performed using Monocle3 ^60^.

## Author Contributions

LK conceived the study. SuP, TB, ET, JH, ZZ, KB and SrP designed the experiments together with LK. SuP created the organoid datasets for bulk and scRNAseq analysis. JH, SuP, and KB optimized the ectodermal organoid protocol. ET analyzed the scRNAseq data with LK with contribution from SuP, CP, JH, and DM. Bulk RNAseq analysis was done by SuP, ET, SrP, and JH with LK. DM performed initial quality analysis and genome alignment of RNAseq samples. JH, SuP, KB and SrP performed immunostaining, HCR-FISH and quantitative image analysis of ectodermal organoids. TB prepared all DGS wet lab experiments and analyzed the DGS cell biology assays. AA provided the DGS cells and support on their maintenance. The neural crest derivative culture optimizations, RNA extraction, QPCR and quantitative immunostaining and related data analyses were performed by (ZZ, 3 lineages), TB (3 lineages), SuP (2 lineages) and KB and SrP, (1 lineage). ET prepared the figures. LK wrote the manuscript with input from ET, SrP and all authors. All authors wrote M&M of their work. LK supervised the study.

## Acknowledgments

This research was supported by the NIDCR Imaging Core: ZIC DE000750-01, and NIDCR Genomics and Computational Biology Core: ZIC DC000086.

## Disclaimer

This research was supported by the Intramural Research Program of the National Institutes of Health (NIH). The contributions of the NIH authors were made as part of their official duties as NIH federal employees, are in compliance with agency policy requirements, and are considered works of the United States Government. However, the findings and conclusions presented in this paper are those of the authors and do not necessarily reflect the views of the NIH or the U.S. Department of Health and Human Services.

## STAR Methods

### Cells

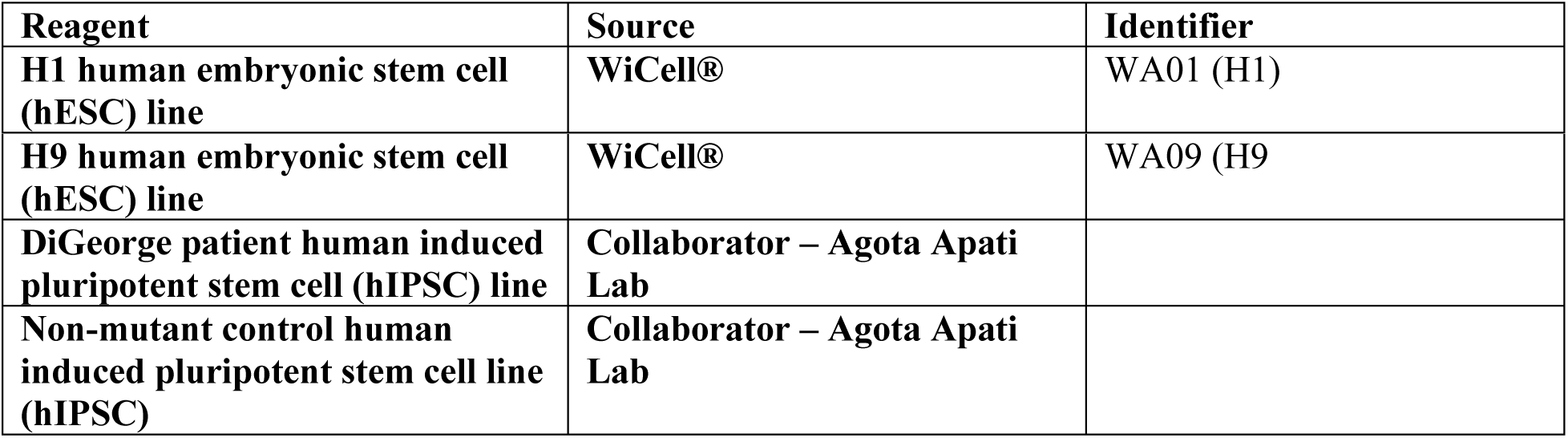

| Reagent | Source | Identifier |
| --- | --- | --- |
| H1 human embryonic stem cell (hESC) line | WiCell® | WA01 (H1) |
| **H9 human embryonic stem cell (hESC) line** | **WiCell®** | **WA09 (H9** |
| **DiGeorge patient human induced pluripotent stem cell (hiPSC) line** | **Collaborator – Agota Apati Lab** |  |
| **Non-mutant control human induced pluripotent stem cell line (hiPSC)** | **Collaborator – Agota Apati Lab** |  |

### ESC Cell Culture Reagents

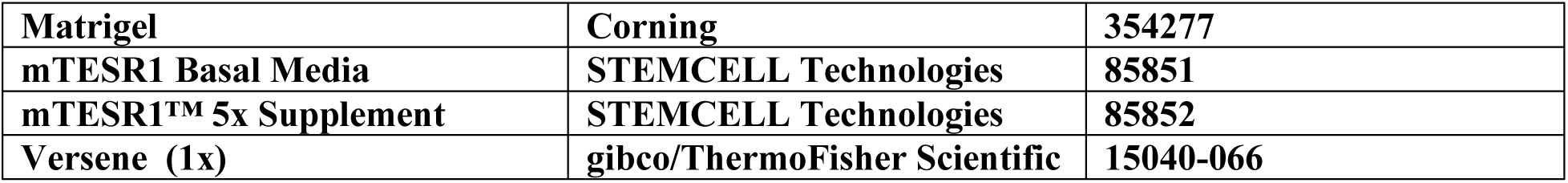

|  |  |  |
| --- | --- | --- |
| **Matrigel** | **Corning** | **354277** |
| **mTESR1 Basal Media** | **STEMCELL Technologies** | **85851** |
| **mTESR1™ 5x Supplement** | **STEMCELL Technologies** | **85852** |
| **Versene (1x)** | **gibco/ThermoFisher Scientific** | **15040-066** |

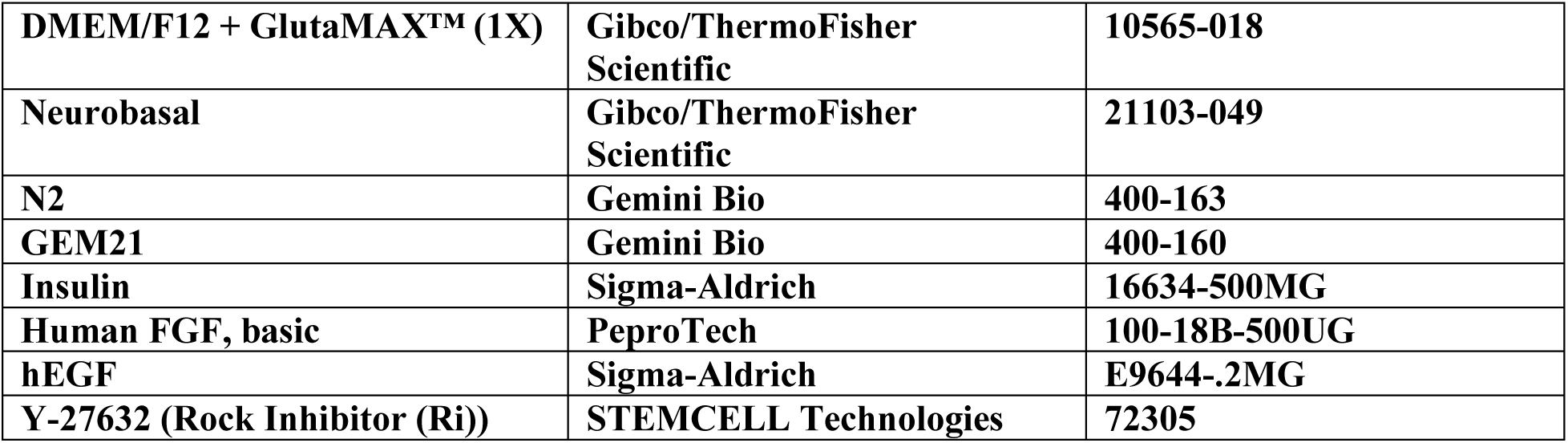

Neural Crest Media
|  |  |  |
| --- | --- | --- |
| **DMEM/F12 + GlutaMAX™ (1X)** | **Gibco/ThermoFisher Scientific** | **10565-018** |
| **Neurobasal** | **Gibco/ThermoFisher Scientific** | **21103-049** |
| **N2** | **Gemini Bio** | **400-163** |
| **GEM21** | **Gemini Bio** | **400-160** |
| **Insulin** | **Sigma-Aldrich** | **16634-500MG** |
| **Human FGF, basic** | **PeproTech** | **100-18B-500UG** |
| **hEGF** | **Sigma-Aldrich** | **E9644-.2MG** |
| **Y-27632 (Rock Inhibitor (Ri))** | **STEMCELL Technologies** | **72305** |

### Dissociation Reagents

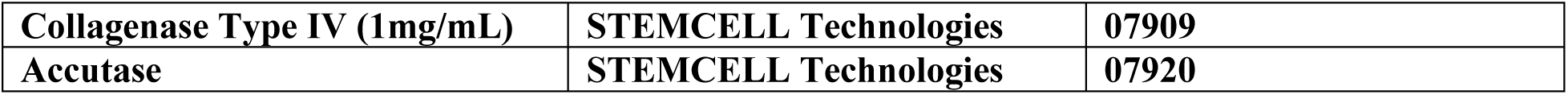

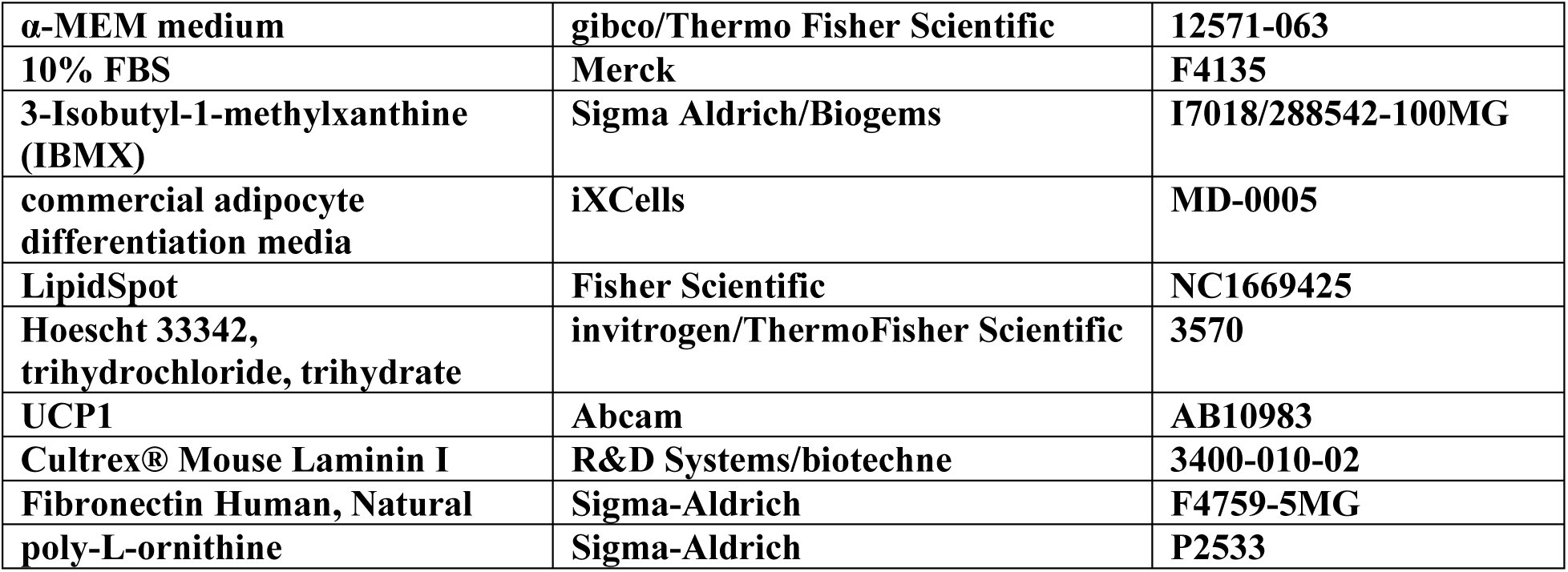
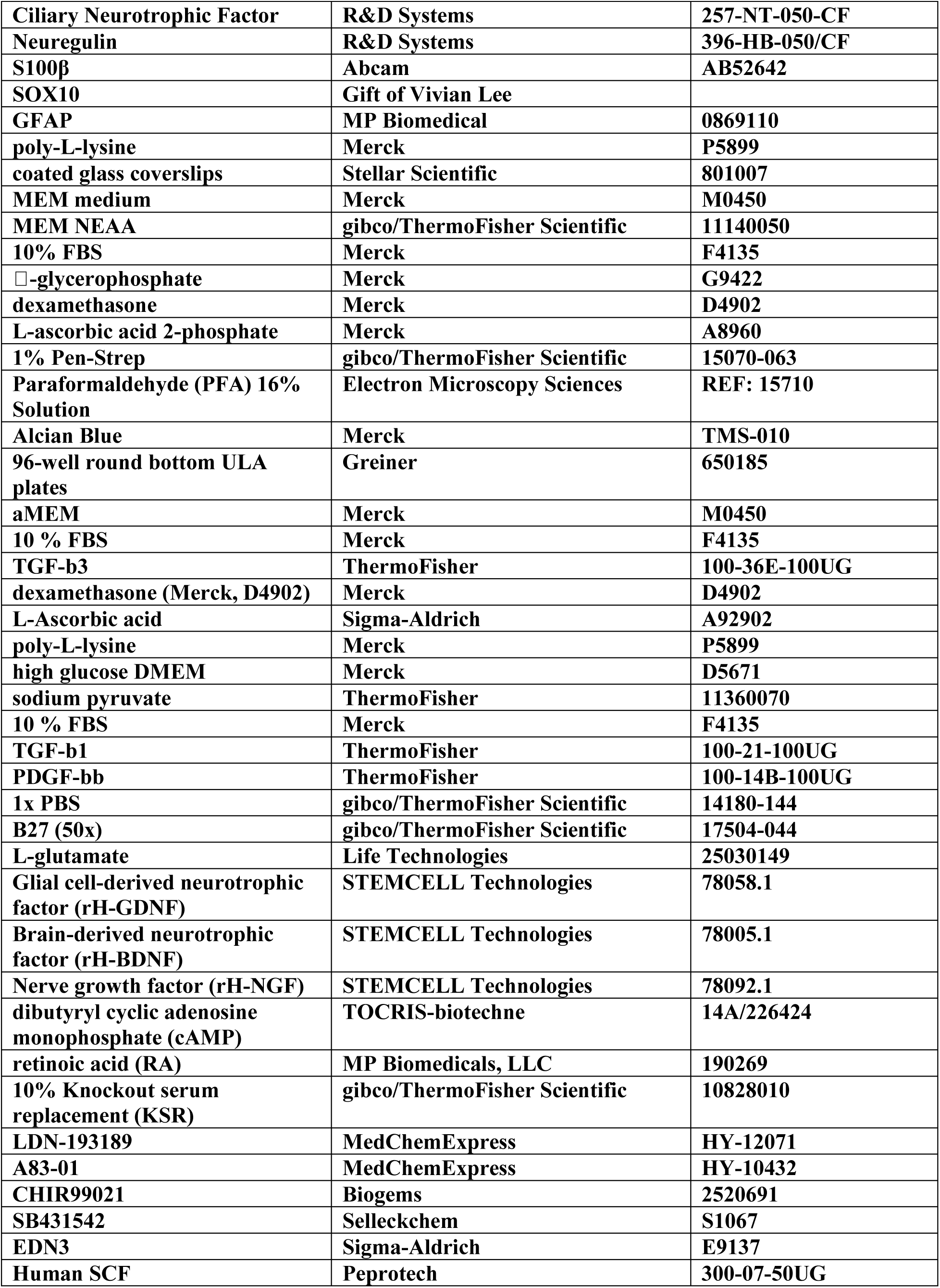
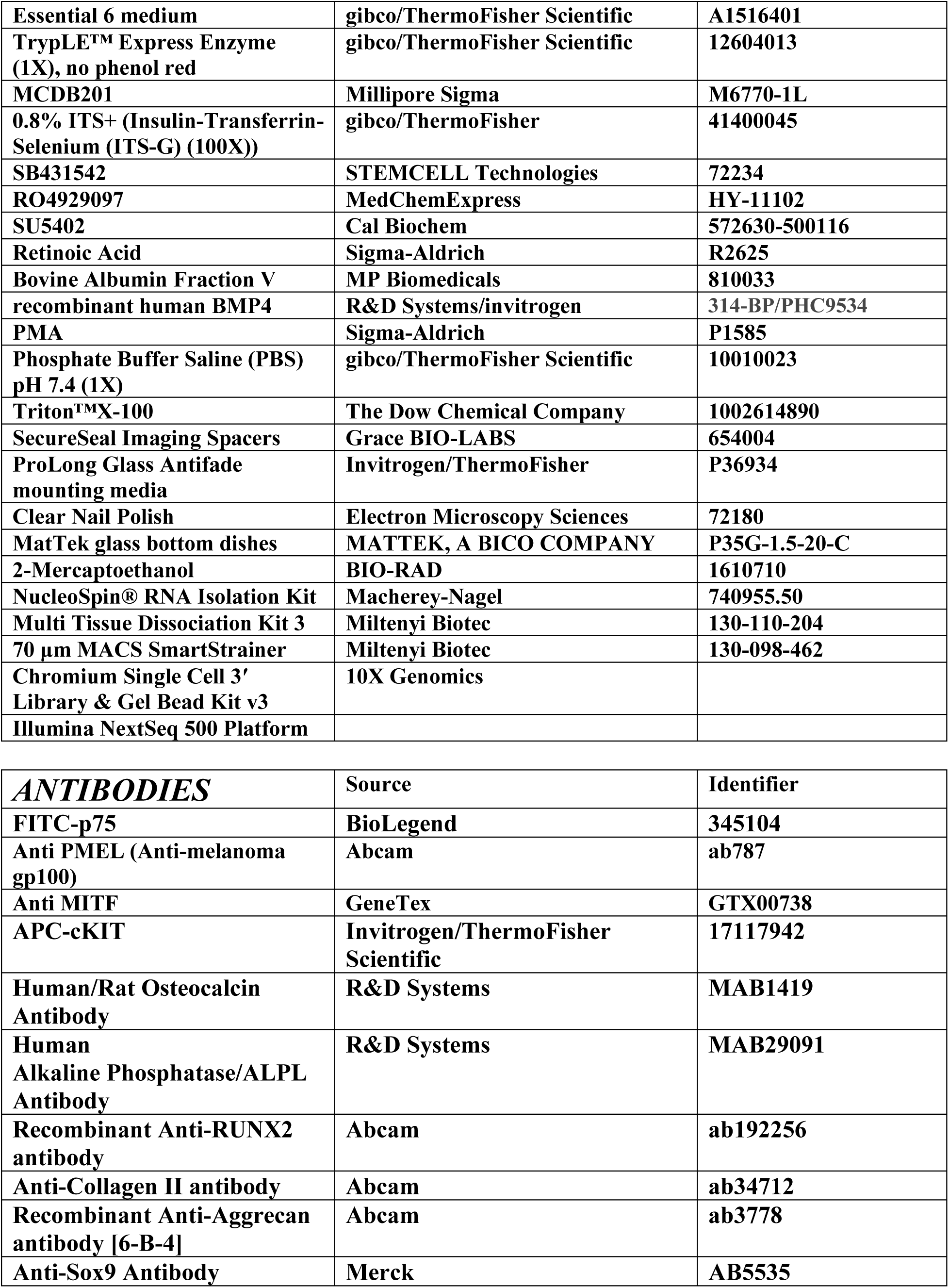
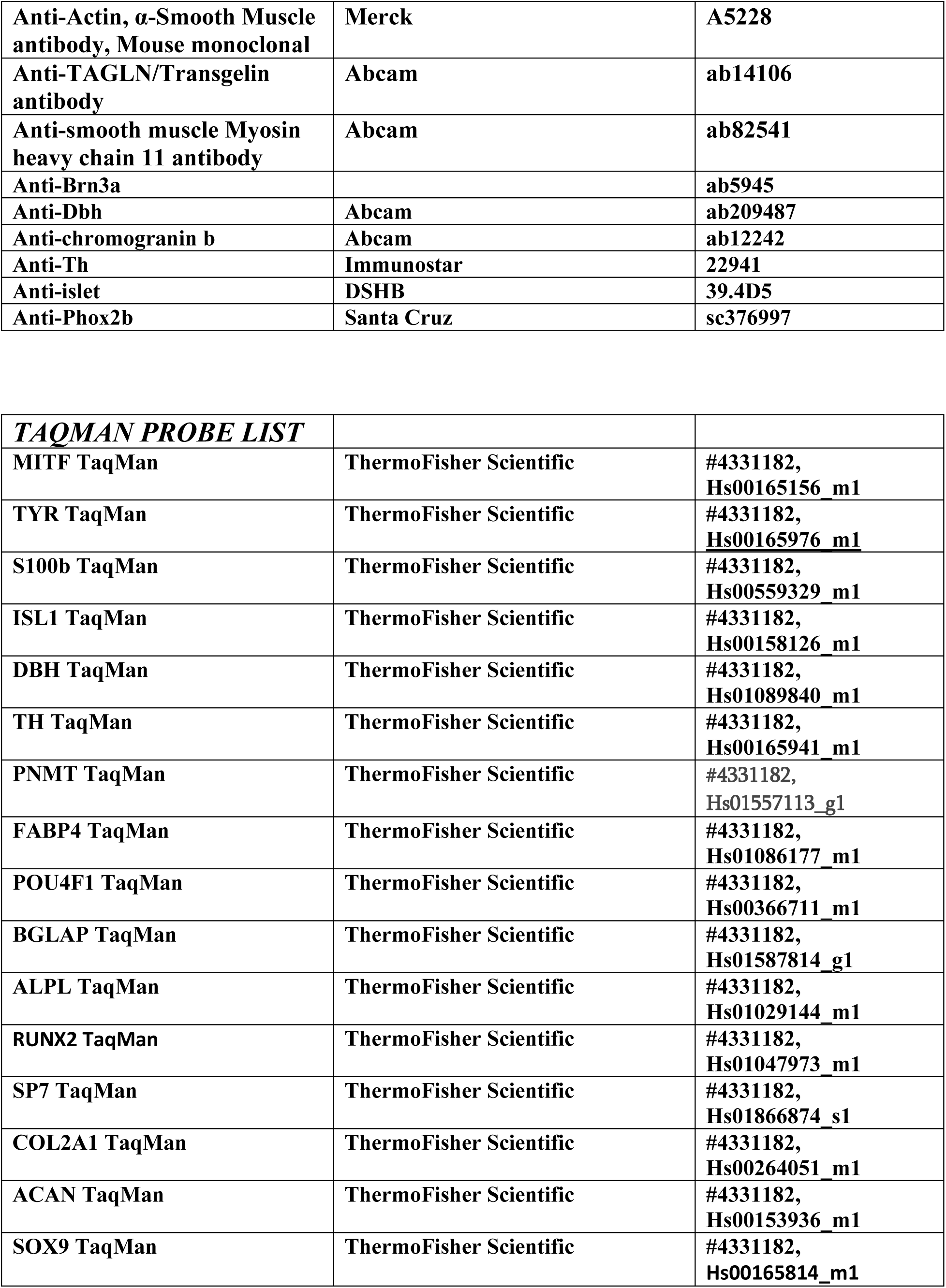
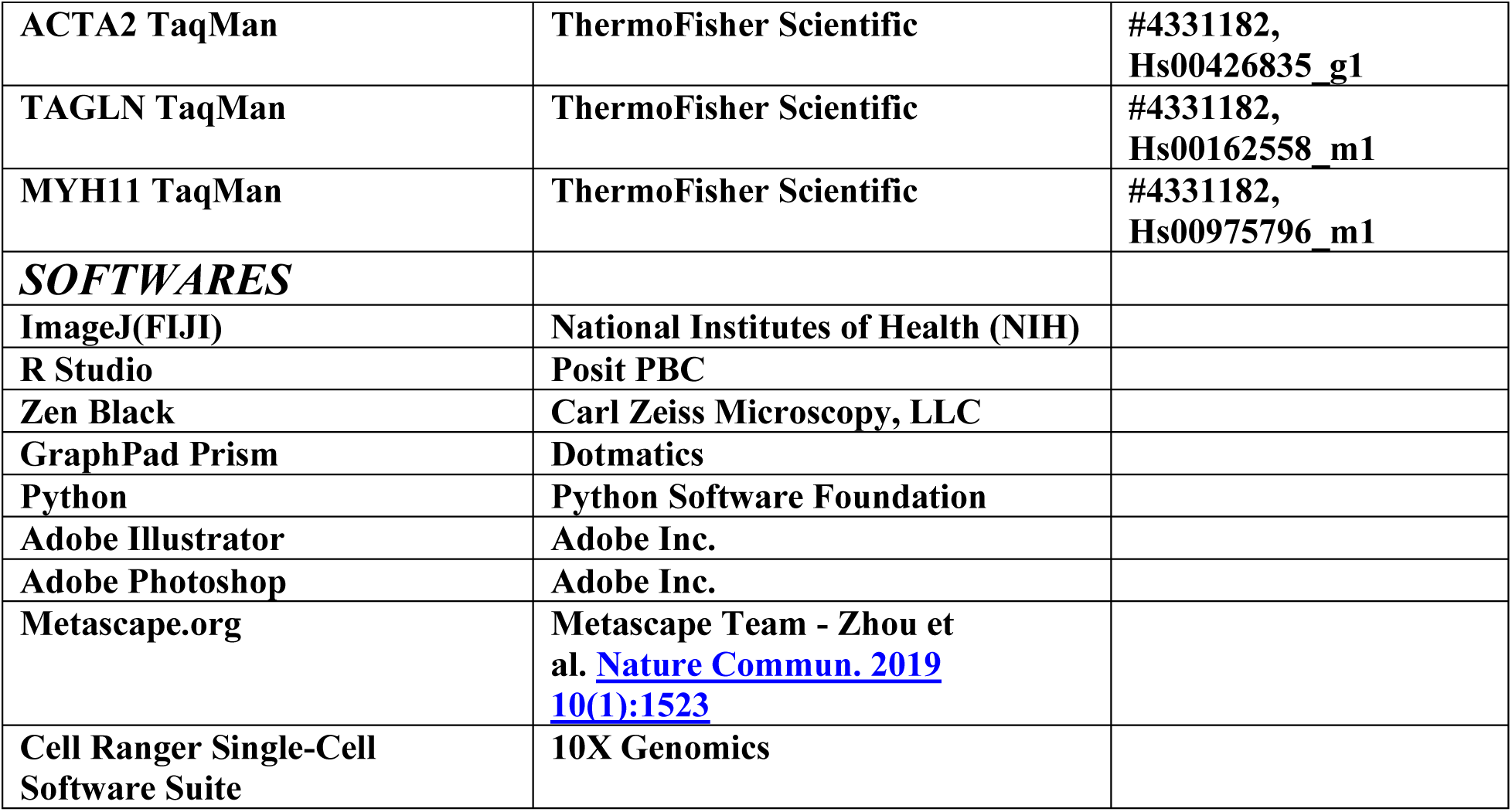

---

**Supp. Fig. 1.**
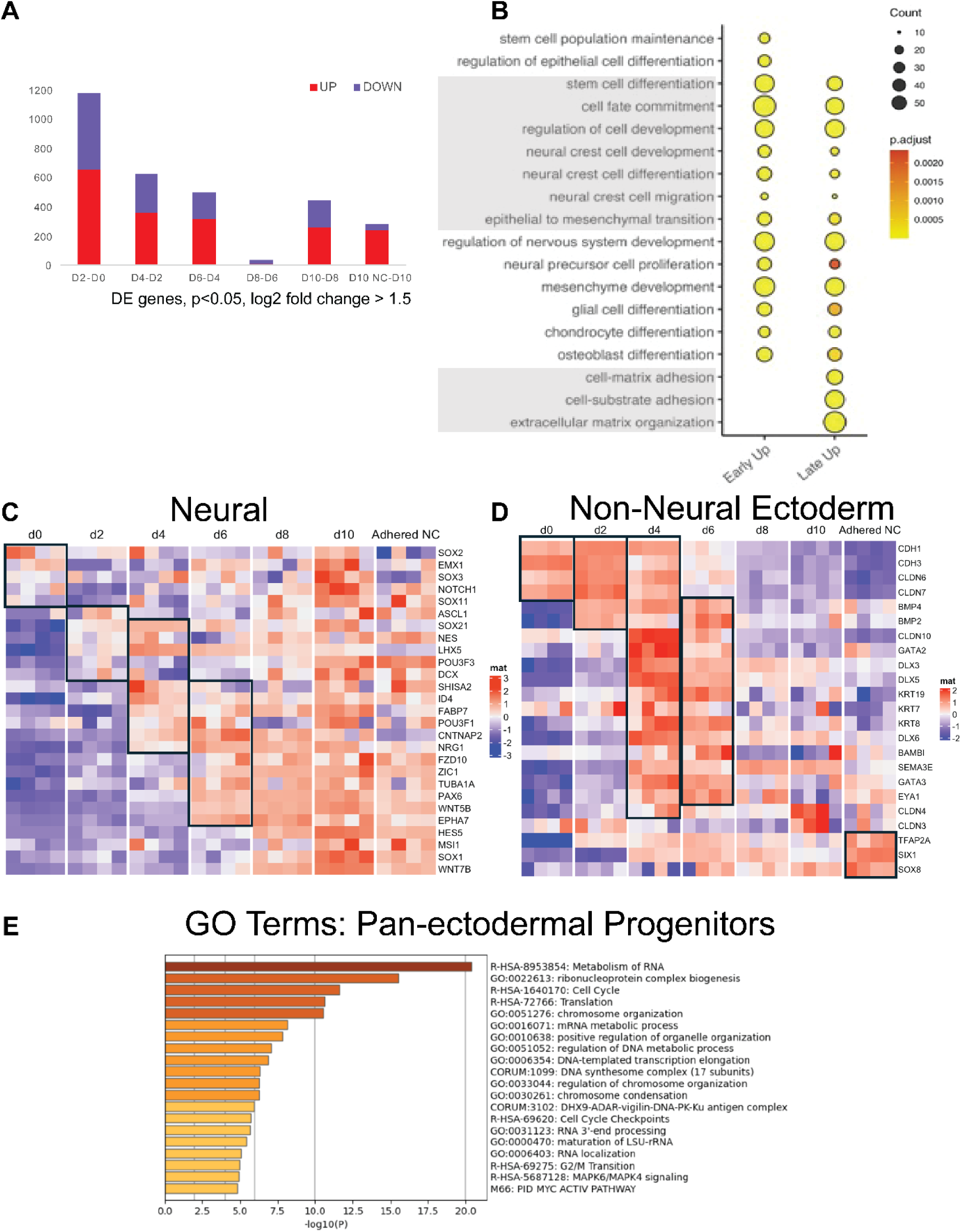
Bulk and scRNAseq analysis confirm transcriptional homology of ectodermal organoids to the ectodermal patterning process of avian embryos. **A)** Bar graphs show the amount of differentially expressed genes between developmental days in ectodermal organoids in the Bulk RNAseq data set. The transition of embryonic stem cells to ectodermal organoids shows the biggest change, and floating organoids between days 6 and 8 change the least. **B**) Gene Ontology terms from bulk RNAseq data set pooled to early (days 2-4) and late (days 6-10) stages shows cellular functions related to stem cell maintenance and fate commitment as well as neural crest formation emphasized in the early stages. Processes linked to neural, glial, and mesenchymal cell type differentiation and extracellular matrix production and organization are emphasized at the later stages of ectodermal organoid development. **C**) A heatmap shows that genes reflecting neural development of the central nervous system are expressed in a time dependent manner like what is known from the embryo. **D**) A heatmap shows that genes reflecting non-neural, future epidermal commitment are expressed in a time dependent manner in like what is known from the embryo. **E**) Top 20 Gene Ontology terms from scRNAseq data of the subpopulation of Pan-Ectodermal Progenitors highlights processes involved in RNA processing and metabolism, cell cycle regulation and chromosome condensation.

**Supp. Fig. 2.**
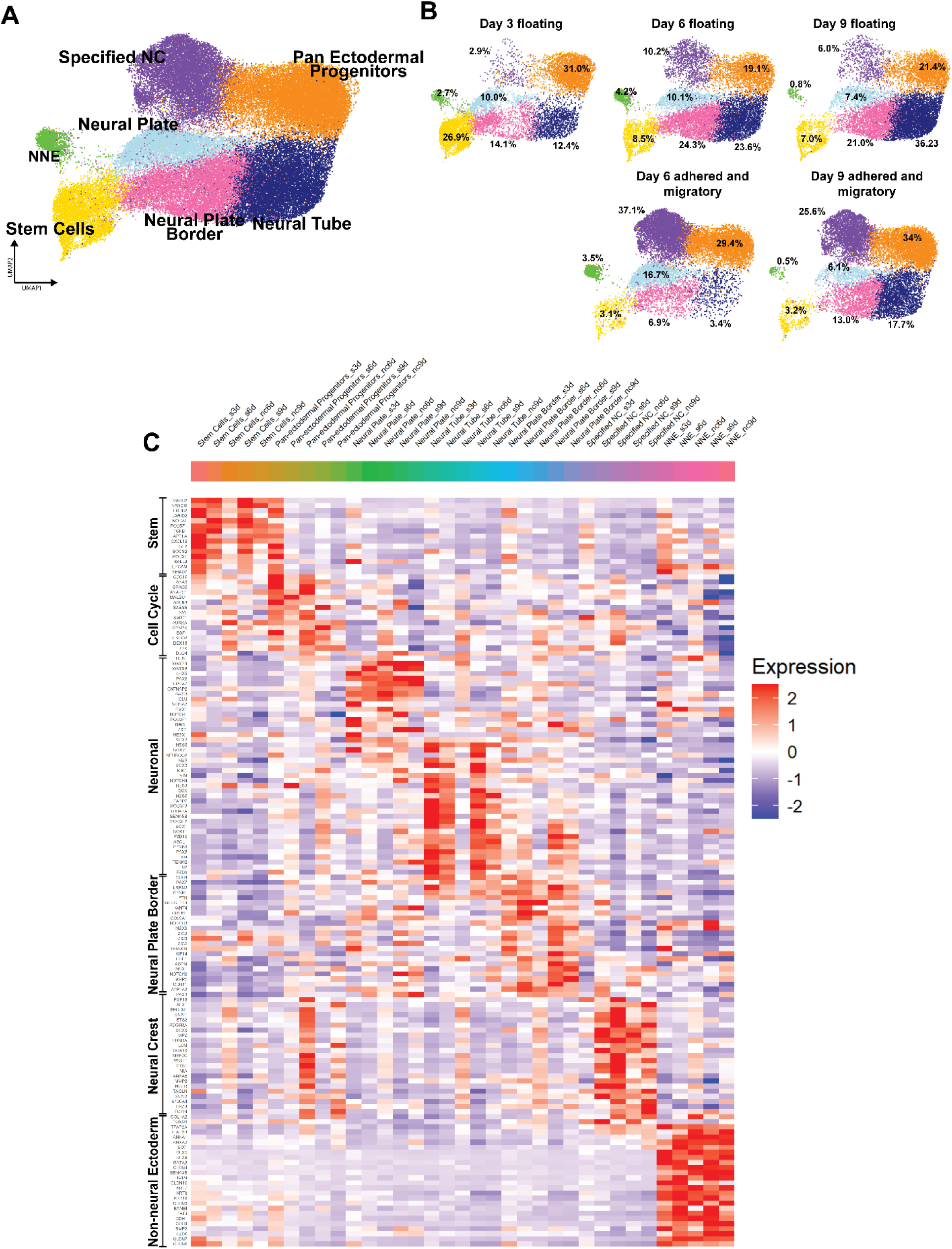
Ectodermal organoids are actively ongoing the whole process of ectodermal patterning - they consist of cells from ectodermal stem cells to immature and fully specified neural crest, future CNS and skin. **A)** UMAP clusters of ectodermal subsets of the ectodermal organoids pooled from all developmental stages. **B)** Subsets of ectodermal populations shown per sample and stage. **C)** Expression of differentially expressed genes (in the same sequence as for the pooled data in Fig2H).

**Supp. Fig. 3.**
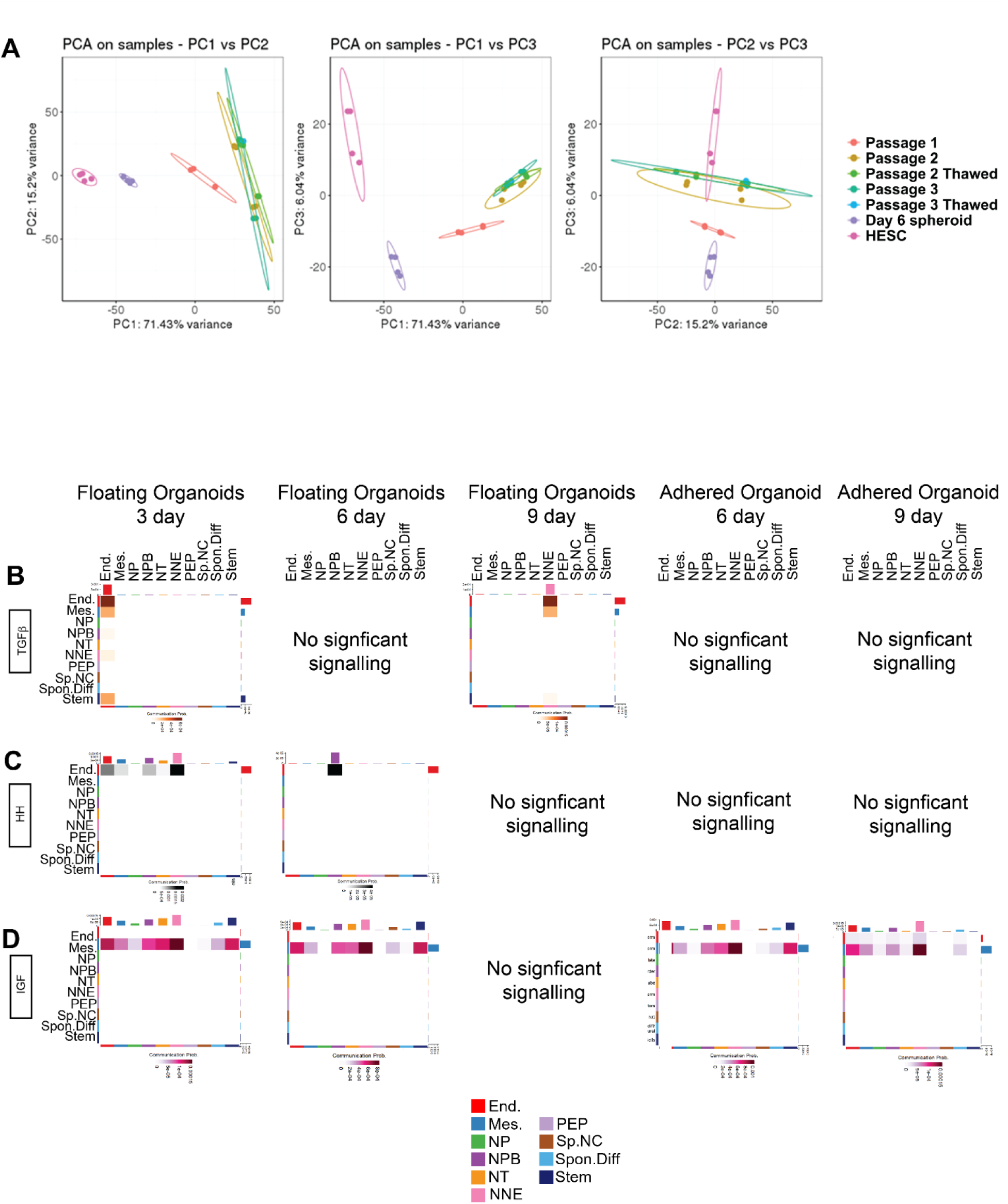
Cell-to-cell signaling of the ectodermal organoids reveal highly conserved mechanisms and novel interaction details. **A)** PCA plots demonstrate that passage 1 of migratory neural crest cells is similar to passages 2 and 3, and that cells retain their transcriptional profile after a freezing and thawing cycle. **B**) Cell Chat signaling analysis demonstrates minimal input of TGF-beta signalling on ectodermal development; mesodermal and endodermal input to non-neural ectoderm (NNE) is detected in floating organoids at Day 9. **C**) Endodermally produced Hedhehog signaling is received by neural plate border (NPB) and NNE in floating ectodermal organoids on days 3 and 6. **D**) Mesodermal IGF is received by the NNE, NPB, Neural Plate (NP) and ectodermal stem cell populations.

**Supp. Fig. 4.**
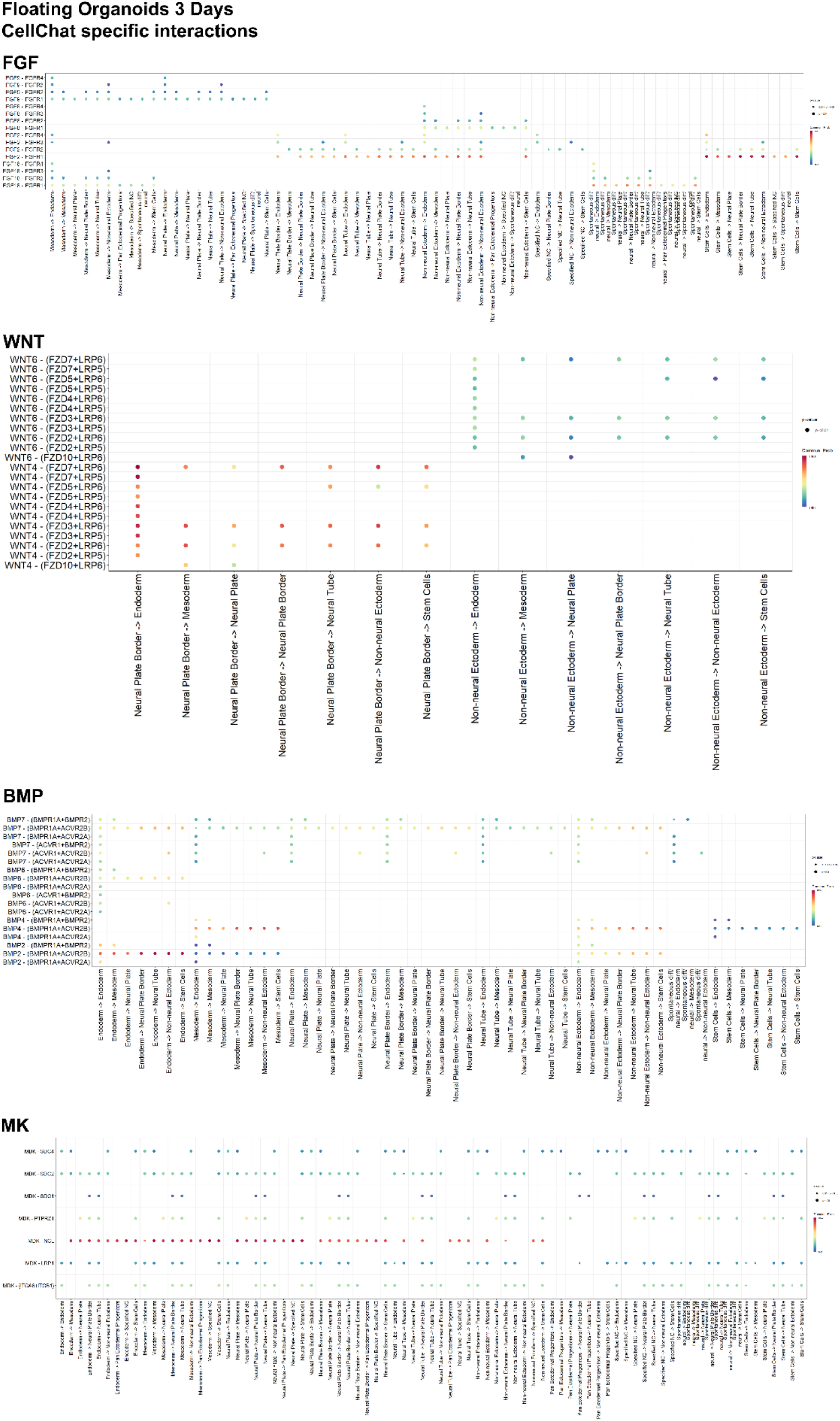
Cell-to-cell signaling of the ectodermal organoids reveal highly conserved mechanisms and novel interaction details. **A)** Detailed ligand-receptor pairs and interaction strengths at day 3 of the ectodermal organoids for FGF B) WNT C) BMP and D) Midkine signalling pathways.

**Supp. Fig. 5.**
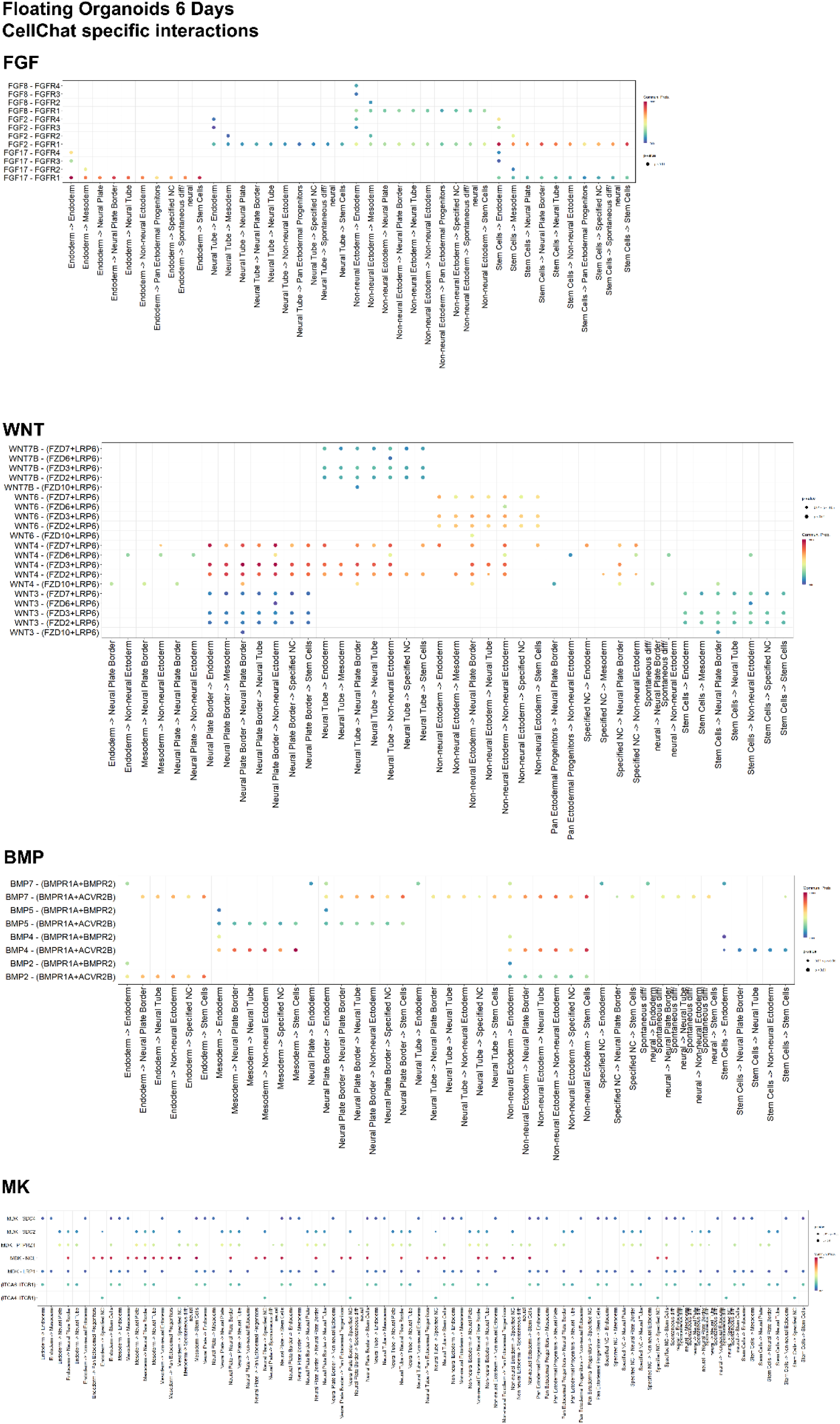
Cell-to-cell signaling of the ectodermal organoids reveal highly conserved mechanisms and novel interaction details. **A)** Detailed ligand-receptor pairs and interaction strengths at day 6 of the floating ectodermal organoids for FGF B) WNT C) BMP and D) Midkine signalling pathways.

**Supp. Fig. 6.**
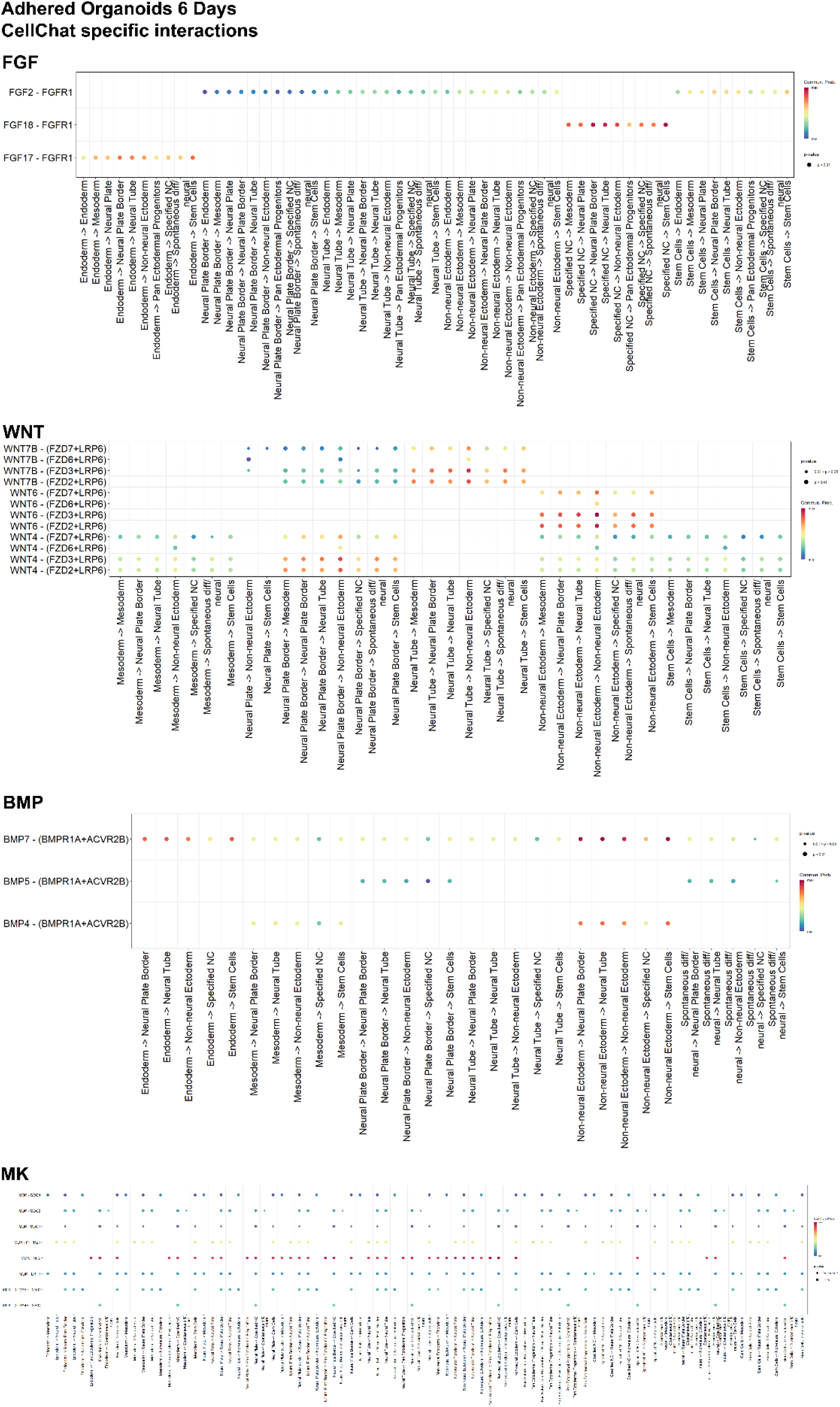
Cell-to-cell signaling of the ectodermal organoids reveal highly conserved mechanisms and novel interaction details. **A)** Detailed ligand-receptor pairs and interaction strengths at day 6 of the adhered ectodermal organoids for FGF B) WNT C) BMP and D) Midkine signalling pathways.

**Supp. Fig. 7.**
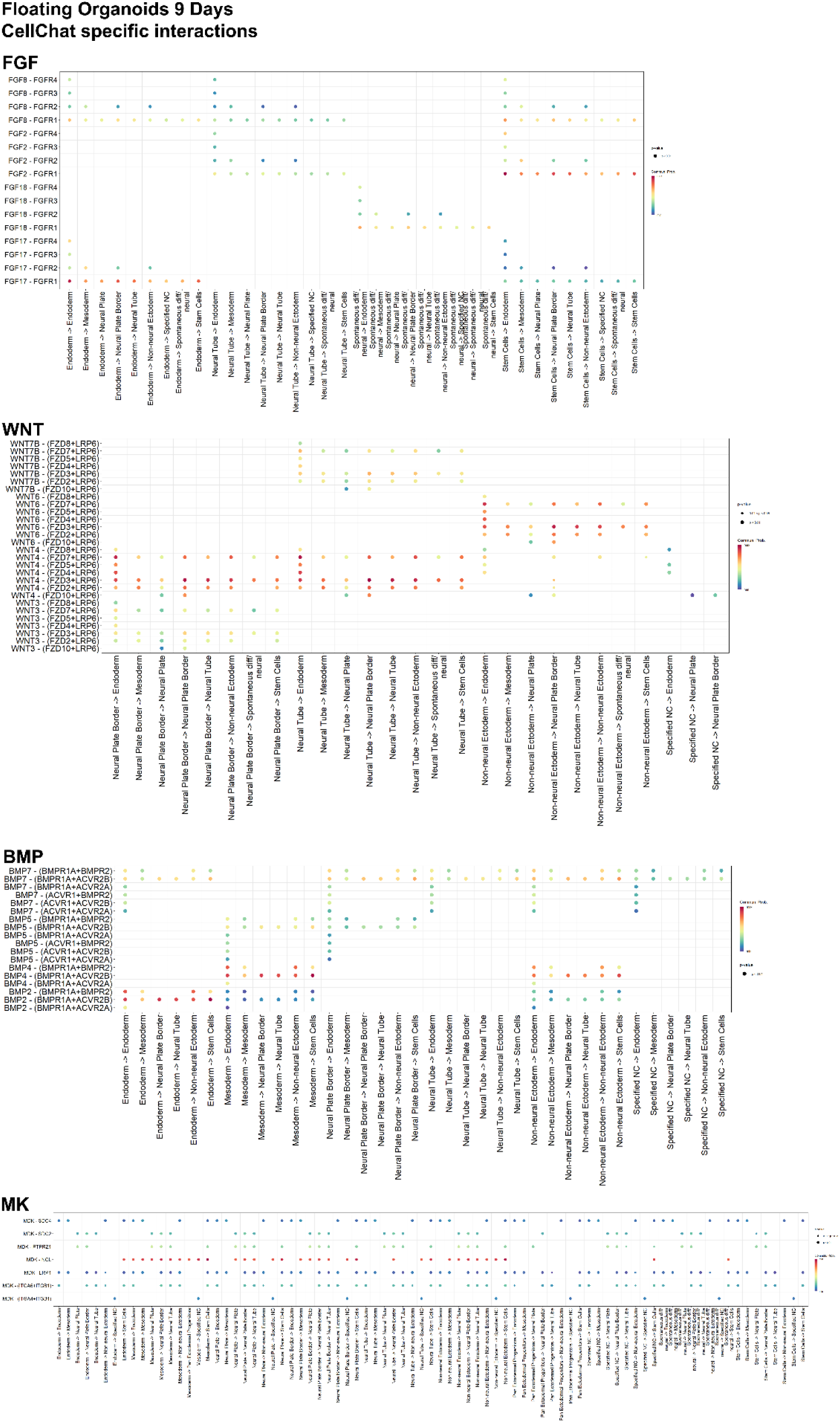
Cell-to-cell signaling of the ectodermal organoids reveal highly conserved mechanisms and novel interaction details. **A)** Detailed ligand-receptor pairs and interaction strengths at day 9 of the floating ectodermal organoids for FGF B) WNT C) BMP and D) Midkine signalling pathways.

**Supp. Fig. 8.**
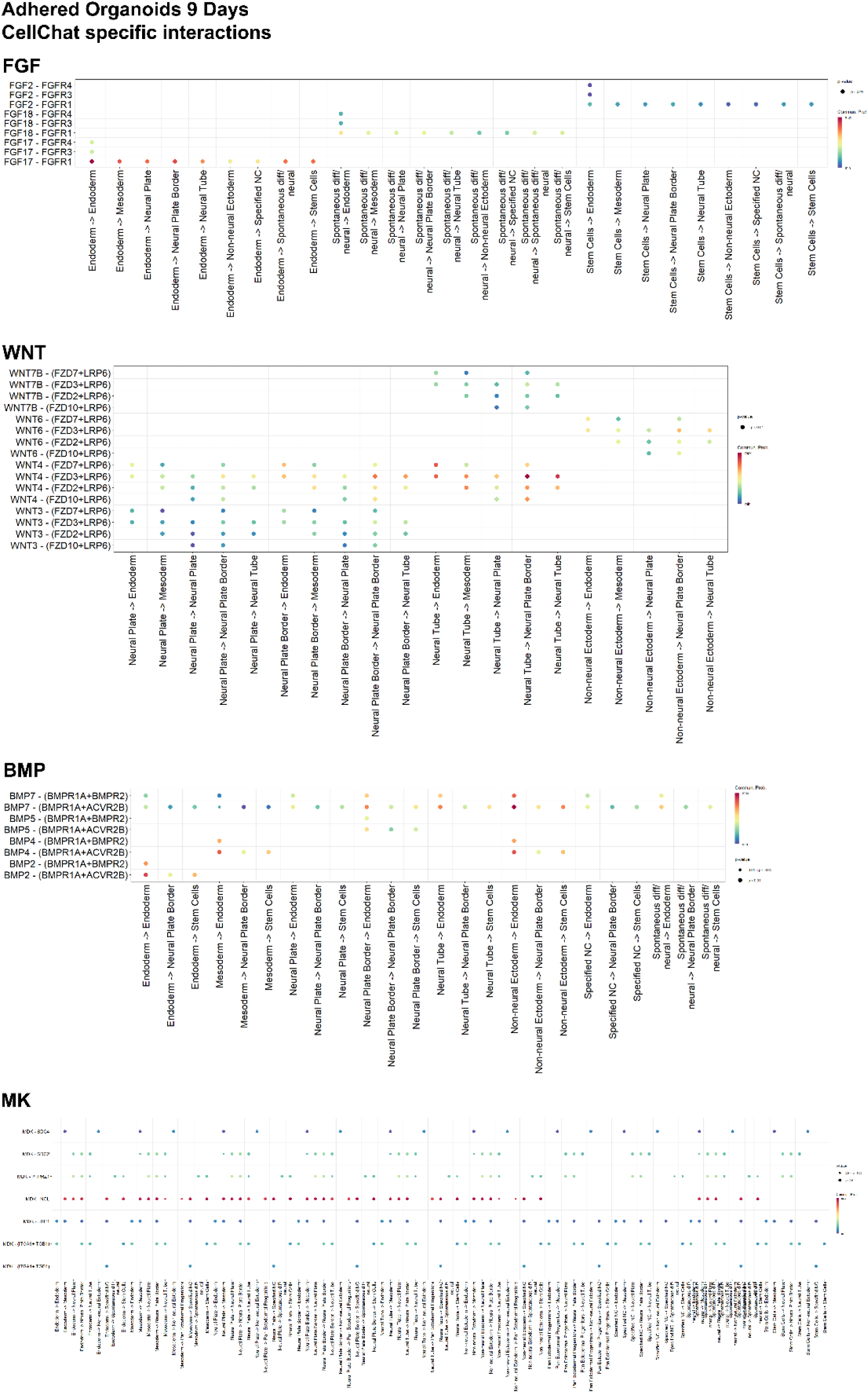
Cell-to-cell signaling of the ectodermal organoids reveal highly conserved mechanisms and novel interaction details. **A)** Detailed ligand-receptor pairs and interaction strengths at day 9 of the adhered ectodermal organoids for FGF B) WNT C) BMP and D) Midkine signalling pathways.

**Supp. Fig. 9.**
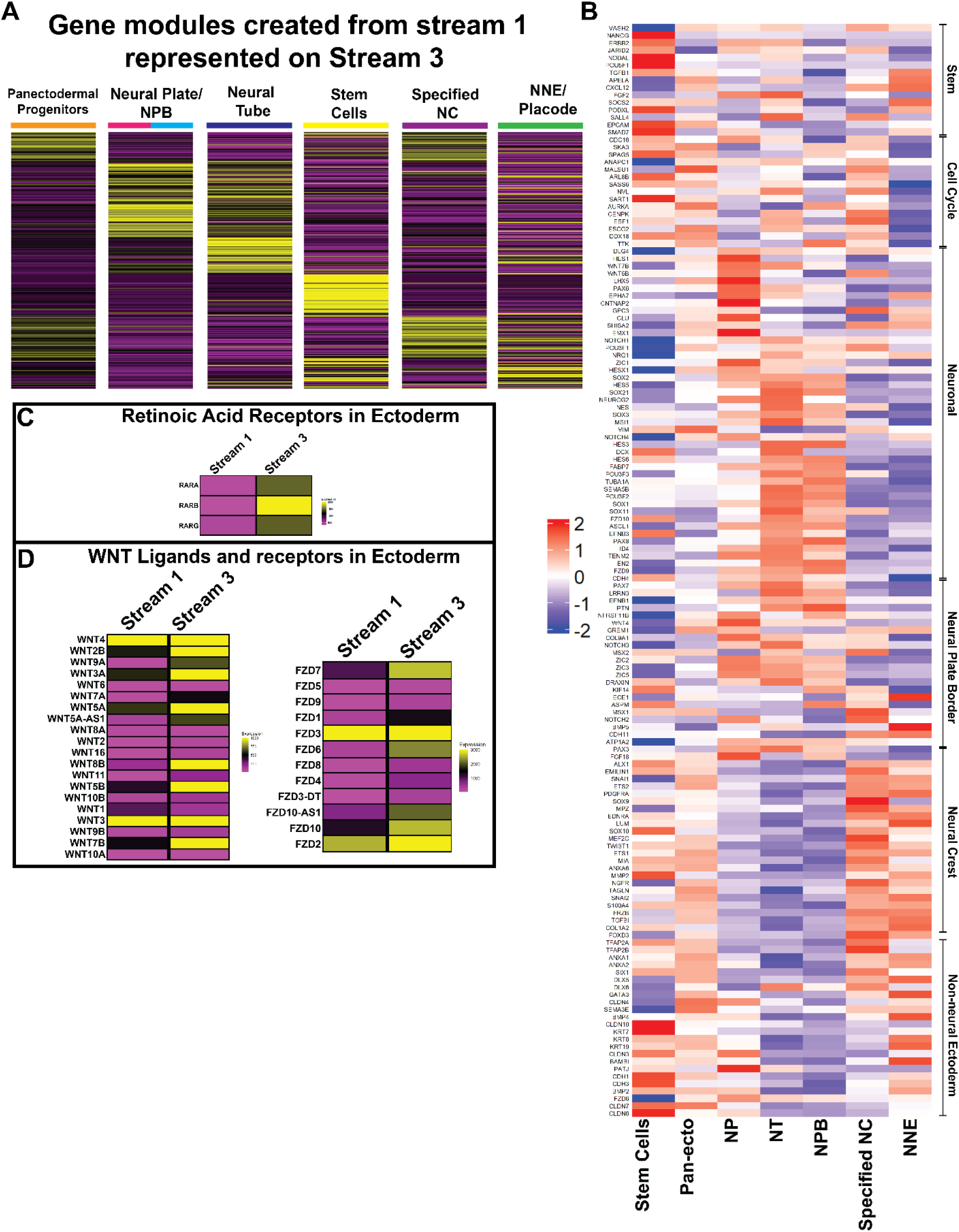
Subpopulations of ectodermal cell types in posteriorized stream 3 of adhered organoids are transcriptionally similar to Stream 1 analyzed by scRNAseq. **A)** Developmental Gene Modules were created from the cranial subpopulations of Day 6 and Day 0 adhered organoids (Stream 1) and the expression of the respective genes in each cell type was plotted on the Stream 3 subpopulations. All modules show high expression in the correct, corresponding cell type. **B**) A heatmap depicting the same list of differentially expressed genes in the same order as in Fig 2H on Stream 3 cell populations shows high similarity between the anterior and posterior subpopulations. **C**) Expression of Retinoic acid receptors increases in Stream 3 as compared to Stream 1. **D**) Expression of WNT ligands and receptors increases in Stream 3 as compared to Stream 1.

**Supp. Fig. 10.**
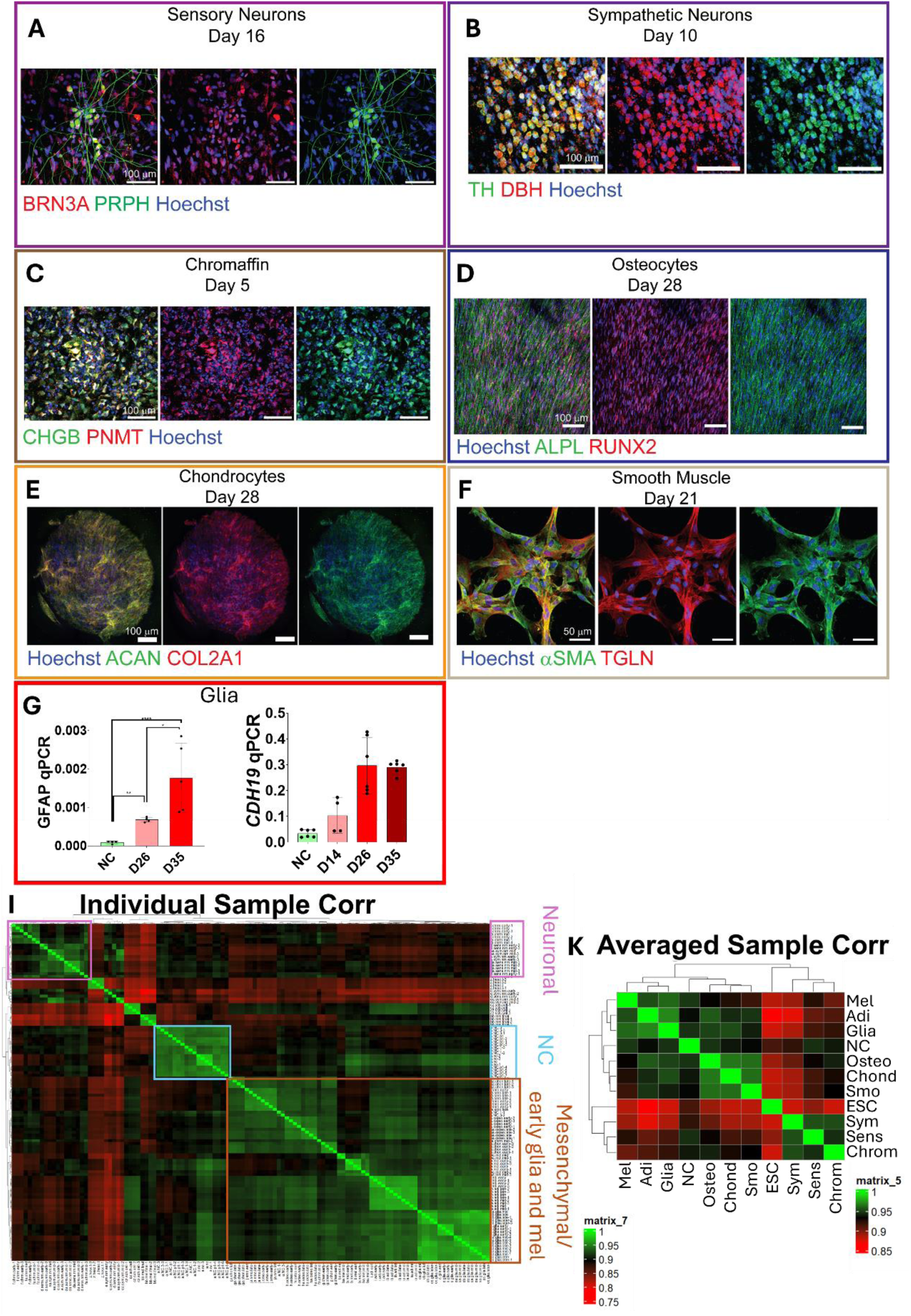
Differentiation of nine diverse neural crest derivative cell types from neural crest cells. **A**) Immunostaining of sensory neurons with POU4F1(BRN3A) and Peripherin. **B**) Immunostaining of sympathetic neurons DBH, and TH shows typical, granular expression. **C**), Immunostaining of Chromaffin cells for Phenylethanolamine N-methylstransferase (PNMT), a crucial enzyme in adrenalin (epinephrin) production cascade as well as for Chomogranin-B (CHGB), a protein that is part of the secretory granules in chromaffin cells (PMID: 13863458; PMID: 6053402), show typical granular expression. **D**) Immunofluorescence with RUNX2 and Alkaline Phosphatase show a synchronized linear organization of osteocytes at day 28. **E**) Chondropheroids express the typical cartilage matrix proteins Aggrecan and Collagen 2A. **F**) Immunostaining for smooth muscle cells actins shows typical expression patterns with anti-Smooth Muscle Actin and - Transgelin (*aka* SM22) antibodies. **G**) Q-PCR shows increased expression of GFAP (NC SD = ±3.408 e-005, D26 SD = ±6.19e-005, D26 SD = ±0.0009) and the schwann cell marker CHD19^216^ (NC SD = ±0.0148, D14 SD = ±0.069, D26 SD = ±0.1092, D35 SD = ±0.02503) during maturation of glial cells. **H**) Correlation plots from bulk RNAseq data show highest correlation between the 1) group of neuronal and chromaffin cells and, on the other hand, 2) between the osteocytes, smooth muscle and chondrocytes as well as the group of early melanocytes, adipocytes and glial cells, which all correlate with each other and show anticorrelation with the neuronal lineage group. **I**) Correlation scores counted from averaged samples from each cell type show a similar pattern as the heatmap of the individual samples.

**Supp. Fig. 11.**
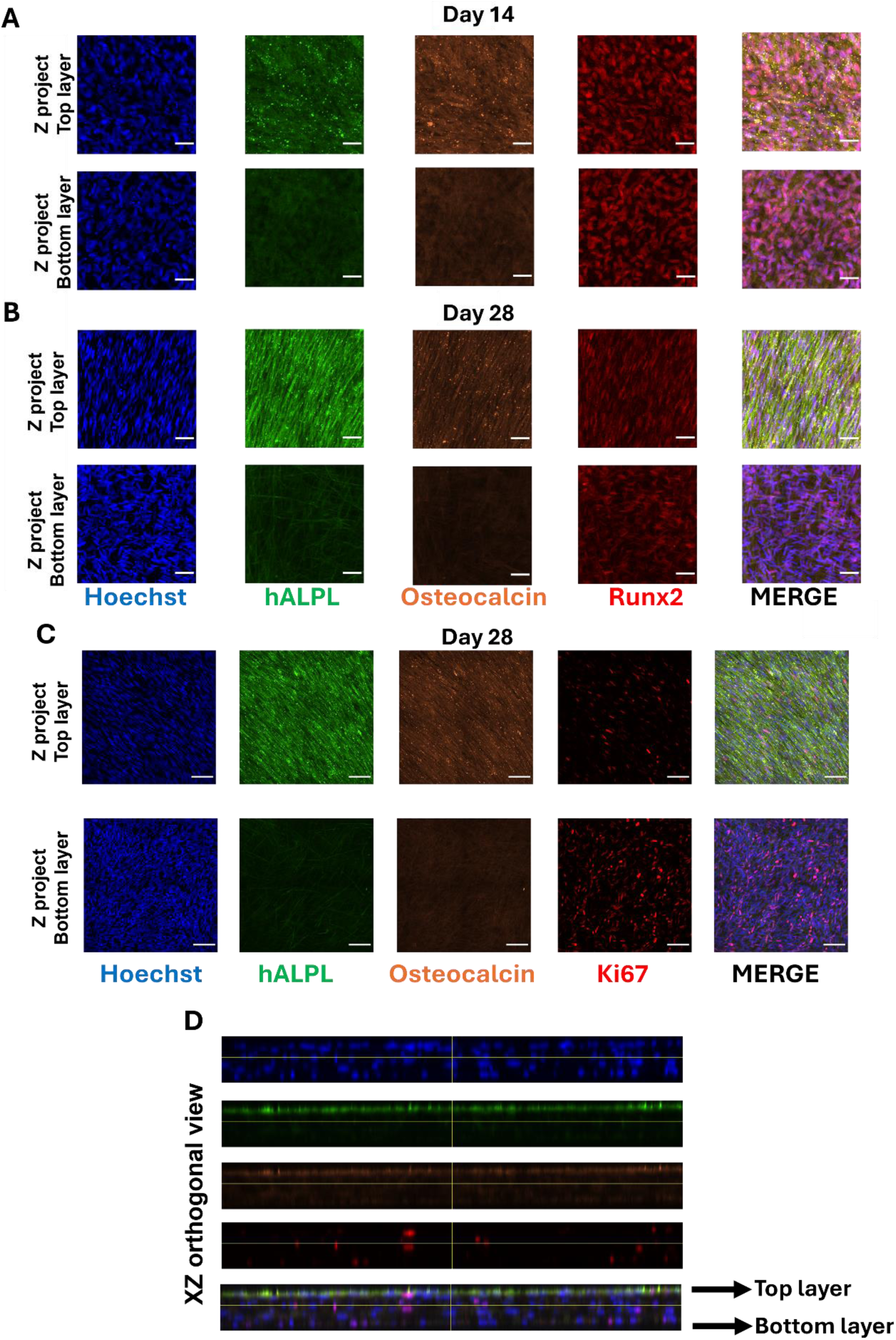
Osteocytes grow as layered structures with an organized mineralizing layer on top of a non-organized, proliferative layer of presumably more progenitor-like osteoblasts. **A)** Immunostaining of mid-point D14 and **B)** endpoint D28 osteoblast cultures with antibodies to lineage driving transcription factor RUNX2 and mineralizing matrix proteins Alkalin Phosphatase and Osteocalcin. Expression of mineralizing proteins starts in the top layer at day 14 but don’t reach a synchronized stripe like structure before the end of the culture. **C**) Immunostaining of endpoint D28 osteoblast cultures with antibodies to the proliferation market Ki67, Alkalin Phosphatase and Osteocalcin at day 28 shows that the bottom layer contains more proliferative cells **D)** XZ orthogonal view of C.

**Supp. Fig. 12.**
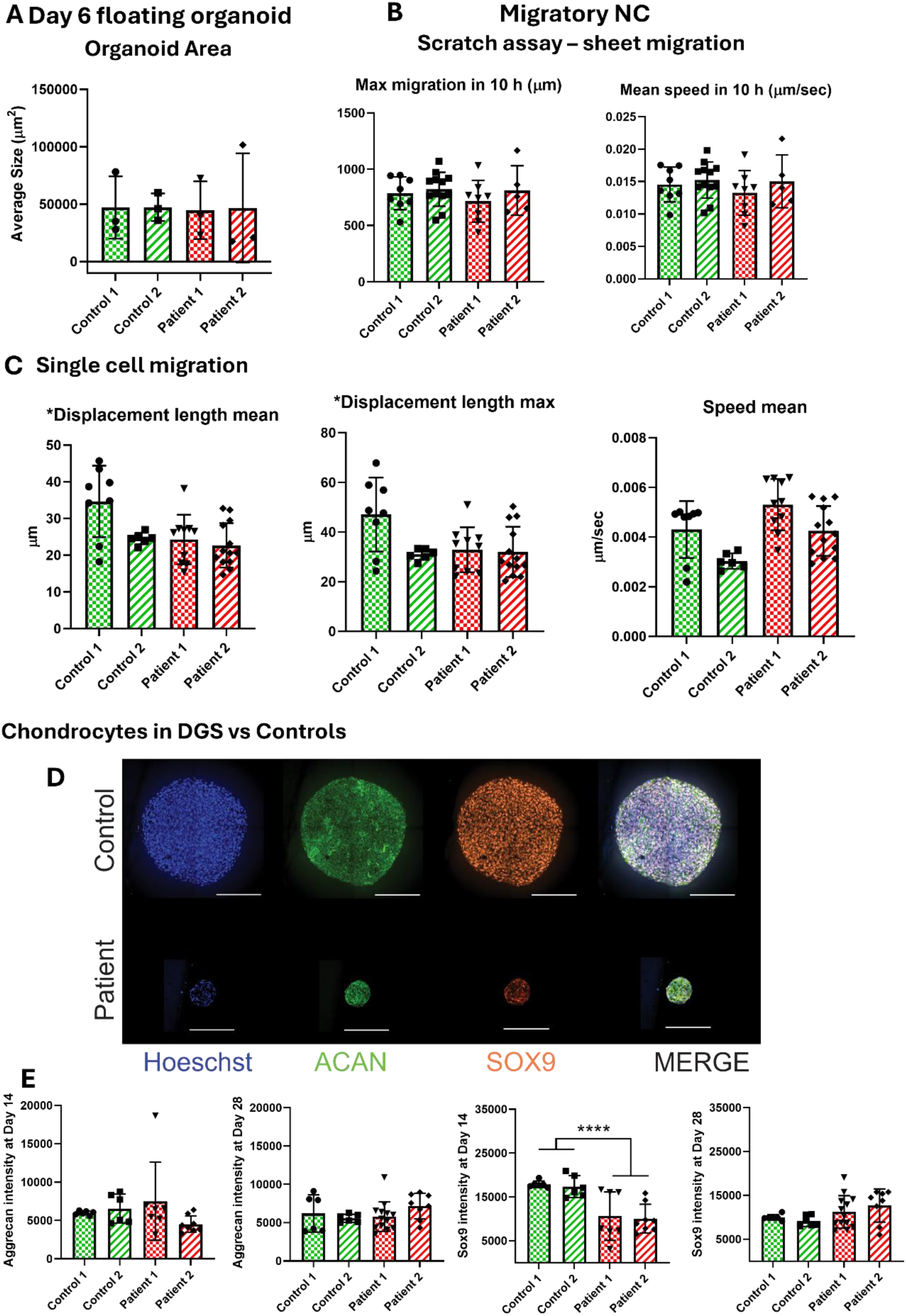
GS NC cells can migrate normally but have defects in chondrogenesis. **A**)Floating ectodermal organoids from DGS patients are similar in size as compared to control organoids. (Control 1 SD = ±27133, Control 2 SD = ±12041, Patient 1 SD = ±25198, Patient 2 SD = ±47587) **B**) Scratch assay shows no change in maximum length and migration speed over 10 hours of culture time of P1 NC cells of DGS vs control cells. (Max Migration: Control 1 SD = ±145.3, Control 2 = ±150.6, Patient 1 SD = ±186.0, Patient 2 = ±220.1. Mean Speed 10h: Control 1 SD = ±0.0027, Control 2 SD = ±0.0028, Patient 1 SD = ±0.0034, Patient 2 SD = ±0.0041) **C**) Measurement of individual migrating P1 NC cells by live imaging show no change in mean or maximum displacement length or in mean speed of the moving cells vetween DGS patient and control cells. (Displacement Mean: Control 1 SD = ±9.725, Control 2 = ±1.581, Patient 1 SD = ±6.737, Patient 2 = ±6.052. Displacement Max: Control 1 SD = ±14.92, Control 2 SD = ±2.259, Patient 1 SD = ±9.057, Patient 2 = ±10.10. Speed Mean: Control 1 SD = ±0.0011, Control 2 SD = ±0.0003, Patient 1 SD = ±0.0010, Patient 2 SD = ±0.0010) D)Immunostaining of DGS and control derived chondrocytes with antibodies to Aggregan and SOX9 on day 28 shows a remarkable size difference of chondrospheroids. **E**) Quantification of immunostaining intensity (normalized to nuclear Hoecst insentity) of Aggregan shows no change in DGS patient chondrospheroids as compared to controls whereas, SOX9 intensity is reduced in DGS patient chondrospheroids at 14 day midpoint but not in the 28 day endpoint of the differentiation cultures. (ACAN intensity D14: Control 1 SD = ±268.9, Control 2 SD = ±1901, Patient 1 SD = ±5086, Patient 2 = ±1081. ACAN intensity D28: Control 1 SD = ±268.9, Control 2 SD = ±1901, Patient 1 SD = ±5086, Patient 2 = ±1081. SOX9 intensity D14: Control 1 SD = ±772.9, Control 2 SD = ±2514, Patient 1 SD = ±5520, Patient 2 = ±3300. SOX9 intensity D28: Control 1 SD = ±680, Control 2 SD = ±1234, Patient 1 SD = ±3722, Patient 2 SD = ±3782).

**Supp. Fig. 13.**
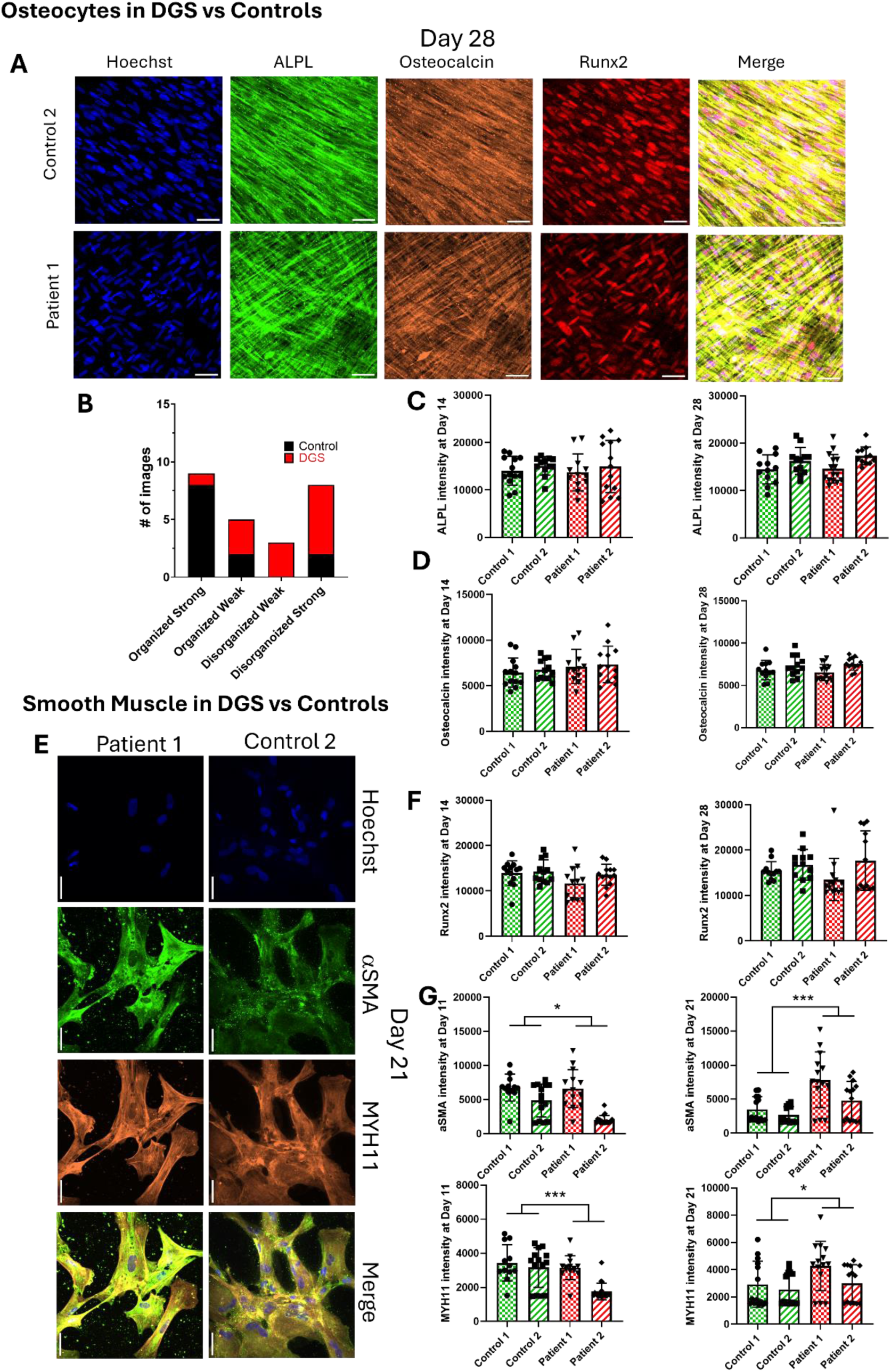
DGS NC cells show defects in osteoblast and smooth muscle cell differentiation. **A**) Osteocytes immunostained with Alkalin Phosphatase, Osteocalcin and RUNX2 and Hoescst show disorganized nuclei and mineralizing proteins in DGS patient cells as compared to controls. **B**) Quantified how many images in control and DGS appeared to be disorganized and ranked based on the strength of certainty of phenotype. **C)** Quantification of intensity for immunostaining of Alkalin Phosphatase shows no difference in osteocytes between control and DGS patient cells at day 14 and 28 of culture. (ALPL intensity D14: Control 1 SD = ±3029, Control 2 SD = ±1970, Patient 1 SD = ±3899, Patient 2 SD = ±5517. ALPL intensity D28: Control 1 SD = ±3063, Control 2 SD = ±2836, Patient 1 SD = ±3015, Patient 2 SD = ±1861) **D**) Quantification of intensity of immunostaining for Osteocalcin shows no difference in osteocytes between control and DGS patient cells at day 14 and 28 of culture. (OST intensity D14: Control 1 SD = ±1609, Control 2 SD = ±1127, Patient 1 SD = ±1866, Patient 2 SD = ±2006. OST intensity D28: Control 1 SD = ±1148, Control 2 SD = ±1232, Patient 1 SD = ±952.4, Patient 2 SD = ±755.2) E) Gross morphological analysis of smooth muscle cells reveals no difference between DGS patient cells and controls immunostained with antibodies to Smooth Muscle Actin and Myosin 11 F) Quantification of intensity of immunostaining for RUNX2 shows no difference in osteocytes between control and DGS patient cells at day 14 and 28 of culture. (RUNX2 intensity D14: Control 1 SD = ±2733, Control 2 SD = ±2634, Patient 1 SD = ±3677, Patient 2 SD = ±2327. RUNX2 intensity D28: Control 1 SD = ±2073, Control 2 SD = ±3378, Patient 1 SD = ±4669, Patient 2 SD = ±6565). G) Quantification of immunostaining for Smooth Muscle Actin and Myosin 11 shows increased expression of both in DGS patient derived cells at day 21 (endpoint) of the differentiation cultures. (aSMA intensity D11: Control 1 SD = ±1985, Control 2 SD = ±2417, Patient 1 SD = ±2797, Patient 2 SD = ±647.2. aSMA intensity S21: Control 1 SD = ±1838, Control 2 SD = ±1226, Patient 1 SD = ±4090, Patient 2 SD = ±2829. MYH11 intensity D11: Control 1 SD = ±1071, Control 2 SD = ±1163 Patient 1 SD = ±698.2, Patient 2 SD = ±483.9. MYH11 intensity D21: Control 1 SD = ±1702, Control 2 SD = ±1202, Patient 1 SD = ±1809, Patient 2 SD = ±1352)

**Supp. Fig. 14.**
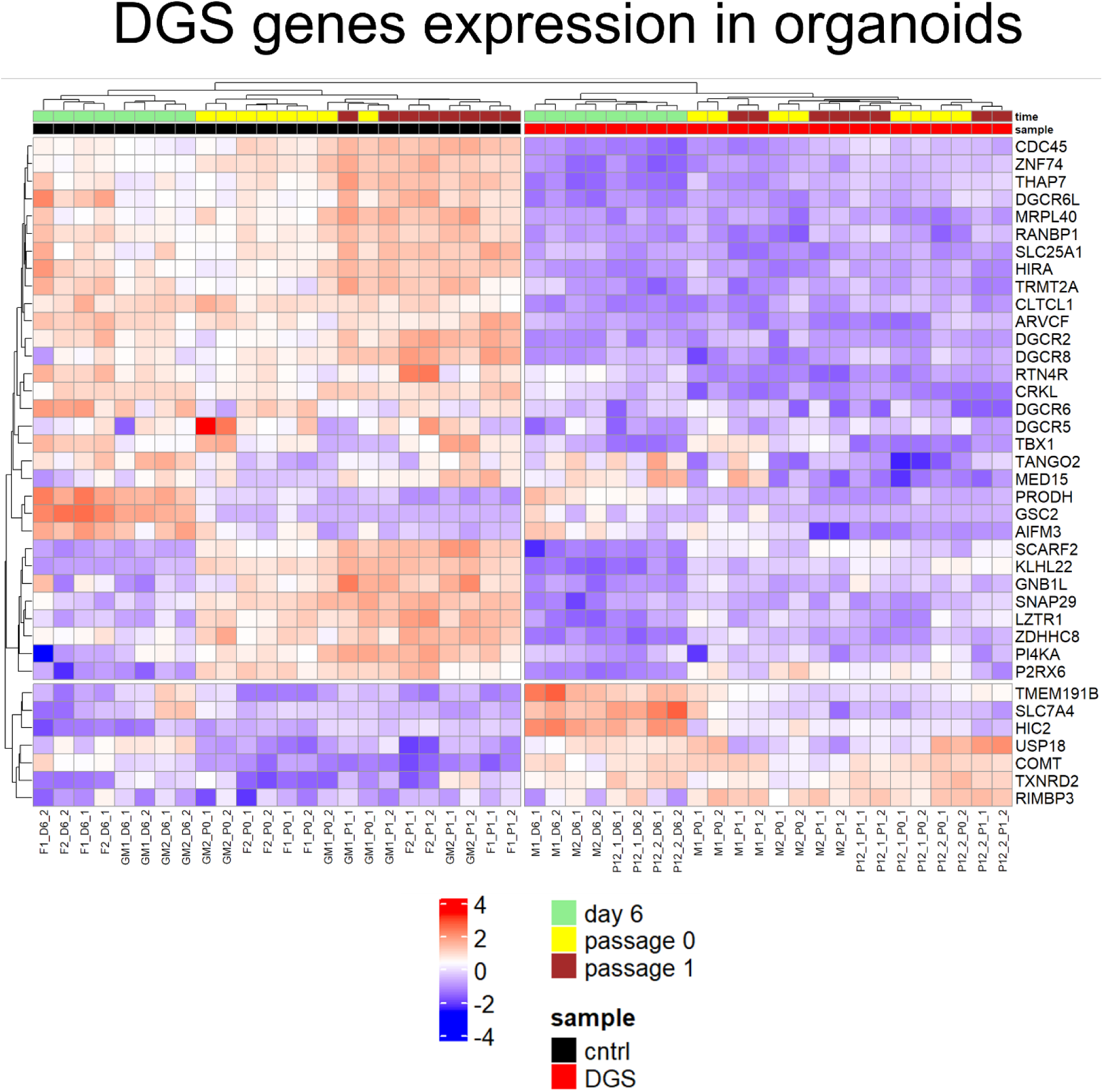
Hemizygosity of DGS genes included in the microdeleted area is reflected in bulk RNAseq samples. As expected, the GDS region genes that are detected by the RNAseq analysis, alhough most of them in biologically irrelevant low levels, are expressed more in ectodermal organoids and migratory NC of control cells. Interestingly, a few genes like USP18 and COMPT show higher expression in DGS cells potentially reflecting a compensation mechanism, although the overall expression. Levels of these genes are extremely low.

**Supp. Fig. 15.**
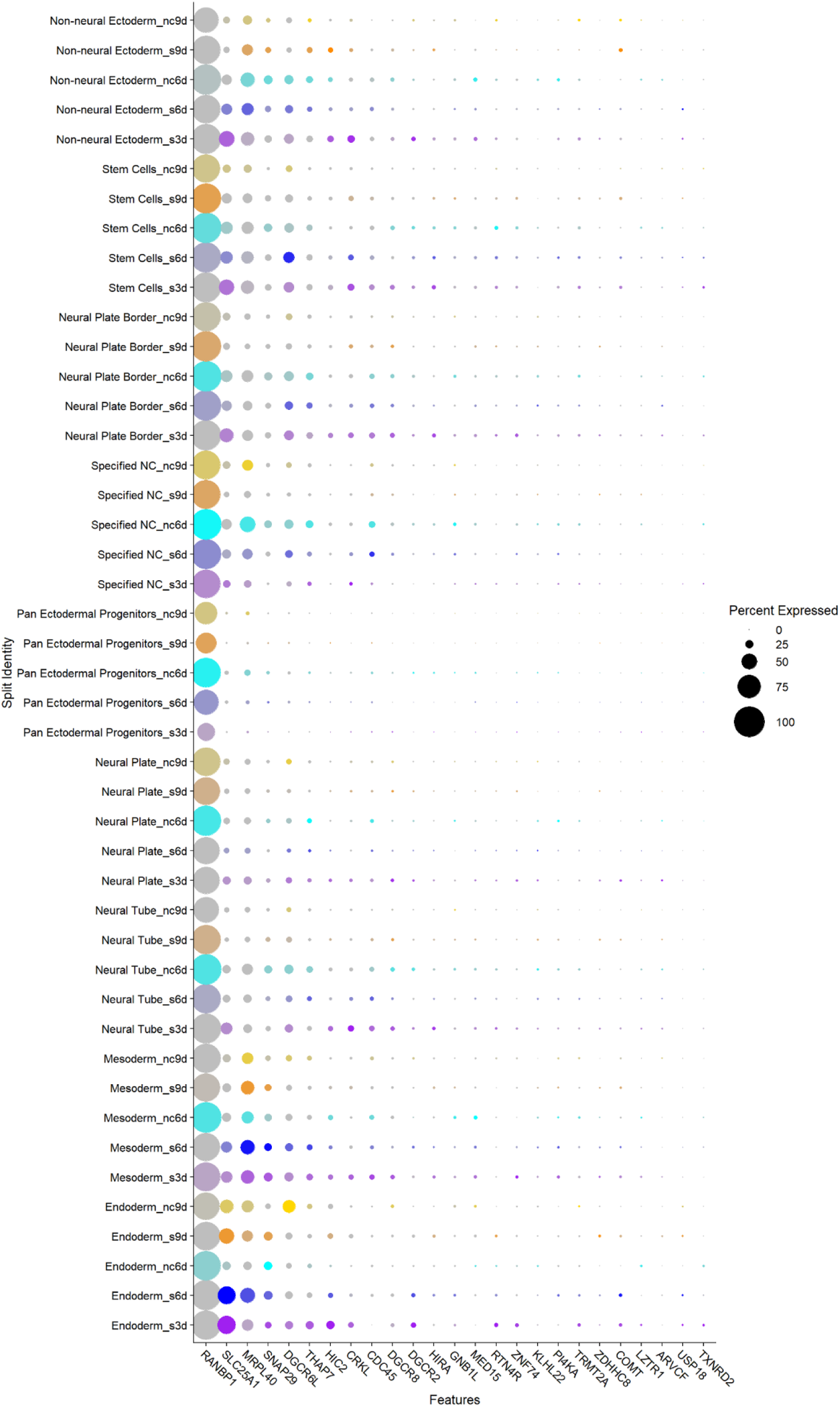
Bubble plot demonstrates relative expression genes that are located in the 3MB DGS microdeleted region in all subpopulations of the ectodermal organoids. Note that only the genes with detectable expression in the scRNAseq data set are plotted.

**Supp. Fig. 16.**
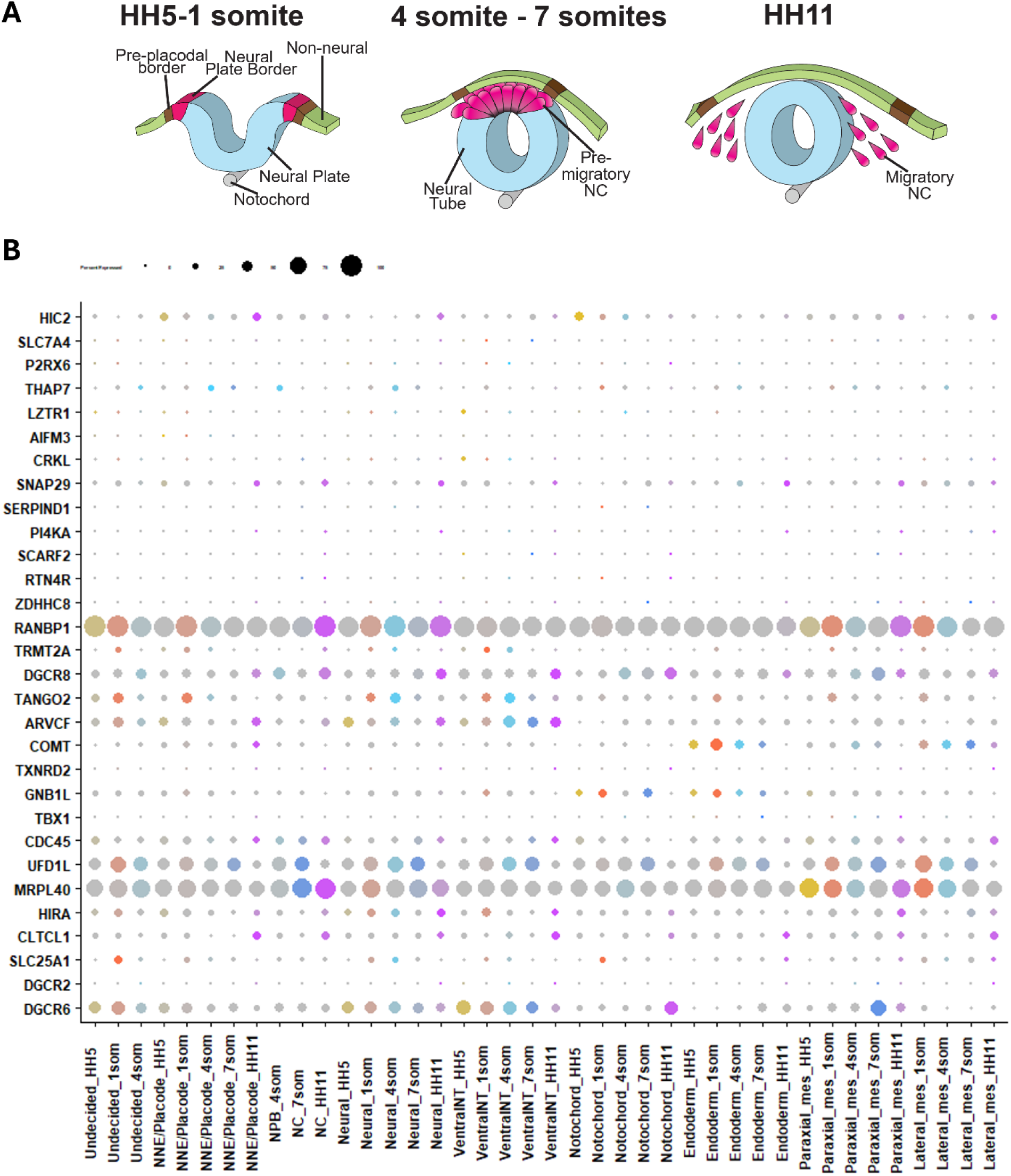
Bubble plot demonstrates relative expression genes that are located in the DGS microdeleted region in all subpopulations of the cranial chicken embryo. Note that only the genes with detectable expression in the scRNAseq data set are plotted.

**Supp. Fig. 17.**
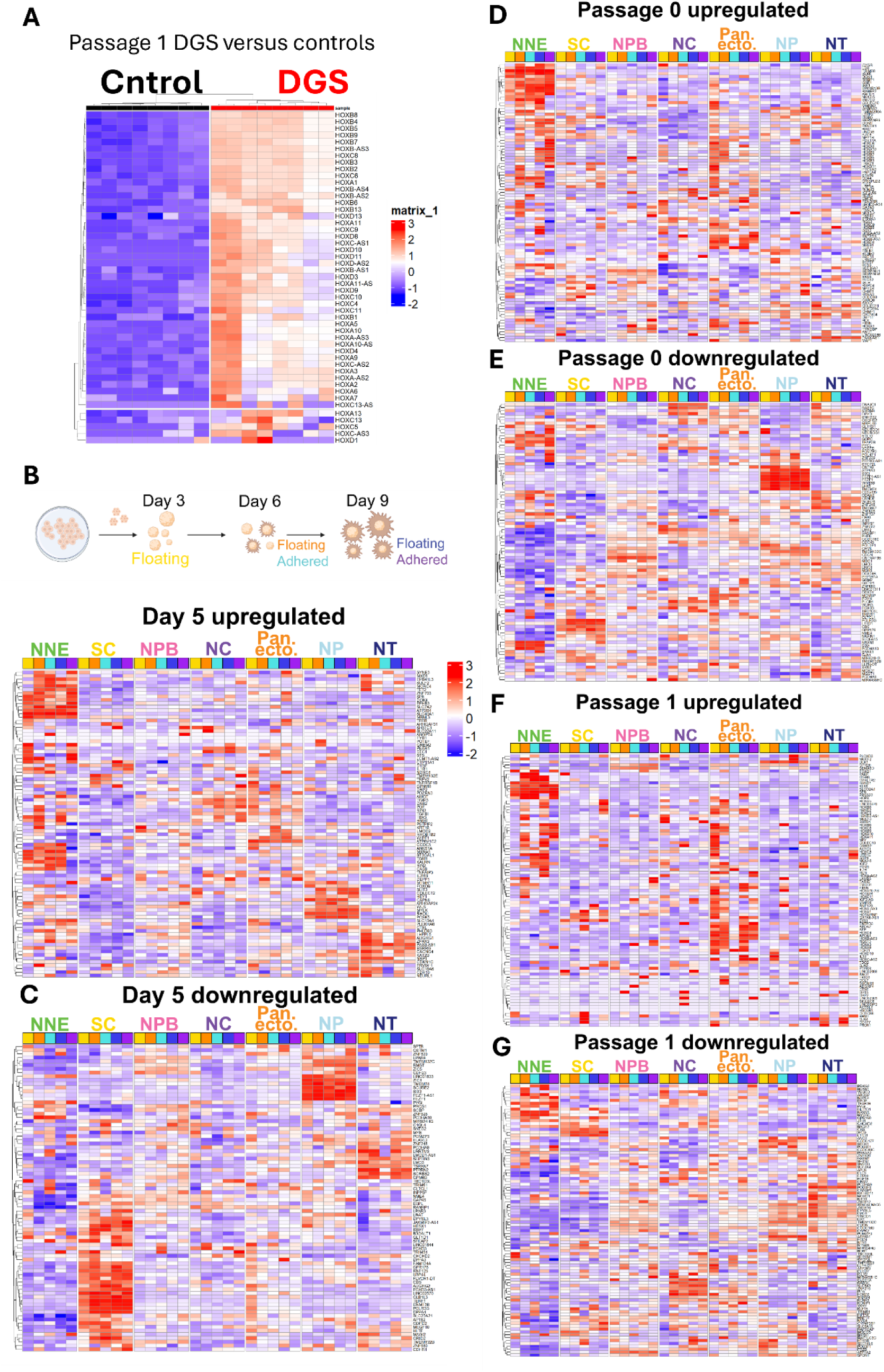
Uppregulated genes of the DGS consist of committed lineage genes, and downregulated genes are normally expressed by the stem cells and early stages of ectodermal patterning. **A)** Heatmap representation showing that HOX gene expression in the Passage 1 NC of DGS patients. **B)** The heatmap shows expression of all top 200 differentially expressed up and **C)** downregulated genes, respectively, of the floating day 5 ectodermal organoids derived from DGS patient cells as compared to controls. The genes from the DGS bulk RNA data set are plotted on the scRNAseq data set of WT cells from the different subpopulations of ectodermal organoids at day 6. The results show a similar pattern as in Figure 8G. **D**) The heatmap shows expression of all top 200 differentially expressed up and **E)** downregulated genes, respectively, of the P0 primary migratory NC cells derived from DGS patient cells as compared to controls. The genes from the DGS bulk RNA data set are plotted on the scRNAseq data set of all WT cells from the different subpopulations of ectodermal organoids at days 3-9. The results show a similar pattern as in Figure 8H. **F**) The heatmap shows expression of all top 200 differentially expressed up and **G)** downregulated genes, respectively, P1 migratory NC cells derived from DGS patient cells as compared to controls. The genes from the DGS bulk RNA data set are plotted on the scRNAseq data set of all WT cells from the different subpopulations of ectodermal organoids at days 3-9. The results show a similar pattern as in Figure 8I. DGS: DiGeorge Syndrome Patient, NNE: Non-Neural Ectoderm, SC: Stem Cells, NPB: Neural Plate Border, NC: Specified Neural Crest, Pan-ecto: Pan-ectodermal Progenitors, NP: Neural Plate, NT: Neural Tube.

